# Integrated Bioinformatics Analysis Deciphering the microRNA Regulation in Protein-Protein Interaction Network in Lung Adenocarcinoma

**DOI:** 10.1101/2020.08.01.232306

**Authors:** Pratyay Sengupta, Sayoni Saha, Moumita Maji, Monidipa Ghosh

## Abstract

**Background:** The architecture of the protein-protein interaction (PPI) network in any organism relies on their gene expression signature. microRNAs (miRNAs) have recently emerged as major post transcriptional regulators that control PPI by targeting mainly untranslated regions of the gene encoding proteins. Here, we aimed to unveil the role of miRNAs in the PPI network for identifying potential molecular targets for lung adenocarcinoma (LUAD).

**Materials and methods:** The expression profiles of miRNAs and mRNAs were collected from the NCBI Gene Expression Omnibus (GEO) database (GSE74190 and GSE116959). Abnormally expressed mRNAs from the data were appointed to construct a PPI network and hence incorporated with the miRNA-mRNA regulatory network. The miRNAs and mRNAs in this network were subjected to functional enrichment. Through the network analysis, hubs were identified and their mutation rate and probability of cooccurrence were calculated.

**Results:** We identified 17 miRNAs and 429 mRNAs signature for differentially altered transcriptome in LUAD. The combined miRNA–mRNA regulatory network exhibited scale-free characteristics. Network analysis showed 5 miRNA (including hsa-miR-486-5p, hsa-miR-200b-5p, and hsa-miR-130b-5p) and 10 mRNA (including ASPM, CCNB1, TTN, TPX2, and BIRC5) which expressively contribute in the LUAD. We further investigated the hub genes and noticed that ASPM and TTN had the maximum rate of mutation and possessed a high tendency of cooccurrence in LUAD.

**Conclusion:** This study provides a unique network approach to the exploration of the underlying molecular mechanism in LUAD. Identified mRNAs and miRNAs may therefore, serve as significant prognostic predictors and therapeutic targets.

## 1. Introduction

Lung adenocarcinoma (LUAD), the most common primary lung cancer, mainly manifests in the form of non-small cell lung cancer (NSCLC) (Myers and Wallen 2020). As with other tumors, lung adenocarcinoma shows heterogeneity by high rates of genetic mutation (Dong et al. 2020). Despite the availability of diverse technologies, such as stereotactic radiotherapy, targeted therapy, minimal invasion methods and immunotherapy, the long-term survival is still poor in LUAD (Ferrara et al. 2018). One of the main reasons for this could be a delayed diagnosis. Thus, it is necessary to understand the molecular mechanisms behind lung adenocarcinoma genesis and metastasis, and identify effective biomarkers of this disease. MicroRNAs (miRNAs) are the 19-25 nucleotides long single-stranded non-coding RNAs, that regulate genes by mainly binding to the 3’ untranslated regions (UTRs) of their target genes (Berindan-Neagoe et al. 2014). In recent, miRNAs were demonstrated to be significantly dysregulated in LUAD tissues and could be a potential target for this disease (Petkova et al. 2022; Han and Li 2018).

In the post-genomic era, biological networks offer exciting possibilities in understanding gene regulation and pattern recognition for autonomous disease identification. Protein-protein interaction (PPI) network is conventionally a static network, where proteins are represented by nodes and interactions by edges (Barabasi and Oltvai 2004). However, in reality, PPI network has dynamic property, which is intrinsically regulated by complex cellular mechanisms through time and space (Liang and Li 2007). Recent studies showed the significance of the miRNA regulation on protein-protein interaction networks (Rahimpour et al. 2022; Du et al. 2019; Lei et al. 2019; Liang and Li 2007). Moreover, different methods have been developed to predict miRNA targets accurately at the genomic level (John et al. 2004; Lewis et al. 2003). But much effort had not been given to understand the relationship between miRNA regulations on PPI in LUAD from a regulatory network approach.

Gene Expression Omnibus (GEO) is a public data repository of array- and sequence-based functional genomics data (Edgar et al. 2002). Owing to the advancement of high-throughput bioinformatics technologies, we aimed to re-analyze the mRNA (GSE116959) and miRNA (GSE74190) expression profile data sets in LUAD from GEO to unveil more specific molecular targets associated with the tumorigenesis of the disease. We sought to construct a regulatory network combining miRNA-mRNA and protein-protein interactions in LUAD. Our study presented an integrative bioinformatics data analysis that could be appointed for the improvement in the clinical consequences aided through endorsement of the potential biomarkers in LUAD.

## 2. Materials and Methods

### 2.1 Data

The transcriptomic data used in this publication were collected from the National Center of Biotechnology Information (NCBI) Gene Expression Omnibus (GEO) (https://www.ncbi.nlm.nih.gov/geo/) (Edgar et al. 2002). This gene expression profile was obtained from the GSE116959 dataset (Moreno Leon et al. 2019). The platform used for this data is the GPL17077 Agilent-039494 SurePrint G3 Human GE v2 8×60K Microarray 039381 (Probe Name version). The GSE74190 dataset contains microRNA expression profiles of LUAD patients. GPL19622 Agilent-019118 Human miRNA Microarray V1 G4470A [miRBase release 9.1 miRNA ID version] platform used for the miRNA profile data.

### 2.2 Screening of Dysregulated miRNA and mRNA

In our study, GEO2R (http://www.ncbi.nlm.nih.gov/geo/geo2r/), an R based interactive web tool, was used to identify differentially expressed miRNAs and mRNAs between tumor and adjacent normal samples separately within each dataset (Barrett et al. 2013). A log fold change (logFC) >2 was set as a cut off for differentially expressed miRNAs and mRNAs. Probes without a corresponding gene symbol were filtered. The data were processed through the paired samples t-test and corrected using the Benjamini – Hochberg method. The adjusted P-value <0.05 cut off was applied. The log2 normalized expression values of differentially expressed miRNAs and mRNAs were employed for the complete unsupervised clustering using online tool MORPHEUS (https://software.broadinstitute.org/morpheus/). Further, Principal component analysis (PCA) and volcano plot of the same were performed using the R based online tool ClustVis (https://biit.cs.ut.ee/clustvis/) and the MATLAB v.R2018a respectively (Metsalu and Vilo 2015).

### 2.3 Protein-Protein Interaction Network Construction

The Protein-protein interaction (PPI) network was constructed and analyzed by using the STRING: functional protein association network database (https://string-db.org). The dysregulated mRNA signature in LUAD was set as the input and the highest confidence (0.90) of active interaction was chosen. The isolated nodes were hidden and supervised clustering (K-means) was performed for recognizing three clusters in the provided data set.

### 2.4 Development of miRNA-mRNA Expression Network

Interactions between dysregulated miRNAs and mRNAs were anticipated using miRWalk 3.0 (http://mirwalk.umm.uni-heidelberg.de/) (Sticht et al. 2018), which compiled the prediction results of both TargetScan(Agarwal et al. 2015) and miRDB(Chen and Wang 2020), and a maximum score (= 1) was considered for the prediction analysis in miRWalk. The miRNA-mRNA interaction network was incorporated with the PPI network for a better understanding of the underlying mechanism in LUAD. The Cytoscape v.3.8.0 (https://cytoscape.org/) was used for visualizing the regulatory network between miRNA-mRNA. Cytohubba, a cytoscape plug-in, was employed to predict top 5 miRNAs and top 10 mRNAs based on their degree. A network with these 5 miRNAs and their common target mRNAs was constructed and analyzed. BoxPlotR (http://shiny.chemgrid.org/boxplotr/), an online tool was used to create violin plots for the degree centrality (DC), closeness centrality (CC), and betweenness centrality (BC) data of the nodes present in the developed interactive network.

### 2.5 Functional Enrichment Analysis

Functional enrichment analysis was done with the significantly dysregulated miRNAs and mRNAs in LUAD development using FUNRICH tool v.3.1.3 (http://www.funrich.org/) taking the human proteome as background from the Gene Ontology database (http://geneontology.org/) (Pathan et al. 2015; Pathan et al. 2017). Top 5 significant cellular components (CC), molecular functions (MF) and biological process (BP) for miRNAs and mRNAs were plotted.

### 2.6 Pathway Analysis

Pathway analysis was carried out for differentially expressed mRNA signatures in LUAD using the open access, manually curated, peer-reviewed pathway database Reactome (https://reactome.org/). A cut-off of p value < 0.05 was set to recognize the altered signaling pathways.

### 2.7 Screening and Overall Survival Analysis of the Hub Genes

The mutation rates and the tendency of cooccurrence of the top 10 hub genes were examined with the cBioPortal platform (http://www.cbioportal.org). Further, the overall survival plots for the hub miRNAs and mRNAs were plotted using the Kaplan–Meier plotter (https://kmplot.com/analysis/) (Gyorffy et al. 2013) to determine their prognostic characteristics.

## 3. Results

### 3.1 Genetic Alteration in miRNAs and mRNAs Expression in LUAD

Our analysis revealed 17 miRNAs and 429 mRNAs to be differentially expressed in LUAD tissues (**Supplementary Table S1 and S2**). Dysregulated 17 miRNAs from the GSE74190 dataset includes 6 over-expressed and 11 under-expressed miRNAs. Analysis of the GSE116959 dataset showed 122 up-regulated and 307 down-regulated mRNAs. Unsupervised clustering of dysregulated mRNAs and miRNAs exhibited a distinct difference between tumor and normal samples (**Figure. 1A-B**). To identify the expression levels of differentially expressed mRNAs and miRNAs the volcano plots were made (**Figure. 1C-D**). The principal component analysis was plotted by using first and second components with 69.5% variability in miRNA level and 59.4% variability in mRNA level, represents the percentage of variation in their respective levels (**Figure. 1E-F**). This results further substantiates the clustering pattern of the dysregulated mRNA and miRNA signatures. The particulars of the data dimension employed in this report are provided in **Table 1**.

**Table 1.** Details of the data sets of lung adenocarcinoma downloaded from the NCBI GEO data portal.

| GSE Accession Numbers | RNA type | Tumor Sample Count | Adjacent Normal Sample Count | Identified RNA |
| --- | --- | --- | --- | --- |
| GSE116959 | messenger RNA | 57 | 11 | 33985 |
| GSE74190 | micro RNA | 36 | 44 | 821 |

**Figure 1:**
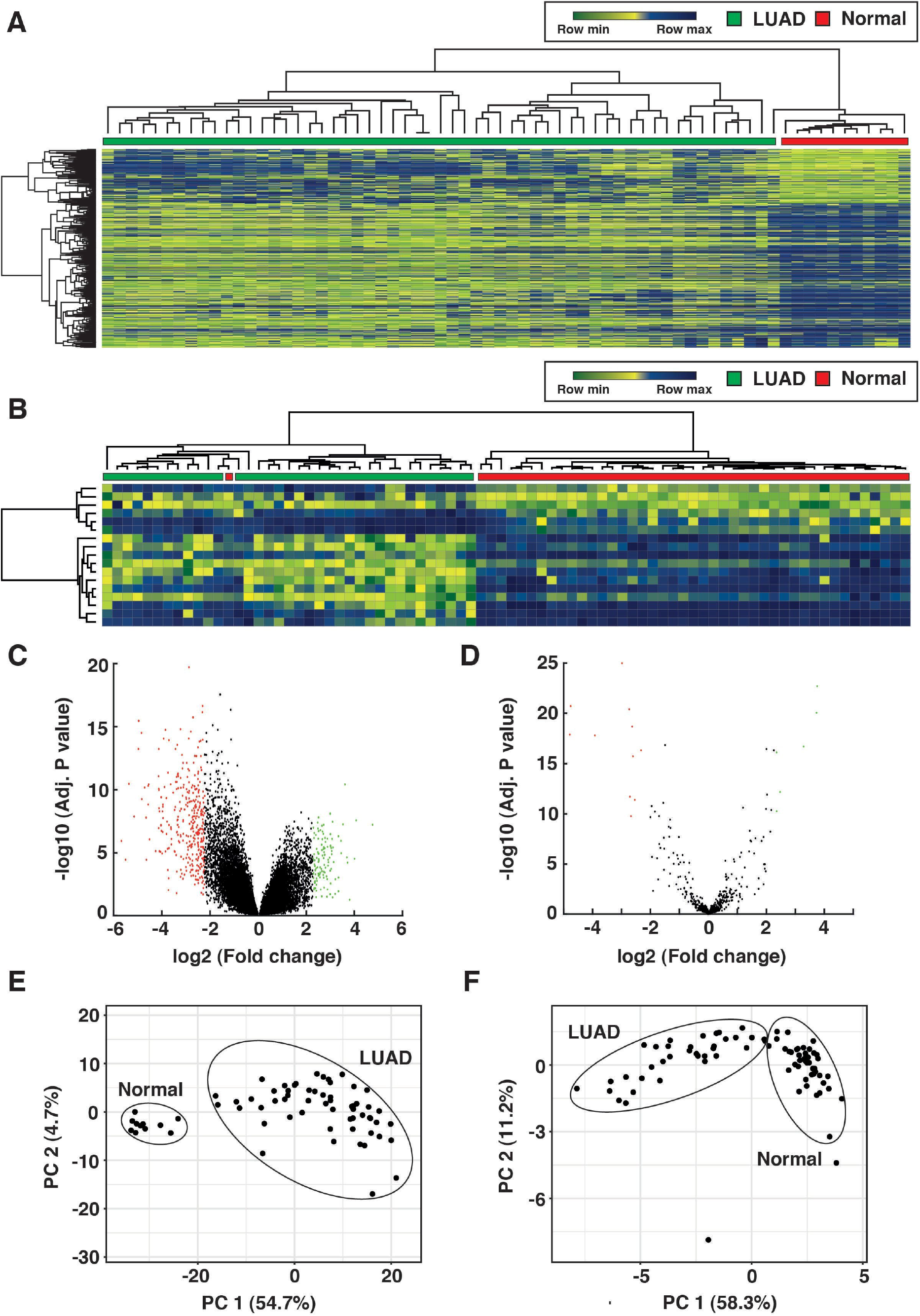
Differential expression pattern analysis of miRNA and mRNA obtained from lung adenocarcinoma tumor and normal samples. **A-B**. Unsupervised clustering of dysregulated mRNAs (A) and miRNAs (B) along with their expression pattern in LUAD is illustrated in the heatmap developed using MORPHEUS (https://software.broadinstitute.org/morpheus/). **C-D**. Relative expression of differentially expressed 465 mRNAs (C) and 17 miRNAs (D) can be observed in the Volcano Plot. These plots were made by using MATLAB v.R2018a. Green, Red and Black dots on the plot display upregulated, downregulated and dysregulated expression respectively. **E-F**. Principal component analysis (PCA) of the dysregulated mRNAs (E) and miRNAs (F) was performed by ClustVis (https://biit.cs.ut.ee/clustvis/) showing their corresponding variations in the clusters of tumor and normal samples.

### 3.2 Interaction of Dysregulated Proteins in LUAD

The 429 differentially expressed mRNAs were assigned for the PPI network formation. The network exposed two major clusters with GNG11 (Guanine nucleotide-binding protein subunit gamma) and CCNB2 (G2/mitotic-specific cyclin-B2).

GNG11 was identified to be a key hub protein that interacted with 15 other proteins including GRK5 (G protein-coupled receptor kinase 5), CALCRL (Calcitonin gene-related peptide type 1 receptor), VIPR1 (Vasoactive intestinal polypeptide receptor 1), NMUR1 (Neuromedin-U receptor 1), ADRB1 (Beta-1 adrenergic receptor), SSTR1 (Beta-1 adrenergic receptor), ADRB2 (Beta-2 adrenergic receptor), CX3CR1 (CX3C chemokine receptor 1), EDN1 (Endothelin-1), CHRM1 (Muscarinic acetylcholine receptor M1), NMU (Muscarinic acetylcholine receptor M1), CACNA2D2 (Voltage-dependent calcium channel subunit alpha-2/delta-2), CXCL13 (C-X-C motif chemokine 13), GPER1 (C-X-C motif chemokine 13), and RAMP3 (Receptor activity-modifying protein 3) (**Supplementary Figure S1A**). CCNB2, another key hub protein interacted with KIF2C (Kinesin-like protein KIF2C), BIRC5 (Baculoviral IAP repeat-containing protein 5), CENPF (Centromere protein F), UBE2C (Ubiquitin-conjugating enzyme E2 C), TPX2 (Targeting protein for Xklp2), SPAG5 (Sperm-associated antigen 5), ASPM (Abnormal spindle-like microcephaly-associated protein), KIAA0101 (PCNA Clamp Associated Factor), CDC45 (Cell division control protein 45 homolog),CDT1 (DNA replication factor Cdt1),HJURP (Holliday junctionrecognitionprotein), CEP55 (Centrosomal protein of 55 kDa), CCNB1 (G2/mitotic-specific cyclin-B1), PKMYT1 (Membrane-associated tyrosine- and threonine-specific cdc2-inhibitorykinase), and CDCA5 (Sororin) (**Supplementary Figure S1B**). The details of the PPI networkwere provided in **Supplementary Table S3**.

### 3.3 miRNA-mRNA Interacting Network Analysis

The role of miRNA on PPI network in LUAD was investigated by constructing a miRNA-mRNA regulatory network and visualized using Cytoscape (**Figure. 2A**). Interaction analysis showed that 17 dysregulated miRNAs aimed 321 dysregulated mRNAs for up-regulating or down-regulating them. In details, 6 over-expressed miRNAs down-regulated 178 mRNAs, whereas 11 under-expressed miRNAs up-regulated 69 mRNAs. Eventually, we incorporate the PPI interactions which include 287 interactions. In the combined interaction network, total number of nodes is 363 with the 1346 connections (**Supplementary Table S4**).

**Figure 2:**
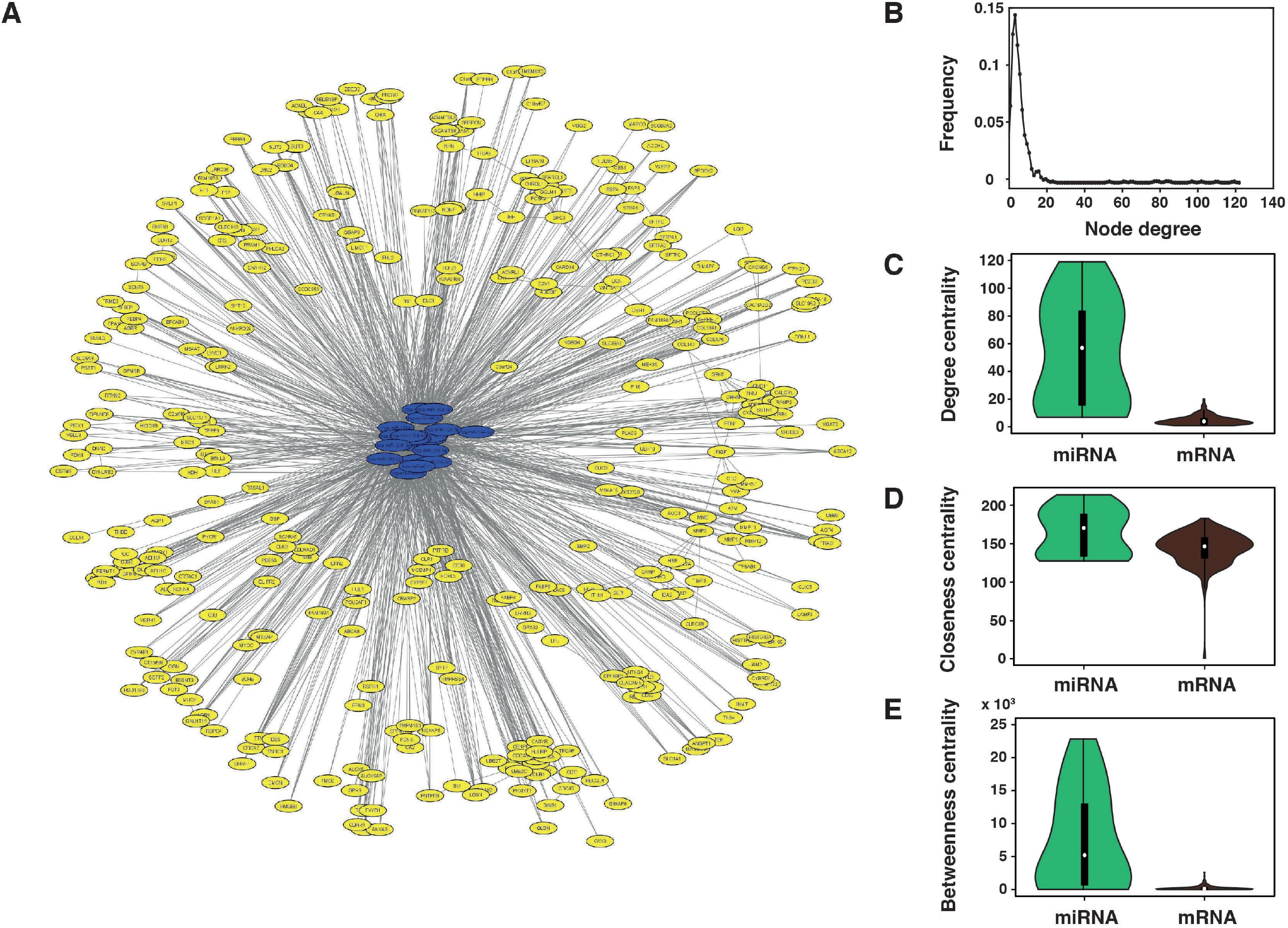
miRNA-mRNA regulatory network construction and characterization. **A**. This regulatory network analysis was visualized in Cytoscape v.3.8.0 (https://cytoscape.org/) showing that 17 dysregulated miRNAs targeting 321 dysregulated mRNAs. Yellow nodes in the regulatory network represent the mRNAs and blue nodes represent the miRNAs.**B**. Degree distribution of the regulatory network. **C**. The difference of degree centrality between miRNAs and mRNAs. The miRNA nodes showed an aberrantly higher degree centrality than mRNA nodes. **D**. The difference of closeness centrality between miRNAs and mRNAs. The miRNA nodes showed a significantly higher closeness centrality than mRNA nodes. **E**. The difference of betweenness centrality between miRNAs and mRNAs. The miRNA nodes had a significantly higher betweenness centrality than mRNAs. P-values were calculated based on Mann – Whitney U test.BoxPlotR (http://shiny.chemgrid.org/boxplotr/) was used to create violin plots for the degree centrality (DC), closeness centrality (CC), and betweenness centrality (BC) data of the nodes present in the developed interactive network.

Degree of a node in a network implies the number of edges connected with the given node. Moreover, the degree distribution can be defined as the probability distribution of degrees over the network. In this study, the degree of distribution of the nodes in the regulatory network was studied, and the power-law distribution was observed (**Figure. 2B**). It indicated that the network exhibited the scale-free characteristics alike typical biological networks. We further analyze the indicators of the network centrality such as DC (degree centrality), CC (closeness centrality), and, BC (betweenness centrality) which recognizes the important single nodes possessing significant activity in a complex network (**Supplementary Table S5**). Our result showed significant alteration in the DC, CC and BC among miRNAs and mRNAs (p value < 0.01 for DC, CC, and BC using Mann Whitney U test). Moreover, the miRNA nodes having higher centrality indicating that the miRNA nodes tended to be critical in this context (**Figure. 2C-E**). However, the scale-free distribution of the miRNA-mRNA network indicate the existance of functionally relevant nodes in the development of LUAD.

### 3.4 Functional Enrichment Analysis of the Dysregulated miRNAs and mRNAs

Gene ontology (GO) enrichment analyses for differentially expressed miRNAs and mRNAs were performed. We identified the top 5 most significant GO terms of each group. Differentially expressed miRNAs were enriched in signal transduction, regulation of nucleic acid metabolism, on the BP level. On CC level, they were enriched in nucleus, cytoplasm and lysosome. Transcription factor activity, GTPase activity, protein serine/ threonine kinase activity and phosphatase activity were significant on MF level for dysregulated miRNAs (**Figure. 3A**).

**Figure 3:**
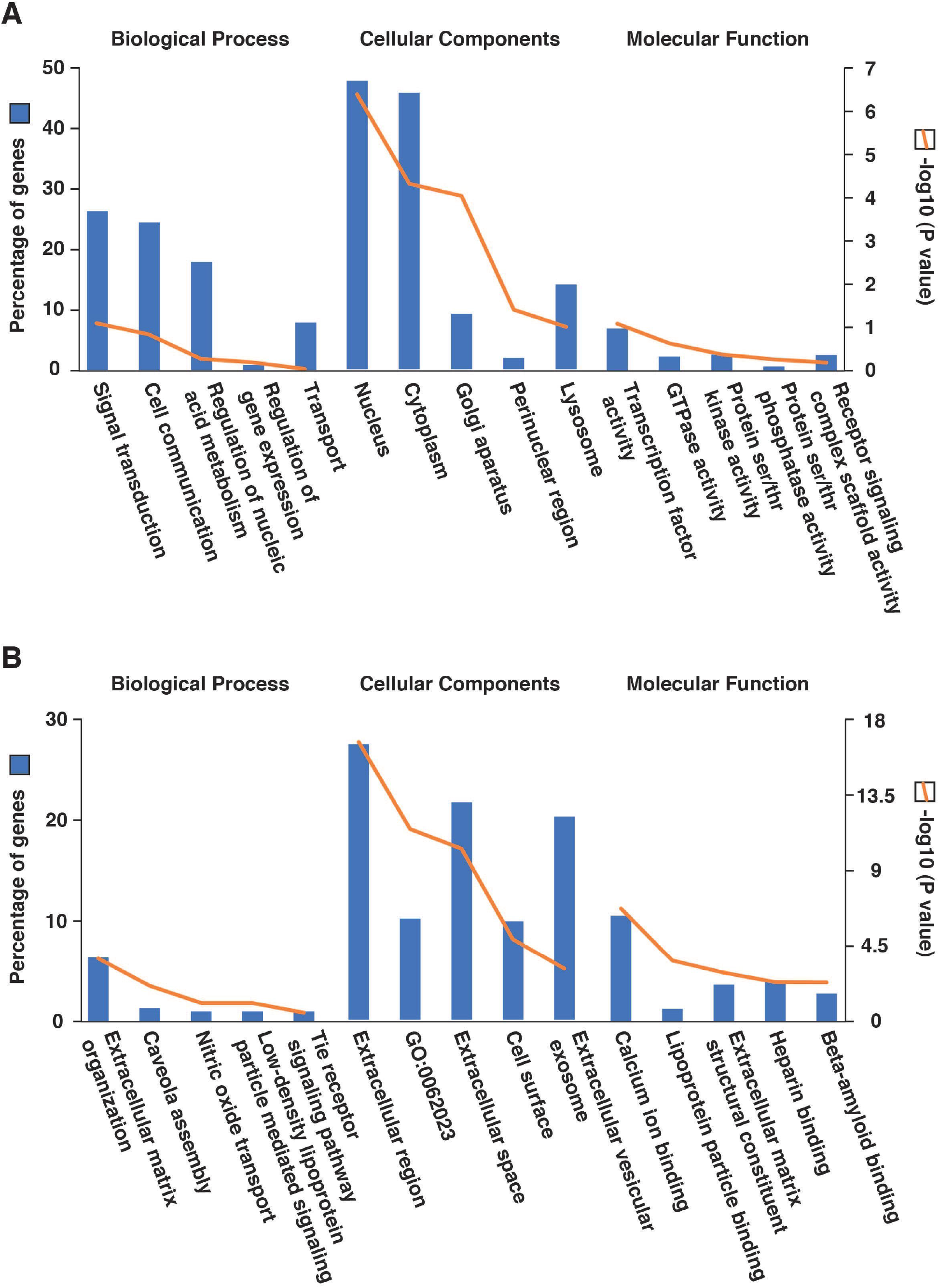
Functional enrichment analysis of the dysregulated miRNA and mRNA signatures. **A-B**. Top five enriched gene ontology terms for each of the biological process, cellular components and molecular functions were shown for miRNA (A) and mRNA (B) signatures using a double y-axis plot. Y-axis in left indicates the percentage of identified dysregulated genes involve in that particular process (represented using blue bars) and y-axis in right indicates the -log10(P-values) (marked using the orange line).

However, differentially expressed mRNAs were enriched in extracellular matrix (ECM) organization, caveola assembly, nitric oxide transport, on the BP level. Cell surface and extracellular regions were enriched on CC level for dysregulated mRNAs. On MF level, calcium ion binding, lipoprotein particle binding, heparin binding were noteworthy (**Figure. 3B**).

### 3.5 Extracellular Matrix Organisation Was the Most Enriched Pathway in LUAD

Considering the signature 429 dysregulated mRNAs, a pathway enrichment analysis was done by using the Reactome pathway analysis database. Top 10 most enhanced pathways were shown (**Figure. 4A**). Extracellular matrix organization pathway was one of the most enriched pathways in the LUAD (p = 1.96 × 10^−5^). Twenty-eight proteins were enriched in the ECM organization pathway. These proteins were further appointed for the network analysis (**Figure. 4B**). Collagen degradation (p = 4.37 × 10^−4^) was among the other key dysregulated pathways in LUAD. The dysregulated proteins involved in the ECM organization pathways were listed in **Supplementary Table S6**.

**Figure 4:**
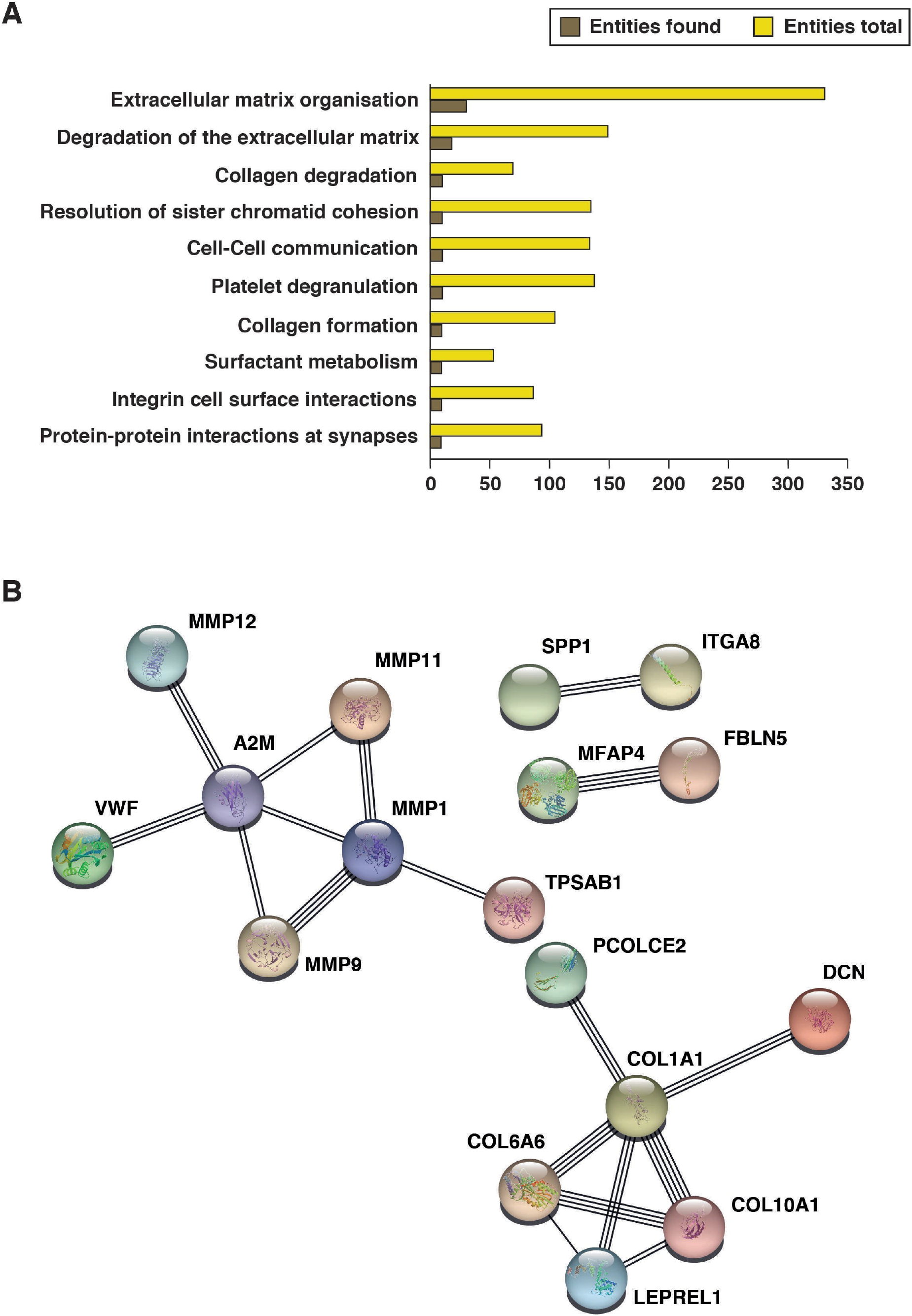
Altered pathways and interaction clusters enriched in LUAD. **A**. Bar graph of showing top ten enriched pathways in lung adenocarcinoma identified using the online pathway analysis tool Reactome (https://reactome.org/). **B**. Protein–protein interaction network comprising the protein clusters associated with the ECM organization pathway with the highest confidence (0.90) obtained using the STRING database (https://string-db.org).

### 3.6 Hub Selection and Analysis

To identify the hub nodes in the network we obtained Cytohubba plug-in of Cytoscape. hsa-miR-486-5p, hsa-miR-200b-5p, hsa-miR-130b-5p, hsa-miR-183-5p, and hsa-miR-139-5p were the top 5 hub miRNAs with the degree 119, 110, 102, 97, and 85 respectively, in the network. On the other hand, ASPM (20), CCNB1 (19), TTN (Titin) (17), CENPF (16), GNG11(15), CCNB2 (15), BIRC5 (15), SPAG5 (15), KIF2C (14) and TPX2 (14) were the top 10 hub genes (with their degree) in the network. The interaction between these hub miRNAs and their common targets were identified and represented (**Figure. 5**).

**Figure 5:**
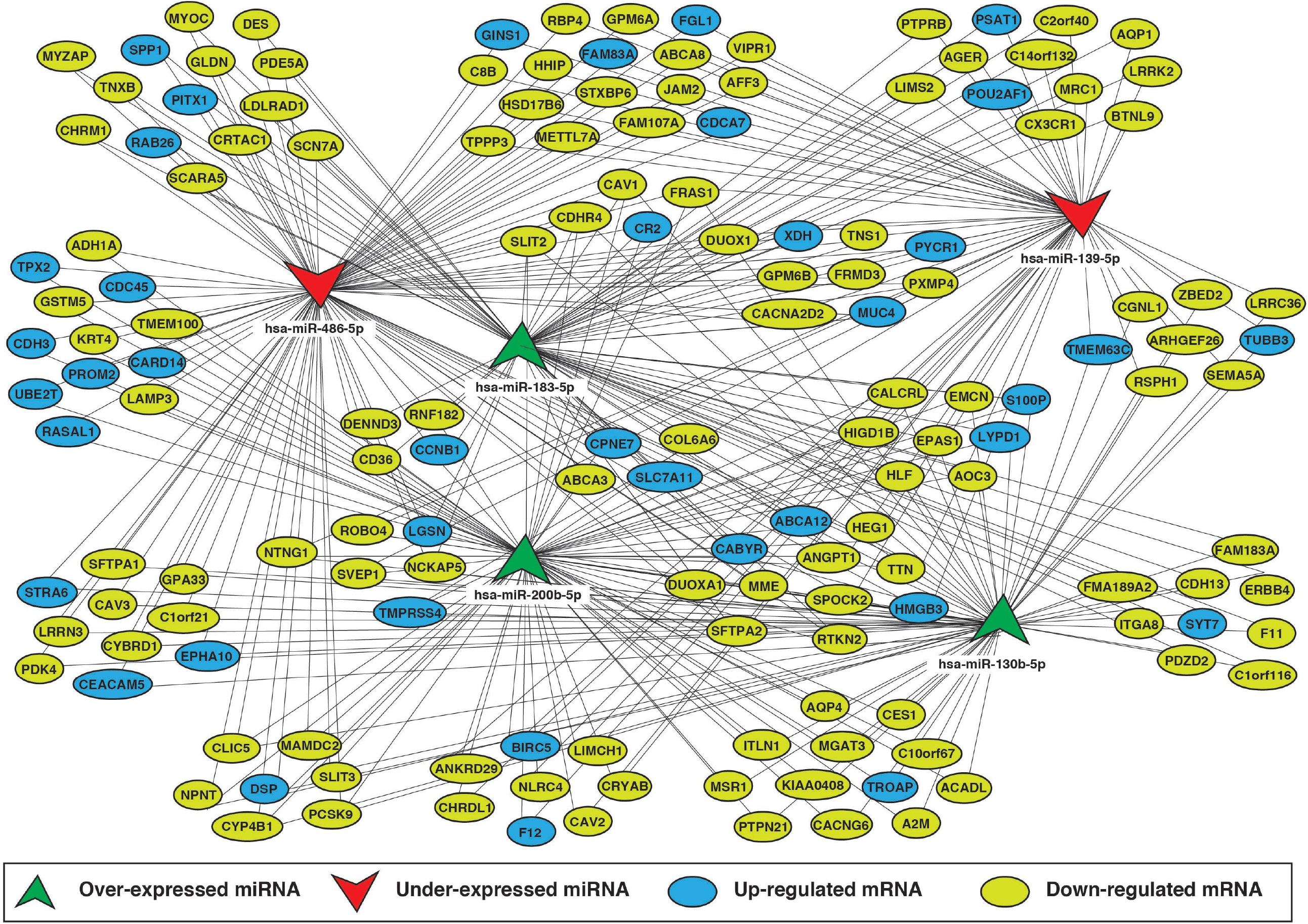
Schematic diagram of interactions between hub miRNA and their common targets. This diagram was visualized in Cytoscape v.3.8.0 (https://cytoscape.org/) and was modified by Adobe Illustrator (Adobe Illustrator CC 2018, Adobe Inc., retrieved from https://adobe.com/products/illustrator).The dysregulated top 5 miRNAs with maximum degree and their common mRNA targets were depicted by a schematic diagram. Their respective expression pattern was also highlighted.

Oncopoint analysis of the hub genes with the cBioPortal showed that TTN (38%) and ASPM (14%) have the exceptionally high genetic mutation rates in LUAD (**Supplementary Figure S2**). We also analyzed the tendency for the cooccurrence of these hub genes (**Supplementary Table S7**). The result showed TTN and ASPM also had a high tendency of cooccurrence in LUAD with a log2 odd ratio 1.368 and p < 0.001.

## 4. Discussion

Understanding the elementary molecular mechanisms of LUAD development to the identification of the therapeutic targets could considerably improve patient prognosis. Large expression profiling data sets have demonstrated to be beneficial in determining the regimen for cancer treatment (Deb et al. 2020; Dong et al. 2020; Klaeger et al. 2017; Zhang et al. 2020). In the post-genomic era, biological networks become a useful tool to understand gene regulation critically. A few studies have been performed to understand the role of miRNAs in LUAD (Jin et al. 2020; Wang et al. 2019). Since the PPI network can easily explain the functions of proteins involved in any disease, it is crucial to understand the effect of miRNA on PPI. In this present report, we have re-investigate and combined the miRNA and mRNA expression data from LUAD patients. We identified 17 differentially expressed miRNAs and 429 differentially expressed mRNAs between lung adenocarcinoma and adjacent normal tissue samples. Biological roles of the dysregulated miRNAs and mRNAs were studied using functional enrichment analyses. The differentially expressed miRNAs were mostly overrepresented in cell communication, signal transduction, regulation of gene expression processes. Earlier studies showed strong evidence suggesting a powerful role of miRNAs in the communication between tumor microenvironment and cancer cells via releasing exosomes, cytokines, growth factors, altering various signaling pathways (Eichelmann et al. 2018). Significantly altered mRNAs were mostly overrepresented in the extracellular matrix organization, caveola assembly. The ECM provides a biomechanical anchorage to the cells. Cancer metastasis is closely related to the detachment of abled cells from the ECM of the primary tumor site. Malignant cells traverse ECM barrier to gain the accessibility of circulation using an anchor independent manner and reattached to the secondary tumor site (Poltavets et al. 2018; Ranganathan et al. 2020). Lim et al. also reported previously the role of ECM organization as a prognostic and predictive indicator for early-stage NSCLC (Lim et al. 2017). Also, the higher CAV1 (Caveolin 1) expression, which leads to caveola assembly, induce filopodia formation, cell migration and metastasis in LUAD cells (Ho et al. 2002). In summary, our results are consistent with these previous reports.

We further analyzed the miRNA-mRNA regulatory network for identifying specific key genes and miRNAs related to LUAD. Among identified top 10 hub genes, ASPM had the highest degree at 20. ASPM gene, which involved in mitotic spindle regulation, is responsible for the coordination of cell cycle. A recent study by Deb et al. showed alteration of phosphorylation pattern in S425 position in ASPM targeted by Serine/threonine-protein kinase Nek2 (NEK2), is a common signature pattern for five cancer types including LUAD (Deb et al. 2020).Wang et al. assessed ASPM expression and the functional significance in LUAD and reported that increasing level of expression of ASPM was positively correlated with poor disease prognosis (Wang et al. 2020). However, ASPM was also reported to be a potential biomarker candidate for bladder cancer(Saleh et al. 2020; Xu et al. 2019) breast cancer (Tang et al. 2019), and prostate cancer (Pai et al. 2019). CCNB1, another hub gene with second highest degree (19), an important mitosis initiator and is a key member of the cyclin family. This gene was previously reported as candidates for the development and prognosis of lung cancer (Arinaga et al. 2003; Soria et al. 2000; Wang et al. 2020), hepatocellular carcinoma (Huang et al. 2018; Long et al. 2019), breast cancer (Ding et al. 2014).In the present study, we found that ASPM and CCNB1 were significantly upregulated in LUAD samples compared to the paired adjacent normal samples. Furthermore, overall survival analysis showed that the LUAD patients having higher expression of these two genes (ASPM and CCNB1) related with the poor overall survival (**Supplementary Figure S3A**).

Moreover, we also identified top 5 miRNA hubs. Among them hsa-miR-200b-5p, hsa-miR-130b-5p and hsa-miR-183-5p were significantly over-expressed with degree 110, 102 and 97. In other hand, hsa-miR-486-5p and hsa-miR-139-5p were under-expressed with degree 119 and 85 respectively. hsa-miR-486-5p, having highest degree in the network, could serve as a potential biomarker for LUAD. Tian et al. validated the significant downregulation of hsa-miR-486-5p in NSCLC tissue, serum and cell samples (Tian et al. 2019). Our data showed hsa-miR-486-5p regulated both ASPM and CCNB1 during the alteration of cellular activities in LUAD patients. In addition, Rui et al. reported a significant involvement of hsa-miR-200b-5p expression in SPC-A1 LUAD cells (Rui et al. 2010). High expression of hsa-miR-130b-5p was associated with the poor overall survival and act as an oncogenic miRNA in NSCLC patients (Hirono et al. 2019). In our study, hsa-miR-130b-5p regulated hub gene SPAG5, CENPF and TTN. The miRNA, hsa-miR-183-5p is involved in cellular migration and invasion (Zaporozhchenko et al. 2016). In contrast to many other studies, we found hsa-miR-183-5p to be upregulated in LUAD patients. However, it plays a significant oncogenic role in other cancer types including breast (Macedo et al. 2017), gastric (Cao et al. 2014), and pancreatic cancer (Lu et al. 2016). Another downregulated miRNA, hsa-miR-139-5p was reported to be associated with occurrence along with development of NSCLC and may serve as a tumor suppressor gene for LUAD(Yong-Hao et al. 2019). The overall survival plots of the hub miRNAs in LUAD were plotted using KM plotter (**Supplementary Figure S3B**).

We further screened the mutation rates of these 10 LUAD hub genes with cBioPortal platform. TTN (38%), ASPM (14%), CENPF (9%), TPX2 (4%) were the four genes with the highest mutation rates, which implies that these genes may play a significant role in tumorigenesis. TTN which is a key component in the vertebrate striated muscles assembly and functioning, has correlation with lung cancer upon missense mutation (Cheng et al. 2019).We observed that TTN was regulated by all five hub miRNAs, worth mentioning. A cooccurrence study among genes showed ASPM had significantly higher cooccurrence log odd ratios with CENPF, TTN, SPAG5, TPX2, and KIF2C. Hub gene KIF2C was reported as a regulator of cell signaling pathway, significantly contribute in LUAD progression (Bai et al. 2019).Survival analysis showed that a higher expression of TPX2, KIF2C is significantly related to the poor overall survival in LUAD patients(Ma et al. 2019) (**Supplementary Figure S3A**). CENPF, another critical hub gene in our network, is associated with the centromere-kinetochore complex and plays a vital role in tumor development in lung (Li et al. 2020), breast (O’Brien et al. 2007; Sun et al. 2019), prostate (Shahid et al. 2018) and nasopharynx (Cao et al. 2010). BIRC5, which is overexpressed in our data, performed a role in apoptosis and cell proliferation (Zhang et al. 2020).CCNB2 which was essential for the regulation of the cell cycle at the G2/M (mitosis) transition, played a significant role in LUAD development (Chen et al. 2020; Zhang et al. 2020). A current study by He et al. reported SPAG5 as “As emerging oncogene” for its overexpression in cancer types (He et al. 2020). SPAG5 was also reported to be upregulated in most of the LUAD cell lines (Kim et al. 2019), which further confirmed our analysis. Nevertheless, these predictions ought to be validated exhaustively through the clinical experiments and accomplished execution in LUAD therapy and treatment.

## 5. Conclusion

From the above discussion it can be concluded that, it is a network-based comprehensive re-investigation of the transcriptomic da ta-sets has been obtained from NCBI GEO. Our study aimed to unveil the role of miRNAs in PPI network of LUAD. miRNA-mRNA regulatory network has identified 5 hub miRNAs and 10 hub mRNAs which may serve as potential biomarkers and can be combined for the treatment of LUAD. However, provided the signaling complexity in cells and tumor heterogeneity, these discoveries require further investigation and validation in larger cohorts of patients.

## Supplementary Materials

**Figure S1:**Protein–protein interaction network showing the top 2 protein clusters with (**A**) GNG11 and (**B**) CCNB2 as hub, involved in the LUAD with the highest confidence (0.90) acquired using the online tool STRING; **Figure S2:** Mutational rate of the hub genes in lung adenocarcinoma were screened using cBioPortal platform; **Figure S3:**Kaplan-Meier plotsshowing overall survival according to high and low expression of mentioned mRNAs (**A**) and miRNAs (**B**) in LUAD;

**Table S1:**Dysregulated (logFC>2) miRNA signature in LUAD; **Table S2:** Dysregulated (logFC>2) mRNA signature in LUAD; **Table S3:** Details of protein-protein interaction network in LUAD; **Table S4:** Interactions of miRNA-mRNA regulatory network in LUAD; **Table S5:** Node details of miRNA-mRNA regulatory network in LUAD; **Table S6:** Proteins enriched in the extracellular matrix organization pathway; **Table S7:** Tendency of cooccurrence of the hub genes in LUAD;

## Author Contributions

Conceptualization, P.S., and M.G.; Methodology, P.S., S.S., and M.M; Software, P.S.; Formal analysis, P.S., M.M. and M.G.; Investigation, P.S., and M.G.; Illustration, P.S.; Resources, M.G.; Data curation, P.S., S.S., and M.M.; Writing—original draft preparation, P.S., and S.S.; Writing—review and editing, M.G.; Visualization, P.S., S.S., M.M., and M.G.; Supervision, M.G.; Project administration, M.G.; All authors have read and agreed to the published version of the manuscript.

## Acknowledgments

We thank the Ministry of Human Resource Development, Government of India and Department of Biotechnology, National Institute of Technology Durgapur for the research support.

## Code availability

Not Applicable

