## Supplementary File 1 for "Integrated Bioinformatics Analysis Deciphering the microRNA Regulation in Protein-Protein Interaction Network in Lung Adenocarcinoma"

### Supplementary Figures

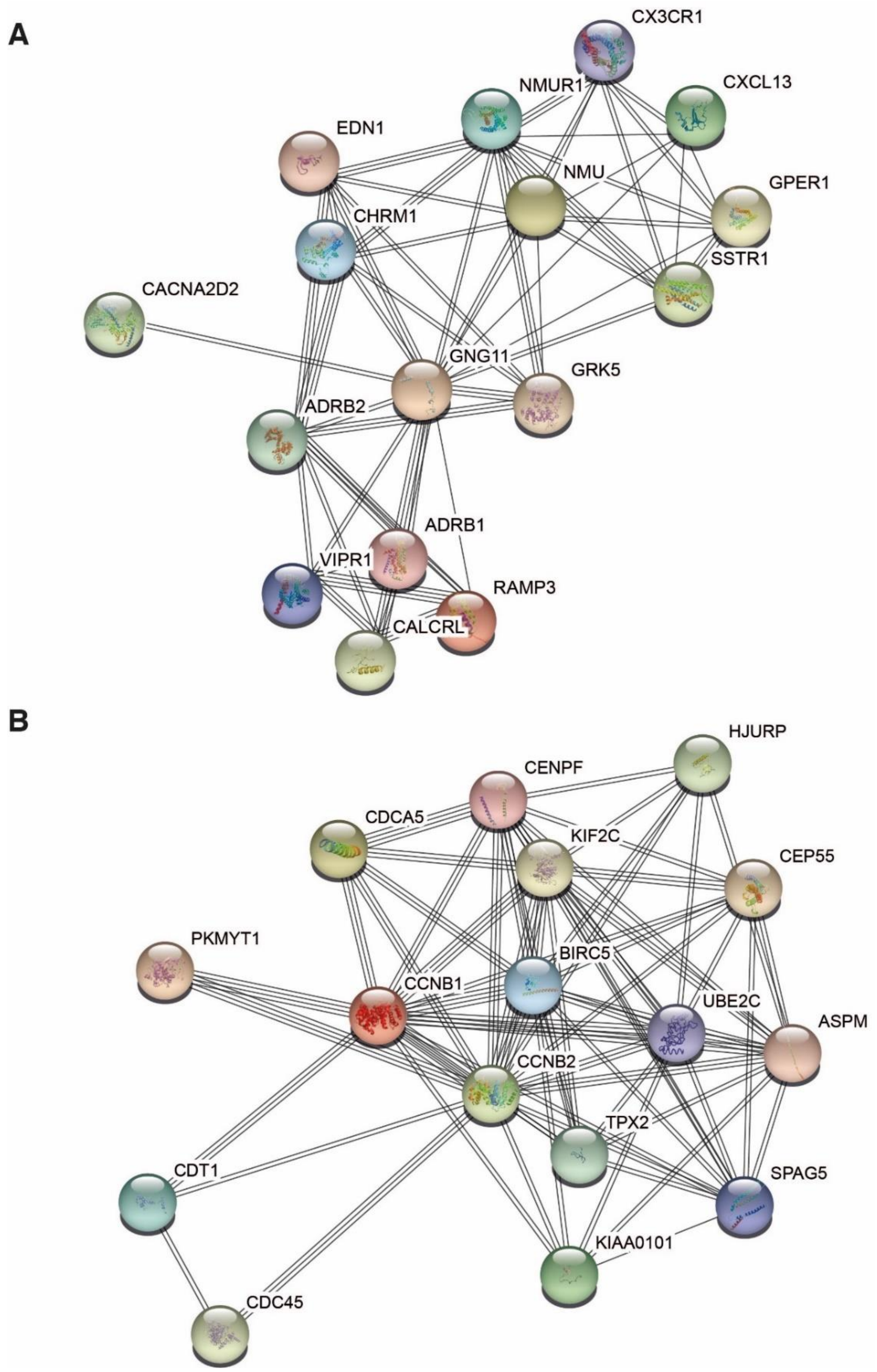


**Supplementary Figure S1:**

Protein‑protein interaction network showing the top 2 protein clusters with (A) GNG11 and (B) CCNB2 as hub, involved in the LUAD with the highest confidence (0.90) acquired using the online tool STRING.

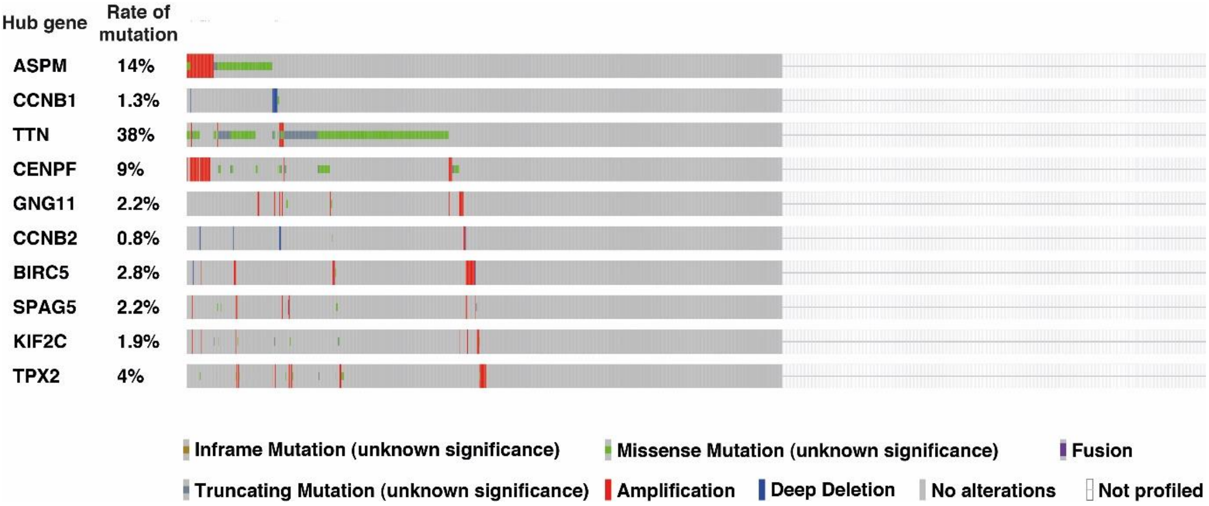


**Supplementary Figure S2:**

Mutational rate of the hub genes in lung adenocarcinoma were screened using cBioPortal platform.

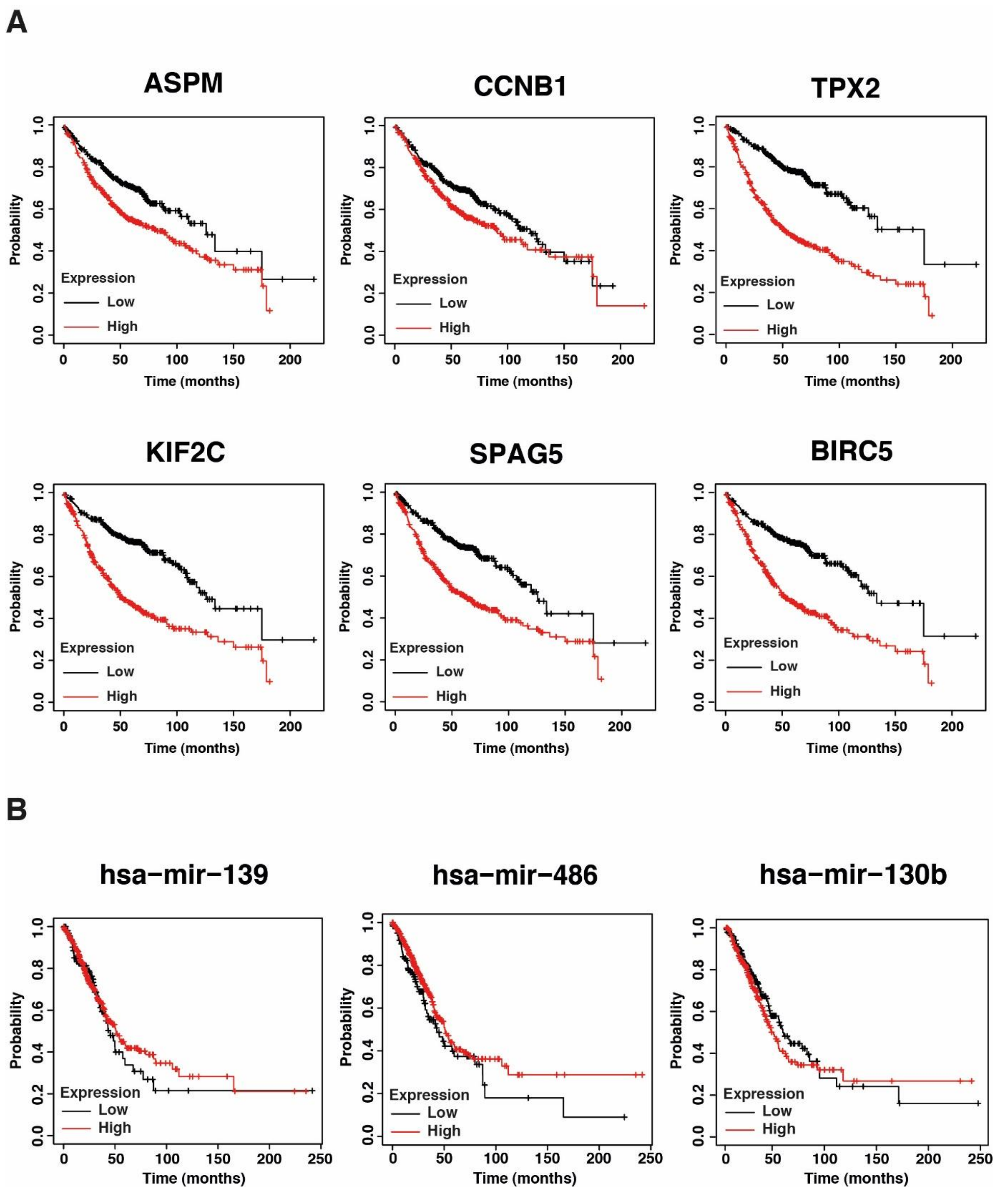


**Supplementary Figure S3:**

Kaplan-Meier plots showing overall survival according to high and low expression of mentioned mRNAs (A) and miRNAs (B) in LUAD.

Sengupta et al. 2020; Integrated Bioinformatics Analysis Decipher the microRNA Regulation in Protein-Protein Interaction Network in Lung Adenocarcinoma

### Supplementary Tables

#### Supplementary Table S1:

Dysregulated (logFC >2) miRNA signature in LUAD

| ID | adj.P.Val | P.Value | t | B | logFC | miRNA_ID |
|---|---|---|---|---|---|---|
| hsa-miR-126 | 2.51E-25 | 3.05E-28 | -16.97988 | 53.87438 | -2.606823 | hsa-miR-126 |
| hsa-miR-144 | 4.12E-21 | 1.00E-23 | -14.29145 | 43.57032 | -4.144946 | hsa-miR-144 |
| hsa-miR-126* | 8.40E-21 | 3.07E-23 | -14.01579 | 42.46067 | -2.39406 | hsa-miR-126* |
| hsa-miR-183 | 9.66E-21 | 4.70E-23 | 13.91119 | 42.03702 | 3.223301 | hsa-miR-183 |
| hsa-miR-218 | 4.23E-19 | 2.58E-21 | -12.94572 | 38.06224 | -2.302513 | hsa-miR-218 |
| hsa-miR-135b | 2.56E-18 | 1.89E-20 | 12.47475 | 36.08223 | 3.20692 | hsa-miR-135b |
| hsa-miR-451 | 2.56E-18 | 2.18E-20 | -12.44139 | 35.94098 | -4.1685 | hsa-miR-451 |
| hsa-miR-486-5p | 3.19E-18 | 3.11E-20 | -12.3584 | 35.58912 | -3.417527 | hsa-miR-486-5p |
| hsa-miR-30a | 9.21E-17 | 1.28E-18 | -11.49705 | 31.89247 | -2.039716 | hsa-miR-30a |
| hsa-miR-139-5p | 3.51E-16 | 5.55E-18 | -11.16276 | 30.43748 | -2.283491 | hsa-miR-139-5p |
| hsa-miR-96 | 2.25E-15 | 3.84E-17 | 10.72498 | 28.51692 | 2.818292 | hsa-miR-96 |
| hsa-miR-200b | 8.12E-15 | 1.48E-16 | 10.42061 | 27.17257 | 2.012788 | hsa-miR-200b |
| hsa-miR-144* | 3.30E-12 | 6.82E-14 | -9.05712 | 21.08639 | -2.372553 | hsa-miR-144* |
| hsa-miR-30a* | 6.81E-12 | 1.49E-13 | -8.8839 | 20.30921 | -2.226259 | hsa-miR-30a* |
| hsa-miR-21 | 4.17E-11 | 1.12E-12 | 8.43939 | 18.3154 | 2.114328 | hsa-miR-21 |
| hsa-miR-338-3p | 2.66E-10 | 8.75E-12 | -7.9835 | 16.27615 | -2.341491 | hsa-miR-338-3p |
| hsa-miR-130b | 2.39E-09 | 8.72E-11 | 7.47168 | 14.00178 | 2.014196 | hsa-miR-130b |

**Supplementary Table S2:**

Dysregulated (logFC >2) mRNA signature in LUAD

| ID | adj.P.Val | P.Value | t | B | logFC | GENE_SYMBOL |
|---|---|---|---|---|---|---|
| A_23_P116898 | 1.09E-05 | 2.63E-07 | -5.7180226 | 6.576717 | -2.00431966 | A2M |
| A_33_P3367984 | 7.75E-05 | 2.72E-06 | 5.1186308 | 4.321956 | 2.42646013 | ABCA12 |
| A_23_P140876 | 3.69E-05 | 1.11E-06 | -5.3518812 | 5.187831 | -2.29932297 | ABCA3 |
| A_33_P3342305 | 8.35E-09 | 5.56E-11 | -7.7785527 | 14.794898 | -2.07434094 | ABCA8 |
| A_23_P218858 | 6.55E-11 | 1.88E-13 | -9.1434345 | 20.335713 | -2.29235724 | ABI3BP |
| A_33_P3303717 | 2.00E-15 | 9.09E-19 | -12.1928823 | 32.204801 | -2.99888749 | ACADL |
| A_33_P3395008 | 1.79E-05 | 4.74E-07 | -5.5690925 | 6.007745 | -2.26501674 | ACOXL |
| A_24_P945113 | 3.68E-08 | 3.28E-10 | -7.3532239 | 13.067407 | -2.13461183 | ACVRL1 |
| A_33_P3308347 | 1.04E-09 | 5.00E-12 | -8.3548172 | 17.139604 | -3.67833104 | ADAMTS8 |
| A_24_P291658 | 1.74E-08 | 1.35E-10 | -7.5670686 | 13.935185 | -3.31767818 | ADH1A |
| A_33_P3353737 | 1.28E-14 | 8.58E-18 | -11.6146536 | 30.034876 | -2.84489057 | ADH1B |
| A_23_P81158 | 3.09E-06 | 5.85E-08 | -6.0936238 | 8.032987 | -3.45092536 | ADH1C |
| A_33_P3310189 | 2.37E-06 | 4.25E-08 | -6.1723551 | 8.341657 | -2.59164817 | ADRB1 |
| A_23_P145024 | 9.76E-11 | 3.07E-13 | -9.0250735 | 19.857847 | -2.68470852 | ADRB2 |
| A_21_P0000190 | 6.49E-08 | 6.43E-10 | -7.191465 | 12.41249 | -2.0364409 | AFF3 |

| | | | | | | |
|---|---|---|---|---|---|---|
| A_23_P93360 | 2.20E-09 | 1.22E-11 | -8.1424975 | 16.275862 | -4.63553788 | AGER |
| A_33_P3279470 | 5.65E-12 | 1.03E-14 | -9.8481081 | 23.160968 | -3.24870331 | AGRP |
| A_23_P93641 | 1.35E-01 | 3.14E-02 | 2.1968965 | -4.406437 | 2.21591867 | AKR1B10 |
| A_23_P104464 | 1.17E-06 | 1.83E-08 | -6.3803328 | 9.161942 | -2.24076894 | ALOX5 |
| A_24_P347378 | 1.39E-07 | 1.56E-09 | -6.9773183 | 11.548133 | -2.36158562 | ALOX5AP |
| A_23_P216023 | 2.78E-09 | 1.58E-11 | -8.0802832 | 16.022679 | -2.29284403 | ANGPT1 |
| A_23_P412577 | 4.83E-07 | 6.49E-09 | -6.6329201 | 10.166541 | -2.08224898 | ANKRD29 |
| A_23_P121716 | 1.44E-08 | 1.06E-10 | -7.6232802 | 14.163577 | -2.55398887 | ANXA3 |
| A_32_P105549 | 1.50E-04 | 6.15E-06 | -4.9039676 | 3.53999 | -3.04403436 | ANXA8L2 |
| A_23_P426305 | 1.89E-10 | 6.59E-13 | -8.8411985 | 19.113975 | -2.91316992 | AOC3 |
| A_23_P372834 | 5.35E-04 | 2.89E-05 | -4.4845143 | 2.0592 | -2.09001653 | AQP1 |
| A_33_P3420757 | 1.81E-05 | 4.81E-07 | -5.5651892 | 5.992904 | -2.6452933 | AQP4 |
| A_23_P69100 | 7.56E-12 | 1.51E-14 | -9.7543297 | 22.787201 | -2.37828229 | ARHGEF26 |
| A_23_P52017 | 6.22E-07 | 8.73E-09 | 6.5604819 | 9.877591 | 2.33708984 | ASPM |
| A_23_P78980 | 6.35E-09 | 4.04E-11 | 7.8548805 | 15.105411 | 3.60842133 | B3GNT3 |
| A_23_P212050 | 1.29E-09 | 6.55E-12 | -8.290349 | 16.877409 | -2.07206345 | BCHE |
| A_23_P118815 | 3.46E-07 | 4.36E-09 | 6.7296397 | 10.553306 | 3.05025674 | BIRC5 |

| | | | | | | |
|---|---|---|---|---|---|---|
| A_33_P3312466 | 3.95E-12 | 6.49E-15 | -9.9605351 | 23.608052 | -2.57231303 | BTNL9 |
| A_23_P161439 | 2.64E-08 | 2.21E-10 | -7.448186 | 13.452523 | -3.22396381 | C10orf116 |
| A_33_P3376806 | 2.00E-14 | 1.41E-17 | -11.4885841 | 29.556142 | -2.29638099 | C10orf67 |
| A_32_P225345 | 1.79E-04 | 7.65E-06 | -4.8456404 | 3.330191 | -2.03331298 | C11orf88 |
| A_24_P10137 | 7.23E-11 | 2.16E-13 | -9.1097392 | 20.199756 | -2.67335048 | C13orf15 |
| A_23_P151529 | 1.95E-13 | 1.85E-16 | -10.8395799 | 27.06128 | -2.80393219 | C14orf132 |
| A_23_P26024 | 1.12E-03 | 7.14E-05 | 4.2306263 | 1.197312 | 2.12449874 | C15orf48 |
| A_23_P26557 | 1.43E-08 | 1.05E-10 | 7.6254663 | 14.172461 | 2.34698987 | C16orf59 |
| A_23_P330561 | 1.10E-10 | 3.52E-13 | -8.9921856 | 19.724926 | -4.30755088 | C19orf59 |
| A_23_P897 | 1.47E-03 | 1.00E-04 | -4.1332961 | 0.874515 | -2.09519008 | C1orf116 |
| A_32_P15512 | 3.28E-04 | 1.60E-05 | -4.6474162 | 2.626356 | -2.71299132 | C1orf194 |
| A_23_P113161 | 3.96E-08 | 3.62E-10 | -7.3296858 | 12.972016 | -2.06919767 | C1orf21 |
| A_23_P314835 | 6.53E-04 | 3.68E-05 | -4.4176779 | 1.829623 | -3.27216401 | C20orf85 |
| A_33_P3356462 | 1.79E-04 | 7.65E-06 | 4.8456944 | 3.330385 | 2.36596172 | C2CD4A |
| A_33_P3289121 | 1.93E-08 | 1.52E-10 | -7.5375096 | 13.815127 | -3.59564883 | C2orf40 |
| A_23_P362694 | 4.09E-04 | 2.09E-05 | 4.5741389 | 2.369929 | 2.84950382 | C4orf7 |
| A_33_P3284453 | 1.22E-10 | 3.99E-13 | -8.9616209 | 19.601343 | -2.53890491 | C6orf174 |

| | | | | | | |
|---|---|---|---|---|---|---|
| A_33_P3423854 | 1.17E-06 | 1.83E-08 | -6.3803276 | 9.161922 | -2.46827557 | C8B |
| A_23_P170649 | 4.13E-07 | 5.44E-09 | -6.6758526 | 10.33809 | -2.10767307 | C8orf84 |
| A_32_P181527 | 1.32E-05 | 3.27E-07 | -5.662979 | 6.365821 | -2.08549675 | C8orf85 |
| A_33_P3354847 | 1.49E-04 | 6.08E-06 | -4.9070514 | 3.551115 | -2.64637901 | C9orf171 |
| A_23_P217009 | 1.18E-04 | 4.57E-06 | -4.9826425 | 3.824825 | -3.29680147 | C9orf24 |
| A_23_P8913 | 8.40E-05 | 3.00E-06 | -5.0933928 | 4.229248 | -2.15064316 | CA2 |
| A_23_P4096 | 1.07E-16 | 2.23E-20 | -13.1717778 | 35.777557 | -3.43420754 | CA4 |
| A_23_P314712 | 2.06E-05 | 5.61E-07 | 5.5265197 | 5.846084 | 2.59047077 | CABYR |
| A_23_P346900 | 1.22E-07 | 1.34E-09 | -7.0151607 | 11.700621 | -2.98497462 | CACNA2D2 |
| A_23_P501933 | 6.34E-04 | 3.54E-05 | -4.4279482 | 1.864779 | -2.14793272 | CACNG6 |
| A_33_P3306110 | 9.16E-13 | 1.24E-15 | -10.36689 | 25.21417 | -2.36356565 | CALCRL |
| A_23_P253791 | 2.27E-08 | 1.84E-10 | -7.4914719 | 13.6282 | -2.4206342 | CAMP |
| A_23_P207879 | 2.43E-04 | 1.10E-05 | 4.7486135 | 2.983868 | 2.01836256 | CARD14 |
| A_23_P134454 | 1.39E-11 | 3.07E-14 | -9.5814424 | 22.096228 | -3.69317166 | CAV1 |
| A_24_P251599 | 2.05E-14 | 1.50E-17 | -11.4725845 | 29.495245 | -2.13005541 | CAV3 |
| A_23_P166566 | 3.68E-12 | 5.97E-15 | -9.9807409 | 23.688284 | -3.06724998 | CCDC48 |
| A_23_P349566 | 2.31E-10 | 8.53E-13 | -8.7789755 | 18.861881 | -2.14786142 | CCDC85A |

| | | | | | | |
|---|---|---|---|---|---|---|
| A_23_P218369 | 3.77E-04 | 1.88E-05 | -4.6024117 | 2.468618 | -2.12653666 | CCL14 |
| A_23_P122197 | 1.47E-05 | 3.71E-07 | 5.6312163 | 6.244444 | 2.01522807 | CCNB1 |
| A_23_P65757 | 4.73E-06 | 9.61E-08 | 5.9703111 | 7.551783 | 2.45154155 | CCNB2 |
| A_23_P6909 | 9.08E-10 | 4.27E-12 | -8.3926979 | 17.293629 | -2.10263928 | CCRL1 |
| A_23_P113572 | 9.90E-04 | 6.13E-05 | 4.2738427 | 1.34203 | 2.01713489 | CD19 |
| A_23_P111583 | 2.88E-10 | 1.09E-12 | -8.7197603 | 18.621816 | -2.70143935 | CD36 |
| A_23_P85800 | 2.44E-05 | 6.72E-07 | -5.4802535 | 5.670917 | -2.15880773 | CD52 |
| A_23_P57379 | 1.34E-06 | 2.14E-08 | 6.3413084 | 9.007516 | 2.44841759 | CDC45 |
| A_23_P104651 | 5.10E-06 | 1.05E-07 | 5.9484322 | 7.466706 | 2.44144786 | CDCA5 |
| A_23_P251421 | 7.67E-09 | 5.02E-11 | 7.8032886 | 14.895518 | 2.73838671 | CDCA7 |
| A_32_P85999 | 3.40E-10 | 1.33E-12 | -8.6722643 | 18.429161 | -2.16234902 | CDH13 |
| A_23_P49155 | 1.51E-04 | 6.19E-06 | 4.902348 | 3.534149 | 2.3372555 | CDH3 |
| A_33_P3396537 | 4.18E-04 | 2.14E-05 | -4.5671223 | 2.345486 | -2.46842059 | CDHR4 |
| A_23_P30294 | 2.04E-11 | 5.08E-14 | -9.4594447 | 21.607235 | -2.61391388 | CDO1 |
| A_33_P3386262 | 2.34E-07 | 2.81E-09 | 6.8355622 | 10.97803 | 2.54268008 | CDT1 |
| A_23_P153301 | 1.51E-02 | 1.79E-03 | 3.2520836 | -1.825089 | 2.65668808 | CEACAM5 |
| A_23_P401 | 3.34E-07 | 4.18E-09 | 6.7395083 | 10.592827 | 2.4875497 | CENPF |
| A_23_P115872 | 1.31E-05 | 3.24E-07 | 5.6654577 | 6.375302 | 2.24486806 | CEP55 |
| A_33_P3241269 | 4.89E-04 | 2.60E-05 | -4.5141393 | 2.16155 | -2.33176439 | CES1 |

| | | | | | | |
|---|---|---|---|---|---|---|
| A_23_P119562 | 3.07E-08 | 2.64E-10 | -7.4057966 | 13.280562 | -2.45556255 | CFD |
| A_23_P163306 | 7.94E-08 | 8.23E-10 | -7.1323168 | 12.173418 | -2.29774125 | CGNL1 |
| A_23_P34888 | 1.62E-02 | 1.96E-03 | -3.2216991 | -1.910183 | -2.07192146 | CHIA |
| A_24_P168925 | 2.55E-08 | 2.12E-10 | -7.4582277 | 13.493271 | -2.96053417 | CHRDL1 |
| A_33_P3367860 | 1.07E-16 | 2.01E-20 | -13.1992232 | 35.875851 | -2.88733897 | CHRM1 |
| A_33_P3240752 | 4.06E-07 | 5.33E-09 | -6.6805061 | 10.356697 | -5.07813088 | CLDN18 |
| A_33_P3285540 | 8.66E-06 | 1.96E-07 | -5.7924961 | 6.863145 | -2.22984543 | CLDN5 |
| A_24_P215653 | 2.53E-09 | 1.42E-11 | -8.1052828 | 16.124418 | -2.11887512 | CLEC14A |
| A_23_P69497 | 1.15E-09 | 5.68E-12 | -8.3244403 | 17.01607 | -3.56457664 | CLEC3B |
| A_33_P3336686 | 5.98E-06 | 1.27E-07 | -5.9017213 | 7.285385 | -2.28573584 | CLIC3 |
| A_23_P416774 | 2.97E-08 | 2.53E-10 | -7.4156335 | 13.32046 | -3.70620582 | CLIC5 |
| A_23_P214144 | 3.06E-06 | 5.76E-08 | 6.0972237 | 8.047077 | 2.68598893 | COL10A1 |
| A_33_P3304668 | 2.69E-05 | 7.59E-07 | 5.4492249 | 5.55375 | 2.75292157 | COL1A1 |
| A_32_P224525 | 1.24E-07 | 1.36E-09 | -7.0103971 | 11.68142 | -2.52698735 | COL6A6 |
| A_23_P67661 | 1.85E-09 | 9.93E-12 | -8.1907828 | 16.472342 | -2.14283145 | COX7A1 |
| A_23_P18017 | 7.78E-06 | 1.73E-07 | -5.8232862 | 6.981918 | -2.05726812 | CPA3 |
| A_23_P67198 | 1.47E-04 | 5.96E-06 | -4.9122156 | 3.569753 | -2.32364754 | CPAMD8 |

| | | | | | | |
|---|---|---|---|---|---|---|
| A_33_P3262191 | 2.15E-04 | 9.52E-06 | 4.786945 | 3.12028 | 2.03488341 | CPNE7 |
| A_33_P3281985 | 4.24E-03 | 3.69E-04 | 3.7494497 | -0.353469 | 2.02316484 | CR2 |
| A_23_P115064 | 2.61E-05 | 7.28E-07 | 5.4597052 | 5.593296 | 3.221779 | CRABP2 |
| A_23_P431971 | 6.50E-07 | 9.22E-09 | -6.5471317 | 9.82441 | -2.37269586 | CRTAC1 |
| A_24_P206776 | 1.54E-09 | 7.99E-12 | -8.2429773 | 16.684703 | -2.2632908 | CRYAB |
| A_24_P211044 | 5.54E-11 | 1.53E-13 | -9.1921889 | 20.532311 | -2.80906138 | CSH1 |
| A_23_P111888 | 7.89E-07 | 1.15E-08 | 6.4930209 | 9.609088 | 2.66797052 | CTHRC1 |
| A_23_P407565 | 4.42E-07 | 5.89E-09 | -6.6561852 | 10.259477 | -2.38341407 | CX3CR1 |
| A_23_P121695 | 2.31E-04 | 1.04E-05 | 4.7643021 | 3.039635 | 2.23630232 | CXCL13 |
| A_33_P3590259 | 2.64E-02 | 3.65E-03 | 3.0112608 | -2.483219 | 2.29346311 | CXCL14 |
| A_23_P209564 | 8.37E-08 | 8.75E-10 | -7.1175715 | 12.113856 | -2.30995427 | CYBRD1 |
| A_23_P89981 | 1.52E-05 | 3.85E-07 | -5.6217456 | 6.208299 | -2.75716859 | CYP2F1 |
| A_23_P114713 | 2.88E-07 | 3.52E-09 | -6.7810276 | 10.759215 | -4.03752374 | CYP4B1 |
| A_33_P3398156 | 4.33E-05 | 1.34E-06 | -5.3030272 | 5.005167 | -2.13696455 | CYS1 |
| A_33_P3246028 | 1.21E-04 | 4.69E-06 | -4.9756068 | 3.799269 | -2.30585188 | DCDC2B |
| A_33_P3382856 | 4.05E-06 | 8.04E-08 | -6.0145253 | 7.723994 | -2.0318889 | DCN |
| A_23_P31816 | 5.55E-04 | 3.02E-05 | -4.4725981 | 2.018133 | -2.16454793 | DEFA3 |
| A_32_P175301 | 5.14E-10 | 2.19E-12 | -8.5525835 | 17.943339 | -2.04711989 | DENND3 |

| | | | | | | |
|---|---|---|---|---|---|---|
| A_33_P3275801 | 2.53E-06 | 4.58E-08 | -6.1541439 | 8.270164 | -2.13725492 | DES |
| A_24_P940115 | 6.01E-06 | 1.27E-07 | -5.9000749 | 7.279002 | -2.11557121 | DLC1 |
| A_32_P14721 | 3.80E-04 | 1.90E-05 | -4.6002778 | 2.461159 | -2.40061376 | DNAH12 |
| A_33_P3285629 | 1.25E-04 | 4.90E-06 | -4.9639476 | 3.756953 | -2.85360231 | DNAI2 |
| A_33_P3234202 | 2.20E-09 | 1.22E-11 | -8.14077 | 16.268832 | -3.00044514 | DNASE1L3 |
| A_33_P3402565 | 9.14E-05 | 3.34E-06 | 5.0654018 | 4.126662 | 2.11036923 | DSP |
| A_23_P54291 | 4.30E-07 | 5.71E-09 | -6.6637259 | 10.289612 | -2.95406228 | DUOX1 |
| A_33_P3410235 | 4.84E-06 | 9.86E-08 | -5.9637746 | 7.526356 | -2.16450113 | DUOXA1 |
| A_23_P94840 | 4.47E-04 | 2.33E-05 | -4.5441722 | 2.265674 | -2.31970505 | DYNLRB2 |
| A_33_P3610406 | 1.71E-08 | 1.32E-10 | -7.5722512 | 13.956239 | -2.99521058 | ECEL1P2 |
| A_23_P214821 | 1.28E-04 | 5.04E-06 | -4.9566398 | 3.730454 | -2.01434199 | EDN1 |
| A_33_P3229953 | 9.54E-03 | 1.01E-03 | 3.4366743 | -1.295859 | 2.87788973 | EEF1A2 |
| A_23_P146294 | 5.22E-05 | 1.67E-06 | -5.2459759 | 4.792712 | -2.51658069 | EFCAB1 |
| A_23_P382065 | 2.97E-08 | 2.54E-10 | -7.4147545 | 13.316895 | -2.44547429 | EMCN |
| A_23_P304682 | 6.85E-13 | 8.40E-16 | -10.4634511 | 25.593426 | -2.98023977 | EMP2 |
| A_33_P3420224 | 9.00E-06 | 2.05E-07 | 5.7804135 | 6.816591 | 2.56585203 | ENTPD8 |
| A_23_P210210 | 5.09E-08 | 4.87E-10 | -7.2583081 | 12.682935 | -2.18844147 | EPAS1 |

| | | | | | | |
|---|---|---|---|---|---|---|
| A_24_P930741 | 1.91E-05 | 5.10E-07 | 5.550559 | 5.937312 | 2.04441511 | EPHA10 |
| A_23_P130027 | 2.83E-08 | 2.39E-10 | 7.4289672 | 13.374548 | 2.19253263 | EPN3 |
| A_32_P183765 | 6.53E-05 | 2.21E-06 | -5.1732717 | 4.523347 | -2.09741593 | ERBB4 |
| A_23_P41314 | 1.77E-07 | 2.05E-09 | -6.9123982 | 11.286817 | -2.42336058 | F11 |
| A_33_P3233871 | 2.62E-05 | 7.31E-07 | 5.4586767 | 5.589414 | 2.1873739 | F12 |
| A_23_P8820 | 8.51E-14 | 7.40E-17 | -11.0689395 | 27.948635 | -2.65943025 | FABP4 |
| A_23_P59877 | 2.90E-06 | 5.38E-08 | -6.114112 | 8.113208 | -2.13285124 | FABP5 |
| A_23_P84860 | 4.72E-10 | 1.97E-12 | -8.5785819 | 18.048916 | -3.5244893 | FAM107A |
| A_24_P561341 | 1.48E-11 | 3.40E-14 | -9.556599 | 21.996741 | -2.3488788 | FAM150B |
| A_23_P145054 | 5.66E-07 | 7.88E-09 | -6.5855209 | 9.977397 | -2.26489029 | FAM162B |
| A_33_P3244643 | 2.14E-03 | 1.60E-04 | -3.9979399 | 0.433057 | -2.30549347 | FAM183A |
| A_23_P422831 | 8.17E-08 | 8.52E-10 | -7.1238615 | 12.139262 | -2.91432094 | FAM189A2 |
| A_23_P216052 | 2.34E-04 | 1.05E-05 | 4.7595991 | 3.022908 | 2.01980406 | FAM83A |
| A_23_P151805 | 8.75E-09 | 5.92E-11 | -7.7638109 | 14.734937 | -2.38297096 | FBLN5 |
| A_23_P115246 | 2.88E-10 | 1.09E-12 | -8.7199287 | 18.6225 | -4.05819574 | FCN3 |
| A_23_P201211 | 1.99E-04 | 8.71E-06 | 4.8108827 | 3.205739 | 2.225894 | FCRL5 |
| A_23_P80048 | 1.57E-08 | 1.18E-10 | 7.597805 | 14.060057 | 3.09739251 | FER1L4 |
| A_23_P131935 | 1.66E-05 | 4.30E-07 | 5.5940226 | 6.102619 | 2.0961918 | FERMT1 |
| A_33_P3364180 | 6.57E-08 | 6.55E-10 | -7.1873443 | 12.395827 | -2.01267662 | FGD5 |

| | | | | | | |
|---|---|---|---|---|---|---|
| A_23_P20484 | 2.43E-02 | 3.29E-03 | 3.0468515 | -2.388345 | 2.78370799 | FGL1 |
| A_23_P108751 | 8.37E-08 | 8.76E-10 | 7.1171096 | 12.11199 | 2.38708014 | FHL2 |
| A_33_P3325497 | 1.73E-11 | 4.14E-14 | -9.5088649 | 21.805456 | -2.20813977 | FIBIN |
| A_23_P45185 | 5.79E-10 | 2.52E-12 | -8.5192654 | 17.808006 | -3.69666846 | FIGF |
| A_33_P3806965 | 6.05E-03 | 5.74E-04 | 3.6137915 | -0.768998 | 2.5382852 | FLJ13744 |
| A_33_P3480395 | 8.23E-13 | 1.09E-15 | -10.3990164 | 25.340456 | -2.99332327 | FLJ30901 |
| A_21_P0006454 | 1.04E-08 | 7.22E-11 | -7.7162242 | 14.541411 | -2.43488674 | FLJ46446 |
| A_33_P3275702 | 1.13E-12 | 1.57E-15 | -10.3088815 | 24.985877 | -3.70624505 | FMO2 |
| A_23_P118254 | 6.89E-13 | 8.74E-16 | -10.4534606 | 25.554232 | -2.24155966 | FOXF1 |
| A_32_P107876 | 7.20E-06 | 1.58E-07 | -5.8465657 | 7.071853 | -2.14445063 | FRAS1 |
| A_33_P3363425 | 2.22E-11 | 5.66E-14 | -9.4332223 | 21.501986 | -2.22261559 | FRMD3 |
| A_23_P360316 | 5.97E-05 | 1.97E-06 | 5.2034086 | 4.634811 | 2.06360405 | FUT3 |
| A_33_P3228435 | 1.66E-12 | 2.43E-15 | -10.2010781 | 24.560724 | -2.88057905 | FXYD1 |
| A_23_P67847 | 1.01E-04 | 3.75E-06 | 5.034487 | 4.01365 | 2.66439233 | GALNT14 |
| A_23_P52227 | 1.79E-11 | 4.34E-14 | -9.4975272 | 21.759998 | -3.68726996 | GDF10 |
| A_23_P145631 | 2.76E-09 | 1.56E-11 | -8.0823827 | 16.031223 | -2.0357933 | GIMAP6 |
| A_24_P132383 | 1.51E-10 | 5.13E-13 | -8.9011917 | 19.356866 | -2.2315048 | GIMAP8 |
| A_33_P3340025 | 1.22E-06 | 1.90E-08 | 6.3699013 | 9.120641 | 2.15806831 | GINS1 |

| | | | | | | |
|---|---|---|---|---|---|---|
| A_23_P204947 | 2.48E-07 | 3.02E-09 | 6.8184775 | 10.909448 | 2.71438254 | GJB2 |
| A_23_P61317 | 5.00E-06 | 1.03E-07 | -5.9540827 | 7.488669 | -3.55179364 | GKN2 |
| A_24_P49199 | 1.38E-08 | 1.01E-10 | -7.6364002 | 14.216899 | -2.03893016 | GLDN |
| A_23_P414913 | 2.32E-07 | 2.79E-09 | -6.8377013 | 10.986619 | -2.24361961 | GLIPR2 |
| A_23_P111701 | 1.03E-05 | 2.44E-07 | -5.7370336 | 6.649717 | -2.0087754 | GNG11 |
| A_23_P146512 | 3.09E-08 | 2.66E-10 | 7.4037402 | 13.272222 | 2.23122987 | GOLM1 |
| A_24_P319374 | 7.47E-09 | 4.86E-11 | -7.811095 | 14.927274 | -3.00307096 | GPA33 |
| A_33_P3230264 | 1.67E-08 | 1.28E-10 | -7.5788937 | 13.983223 | -2.2717655 | GPC3 |
| A_23_P8640 | 9.25E-05 | 3.38E-06 | -5.0617852 | 4.113425 | -2.17093871 | GPER |
| A_23_P72697 | 4.39E-09 | 2.67E-11 | -7.9540675 | 15.509019 | -3.53714192 | GPIHBP1 |
| A_23_P121545 | 5.37E-22 | 1.06E-26 | -17.378041 | 49.63049 | -2.58067914 | GPM6A |
| A_33_P3218649 | 5.63E-12 | 1.01E-14 | -9.8514747 | 23.174372 | -2.43033942 | GPM6B |
| A_33_P3243429 | 2.21E-16 | 6.11E-20 | -12.902499 | 34.80768 | -4.33056047 | GPR152 |
| A_33_P3285299 | 1.68E-05 | 4.39E-07 | -5.5886743 | 6.082253 | -2.01900578 | GPRIN2 |
| A_23_P3038 | 7.45E-02 | 1.42E-02 | 2.5183506 | -3.706365 | 2.05438221 | GPX2 |
| A_23_P133474 | 4.64E-07 | 6.21E-09 | -6.6436498 | 10.209394 | -2.53487116 | GPX3 |
| A_23_P12884 | 5.39E-10 | 2.32E-12 | -8.5388888 | 17.887717 | -2.40969982 | GRK5 |

| | | | | | | |
|---|---|---|---|---|---|---|
| A_23_P97606 | 1.64E-12 | 2.36E-15 | -10.2084728 | 24.589923 | -2.25929505 | GSTM5 |
| A_24_P142305 | 8.45E-07 | 1.25E-08 | -6.4726705 | 9.528208 | -3.21713486 | HBA2 |
| A_23_P203558 | 6.82E-06 | 1.49E-07 | -5.8612881 | 7.128789 | -3.03592979 | HBB |
| A_24_P75190 | 1.98E-06 | 3.42E-08 | -6.2257456 | 8.551578 | -3.34496435 | HBD |
| A_32_P166693 | 7.02E-10 | 3.13E-12 | -8.4669501 | 17.595445 | -2.24871498 | HEG1 |
| A_21_P0014446 | 6.92E-08 | 6.98E-10 | -7.1719648 | 12.333646 | -3.43430875 | HHIP |
| A_23_P207354 | 8.75E-09 | 5.89E-11 | -7.7647665 | 14.738824 | -2.49976112 | HIGD1B |
| A_33_P3360216 | 5.78E-06 | 1.21E-07 | 5.9117997 | 7.32447 | 2.03926992 | HIST1H2AI |
| A_33_P3257678 | 3.01E-04 | 1.43E-05 | 4.6771513 | 2.731003 | 2.24269161 | HIST2H3A |
| A_33_P3807062 | 1.12E-06 | 1.72E-08 | 6.3945164 | 9.218125 | 2.69773319 | HJURP |
| A_33_P3400578 | 6.27E-07 | 8.82E-09 | -6.5580846 | 9.86804 | -2.65437793 | HLF |
| A_33_P3319041 | 5.76E-06 | 1.21E-07 | 5.9130546 | 7.329338 | 2.66400532 | HMGB3 |
| A_33_P3237359 | 1.38E-05 | 3.43E-07 | 5.6507969 | 6.31924 | 2.22338011 | HMGB3 |
| A_23_P373119 | 1.34E-06 | 2.15E-08 | 6.3398024 | 9.001561 | 2.52940374 | HMGB3P1 |
| A_24_P105191 | 1.45E-07 | 1.64E-09 | 6.9656513 | 11.501144 | 2.909769 | HS6ST2 |
| A_23_P25030 | 3.40E-08 | 2.97E-10 | -7.3775316 | 13.165945 | -3.22656625 | HSD17B6 |
| A_23_P91334 | 2.98E-11 | 7.84E-14 | -9.3542148 | 21.184574 | -2.2281929 | HSPA12B |
| A_23_P161727 | 8.92E-10 | 4.17E-12 | -8.3985414 | 17.317386 | -2.25761107 | HSPB2 |
| A_33_P3338698 | 1.12E-04 | 4.25E-06 | -5.0018189 | 3.894566 | -2.4510454 | IHH |

| | | | | | | |
|---|---|---|---|---|---|---|
| A_33_P3248394 | 5.85E-16 | 1.84E-19 | -12.610278 | 33.743998 | -2.57437206 | INMT |
| A_33_P3347291 | 1.28E-08 | 9.11E-11 | -7.660385 | 14.314388 | -3.63818545 | INMT |
| A_33_P3321293 | 3.97E-08 | 3.64E-10 | 7.3286558 | 12.967843 | 2.37343694 | IQGAP3 |
| A_33_P3336257 | 5.58E-07 | 7.73E-09 | -6.5901422 | 9.995826 | -2.92482711 | IRX1 |
| A_23_P46781 | 1.79E-10 | 6.20E-13 | -8.8557974 | 19.173097 | -2.36881604 | ITGA8 |
| A_23_P95790 | 8.57E-04 | 5.15E-05 | -4.3232261 | 1.50842 | -2.3196504 | ITLN1 |
| A_24_P53778 | 4.61E-12 | 7.92E-15 | -9.9116707 | 23.413872 | -4.8040547 | ITLN2 |
| A_23_P171074 | 2.23E-07 | 2.66E-09 | -6.8491334 | 11.032529 | -2.03794423 | ITM2A |
| A_23_P65918 | 7.67E-06 | 1.70E-07 | 5.8270829 | 6.996578 | 3.18376142 | ITPKA |
| A_32_P310335 | 1.17E-10 | 3.78E-13 | -8.974961 | 19.655288 | -2.31317882 | JAM2 |
| A_23_P429950 | 7.31E-10 | 3.29E-12 | -8.4550862 | 17.54723 | -2.86633444 | KAL1 |
| A_23_P131089 | 2.56E-13 | 2.48E-16 | -10.7668221 | 26.778546 | -2.61262484 | KANK3 |
| A_23_P67529 | 1.57E-03 | 1.09E-04 | 4.1104822 | 0.799492 | 2.06541771 | KCNN4 |
| A_23_P117852 | 1.69E-06 | 2.84E-08 | 6.272019 | 8.733892 | 2.35117 | KIAA0101 |
| A_23_P215048 | 1.89E-10 | 6.60E-13 | -8.8406018 | 19.111559 | -2.56871182 | KIAA0408 |
| A_23_P301521 | 4.98E-09 | 3.08E-11 | -7.9200703 | 15.37067 | -2.32352591 | KIAA1462 |
| A_23_P34788 | 9.79E-06 | 2.27E-07 | 5.7550573 | 6.719 | 2.4118121 | KIF2C |
| A_32_P62963 | 4.11E-04 | 2.10E-05 | 4.5727113 | 2.364955 | 2.43267321 | KRT16P2 |
| A_23_P2674 | 1.55E-05 | 3.94E-07 | -5.6157435 | 6.185404 | -2.29321812 | KRT4 |

| | | | | | | |
|---|---|---|---|---|---|---|
| A_23_P29773 | 3.99E-08 | 3.67E-10 | -7.3265091 | 12.959145 | -2.8222975 | LAMP3 |
| A_33_P3332406 | 1.14E-03 | 7.36E-05 | -4.2222758 | 1.169447 | -2.03712487 | LDLRAD1 |
| A_23_P69179 | 1.27E-04 | 5.00E-06 | -4.9588975 | 3.738639 | -2.24158048 | LEPREL1 |
| A_23_P210330 | 2.12E-07 | 2.50E-09 | -6.8644273 | 11.093967 | -2.0270567 | LGALSL |
| A_23_P21086 | 1.83E-16 | 4.34E-20 | -12.9935748 | 35.136823 | -2.50788557 | LGI3 |
| A_24_P90216 | 1.53E-08 | 1.15E-10 | 7.605211 | 14.09015 | 2.03630186 | LGR4 |
| A_23_P93169 | 4.03E-03 | 3.46E-04 | 3.7682689 | -0.295032 | 2.40774529 | LGSN |
| A_23_P88069 | 5.34E-09 | 3.35E-11 | -7.8995864 | 15.287316 | -2.07836101 | LHFP |
| A_32_P117354 | 8.72E-08 | 9.20E-10 | -7.1054306 | 12.064825 | -2.26359031 | LIMCH1 |
| A_33_P3268304 | 1.54E-08 | 1.16E-10 | -7.603004 | 14.081182 | -2.46008158 | LIMS2 |
| A_23_P53126 | 2.21E-07 | 2.63E-09 | -6.8520002 | 11.044044 | -2.06659219 | LMO2 |
| A_19_P00319646 | 9.36E-06 | 2.16E-07 | 5.7677944 | 6.768005 | 2.15696817 | LOC100499467 |
| A_21_P0014497 | 9.99E-12 | 2.15E-14 | -9.6680479 | 22.442663 | -2.98584932 | LOC100505976 |
| A_19_P00807053 | 3.06E-06 | 5.77E-08 | 6.0968775 | 8.045722 | 2.16704992 | LOC100506411 |
| A_21_P0008986 | 1.08E-15 | 4.37E-19 | -12.3836439 | 32.911134 | -2.20358106 | LOC100506542 |
| A_21_P0014482 | 7.20E-05 | 2.48E-06 | -5.1428618 | 4.411152 | -2.67000379 | LOC100507660 |
| A_33_P3283601 | 9.04E-12 | 1.89E-14 | -9.6991812 | 22.567054 | -2.78805468 | LOC389033 |
| A_19_P00322654 | 3.93E-15 | 1.86E-18 | -12.0070273 | 31.512039 | -2.25040749 | LOC400550 |

| | | | | | | |
|---|---|---|---|---|---|---|
| A_33_P3610138 | 8.51E-18 | 1.18E-21 | -13.9718121 | 38.600106 | -2.4310027 | LOC400568 |
| A_32_P60635 | 1.31E-04 | 5.18E-06 | -4.9495231 | 3.704665 | -2.47545422 | LOC643037 |
| A_23_P359214 | 1.61E-04 | 6.74E-06 | -4.8796079 | 3.452226 | -2.05884672 | LOC643650 |
| A_21_P0009343 | 1.54E-10 | 5.27E-13 | -8.8946905 | 19.330554 | -2.47614014 | LOC645722 |
| A_33_P3692756 | 7.77E-06 | 1.73E-07 | -5.8236747 | 6.983418 | -2.88698033 | LOC723809 |
| A_23_P146233 | 1.35E-05 | 3.37E-07 | -5.6555673 | 6.337477 | -2.54339787 | LPL |
| A_23_P118042 | 5.06E-12 | 8.90E-15 | -9.8832885 | 23.300988 | -4.0797319 | LRRC36 |
| A_33_P3369058 | 1.59E-05 | 4.07E-07 | -5.6080416 | 6.156037 | -2.47184689 | LRRK2 |
| A_23_P31376 | 1.76E-09 | 9.28E-12 | -8.2070903 | 16.538695 | -2.37617484 | LRRN3 |
| A_33_P3268622 | 2.22E-02 | 2.94E-03 | 3.0859313 | -2.2832 | 2.55809626 | LY6D |
| A_23_P397293 | 6.42E-02 | 1.16E-02 | 2.5954558 | -3.526625 | 2.43098569 | LY6K |
| A_32_P101031 | 2.08E-04 | 9.17E-06 | 4.7970219 | 3.15623 | 2.40375079 | LYPD1 |
| A_23_P150457 | 3.68E-12 | 5.84E-15 | -9.9860697 | 23.709437 | -2.16526175 | LYVE1 |
| A_23_P17134 | 3.34E-06 | 6.43E-08 | -6.0699336 | 7.940323 | -2.02760468 | MAL |
| A_23_P157914 | 7.52E-11 | 2.30E-13 | -9.0941185 | 20.136705 | -2.06418123 | MAMDC2 |
| A_23_P101992 | 1.73E-06 | 2.93E-08 | -6.2638402 | 8.701644 | -3.30185277 | MARCO |
| A_23_P116235 | 1.08E-07 | 1.17E-09 | 7.0482954 | 11.834234 | 2.62797595 | MDK |

| | | | | | | |
|---|---|---|---|---|---|---|
| A_23_P415021 | 7.83E-08 | 8.08E-10 | -7.1367749 | 12.191428 | -2.20149122 | METTL7A |
| A_32_P96036 | 1.46E-06 | 2.39E-08 | 6.3143286 | 8.900886 | 2.74051674 | MEX3A |
| A_23_P164057 | 3.95E-08 | 3.57E-10 | -7.3330282 | 12.98556 | -3.23748271 | MFAP4 |
| A_24_P245838 | 9.35E-10 | 4.42E-12 | -8.3847836 | 17.261451 | -3.10512632 | MGAT3 |
| A_23_P204286 | 5.14E-05 | 1.64E-06 | -5.25035 | 4.808967 | -2.02666636 | MGP |
| A_24_P260101 | 1.01E-09 | 4.83E-12 | -8.3630865 | 17.17323 | -2.46066541 | MME |
| A_23_P1691 | 1.44E-05 | 3.62E-07 | 5.6374194 | 6.26813 | 3.51642753 | MMP1 |
| A_23_P57417 | 7.51E-05 | 2.61E-06 | 5.1295257 | 4.362038 | 2.2617177 | MMP11 |
| A_23_P150316 | 3.28E-04 | 1.59E-05 | 4.6481088 | 2.628789 | 2.53202105 | MMP12 |
| A_23_P40174 | 1.01E-04 | 3.78E-06 | 5.0328752 | 4.007767 | 2.30310853 | MMP9 |
| A_33_P3212257 | 4.41E-09 | 2.69E-11 | -7.9522069 | 15.501448 | -2.44496796 | MMRN1 |
| A_32_P6015 | 2.68E-04 | 1.24E-05 | 4.7156502 | 2.866992 | 2.18330775 | MNX1 |
| A_23_P12746 | 1.11E-06 | 1.71E-08 | -6.3962423 | 9.224963 | -2.53648365 | MRC1 |
| A_33_P3312499 | 2.93E-06 | 5.45E-08 | -6.1110272 | 8.101125 | -3.6521564 | MS4A15 |
| A_23_P1904 | 3.68E-08 | 3.27E-10 | -7.3544259 | 13.072278 | -2.05532739 | MS4A2 |
| A_33_P3352098 | 7.31E-08 | 7.45E-10 | -7.1561873 | 12.269872 | -2.25362847 | MS4A7 |
| A_23_P139146 | 2.70E-04 | 1.25E-05 | -4.7125737 | 2.856104 | -2.22120853 | MS4A8B |
| A_24_P372223 | 3.91E-07 | 5.08E-09 | -6.6923452 | 10.404047 | -2.20999349 | MSR1 |
| A_24_P208825 | 1.10E-03 | 7.05E-05 | 4.2344294 | 1.210014 | 2.18404499 | MUC4 |

| | | | | | | |
|---|---|---|---|---|---|---|
| A_23_P23783 | 1.01E-15 | 3.81E-19 | -12.4198248 | 33.044556 | -3.31732949 | MYOC |
| A_33_P3423874 | 1.48E-11 | 3.40E-14 | -9.5570427 | 21.998519 | -3.02640345 | MYZAP |
| A_23_P360079 | 1.90E-09 | 1.02E-11 | -8.1844938 | 16.446753 | -2.58741155 | NCKAP5 |
| A_23_P110266 | 1.43E-05 | 3.59E-07 | -5.6393099 | 6.27535 | -2.06293712 | NDNF |
| A_23_P140748 | 2.96E-09 | 1.69E-11 | -8.063412 | 15.954018 | -2.42635821 | NDRG4 |
| A_23_P119835 | 7.94E-08 | 8.24E-10 | -7.1318655 | 12.171594 | -2.07058626 | NLRC4 |
| A_23_P69537 | 4.07E-03 | 3.50E-04 | 3.7648717 | -0.305596 | 2.08574785 | NMU |
| A_23_P209700 | 1.22E-13 | 1.11E-16 | -10.9668028 | 27.55423 | -2.12022587 | NMUR1 |
| A_23_P253896 | 7.89E-07 | 1.15E-08 | -6.4927374 | 9.607961 | -2.45180993 | NPNT |
| A_33_P3221408 | 3.11E-13 | 3.19E-16 | -10.7034799 | 26.531922 | -2.10688705 | NTNG1 |
| A_23_P82990 | 2.99E-11 | 7.91E-14 | -9.3520226 | 21.175761 | -3.16325998 | OGN |
| A_24_P124624 | 1.36E-06 | 2.18E-08 | -6.3363054 | 8.987735 | -2.13200444 | OLR1 |
| A_23_P257129 | 6.72E-02 | 1.23E-02 | 2.5716337 | -3.582632 | 2.27972202 | PAEP |
| A_32_P62997 | 1.10E-04 | 4.19E-06 | 5.0056386 | 3.908472 | 2.00037192 | PBK |
| A_23_P57709 | 6.46E-09 | 4.12E-11 | -7.8503657 | 15.087042 | -2.79473163 | PCOLCE2 |
| A_23_P109322 | 6.53E-03 | 6.32E-04 | 3.583965 | -0.858984 | 2.26932156 | PCP4 |

| | | | | | | |
|---|---|---|---|---|---|---|
| A_32_P142440 | 1.12E-05 | 2.70E-07 | -5.7111038 | 6.55017 | -2.56855359 | PCSK9 |
| A_33_P3378514 | 9.93E-11 | 3.14E-13 | -9.0194925 | 19.835295 | -2.42112032 | PDE5A |
| A_24_P243749 | 8.86E-07 | 1.32E-08 | -6.4593934 | 9.475471 | -2.49088947 | PDK4 |
| A_23_P7402 | 4.40E-08 | 4.13E-10 | -7.2978172 | 12.842913 | -2.23353251 | PDZD2 |
| A_33_P3627001 | 5.20E-08 | 5.00E-10 | -7.2522534 | 12.658427 | -3.57382749 | PEBP4 |
| A_23_P7965 | 1.40E-02 | 1.62E-03 | -3.2835762 | -1.73628 | -2.99502543 | PGC |
| A_23_P47614 | 1.28E-06 | 2.04E-08 | 6.3528296 | 9.053084 | 2.63966797 | PHLDA2 |
| A_33_P3215640 | 3.03E-10 | 1.17E-12 | -8.7023288 | 18.55112 | -3.53937535 | PI16 |
| A_23_P21485 | 3.95E-08 | 3.59E-10 | -7.3315438 | 12.979545 | -2.49329057 | PID1 |
| A_33_P3240328 | 4.20E-04 | 2.16E-05 | 4.565066 | 2.338327 | 2.8888747 | PITX1 |
| A_33_P3397443 | 1.43E-07 | 1.62E-09 | 6.9692865 | 11.515784 | 2.05698514 | PKMYT1 |
| A_24_P224116 | 4.75E-08 | 4.49E-10 | -7.2779703 | 12.762539 | -3.60765637 | PLA2G1B |
| A_33_P3252781 | 1.33E-12 | 1.89E-15 | -10.2630615 | 24.805313 | -3.02971838 | PLAC9 |
| A_23_P35414 | 6.15E-06 | 1.31E-07 | -5.8930838 | 7.251904 | -2.0168931 | PPP1R3C |
| A_33_P3330608 | 1.33E-06 | 2.13E-08 | -6.3428582 | 9.013645 | -2.08274836 | PRAM1 |
| A_21_P0000058 | 6.38E-08 | 6.29E-10 | 7.1968948 | 12.434448 | 2.06834767 | PROM2 |
| A_23_P259692 | 6.55E-06 | 1.42E-07 | 5.8727953 | 7.173321 | 2.36027234 | PSAT1 |
| A_33_P3235990 | 7.04E-12 | 1.38E-14 | -9.7766994 | 22.876427 | -2.23296416 | PTPN21 |

| | | | | | | |
|---|---|---|---|---|---|---|
| A_23_P53390 | 1.74E-09 | 9.16E-12 | -8.2100904 | 16.550902 | -2.00521538 | PTPRB |
| A_23_P101642 | 6.64E-04 | 3.75E-05 | 4.4122411 | 1.81103 | 2.0911737 | PTPRH |
| A_21_P0005949 | 1.51E-08 | 1.12E-10 | 7.6100724 | 14.109904 | 2.37584921 | PVT1 |
| A_33_P3387365 | 1.73E-09 | 9.02E-12 | -8.2137913 | 16.565959 | -2.06574892 | PXMP4 |
| A_23_P130194 | 3.96E-12 | 6.57E-15 | 9.9573671 | 23.59547 | 3.19881253 | PYCR1 |
| A_19_P00317023 | 9.89E-09 | 6.86E-11 | 7.7282872 | 14.590464 | 2.15654636 | Q6IHG2 |
| A_33_P3209229 | 1.46E-06 | 2.38E-08 | 6.3152291 | 8.904443 | 2.38728597 | RAB26 |
| A_23_P74115 | 4.30E-06 | 8.65E-08 | 5.9963797 | 7.653273 | 2.2116583 | RAD54L |
| A_23_P111860 | 6.19E-05 | 2.07E-06 | -5.1899694 | 4.585072 | -2.19690848 | RADIL |
| A_23_P111737 | 1.51E-11 | 3.49E-14 | -9.5504742 | 21.972207 | -2.33028824 | RAMP3 |
| A_23_P139600 | 3.75E-06 | 7.37E-08 | 6.0360773 | 7.808071 | 2.15506001 | RASAL1 |
| A_23_P75283 | 5.89E-05 | 1.94E-06 | -5.2075405 | 4.650115 | -2.42695912 | RBP4 |
| A_23_P83028 | 2.48E-11 | 6.47E-14 | -9.4009172 | 21.372253 | -2.05607299 | RECK |
| A_33_P3323847 | 2.19E-07 | 2.60E-09 | 6.8543267 | 11.053389 | 2.44446737 | RECQL4 |
| A_33_P3350863 | 2.35E-07 | 2.84E-09 | -6.8336232 | 10.970245 | -3.23770886 | RETN |
| A_33_P3387616 | 1.83E-07 | 2.13E-09 | 6.9026234 | 11.247504 | 2.15918484 | RHPN1 |
| A_23_P399255 | 1.74E-09 | 9.14E-12 | -8.2107686 | 16.553661 | -2.1233311 | RNF182 |
| A_23_P344421 | 3.21E-10 | 1.25E-12 | -8.6873323 | 18.49029 | -2.33032177 | ROBO4 |
| A_33_P3347528 | 9.69E-12 | 2.07E-14 | -9.6775219 | 22.480524 | -4.220475 | RP11-165H20.1 |
| A_23_P102950 | 1.72E-03 | 1.22E-04 | -4.0777478 | 0.692278 | -2.22553563 | RSPH1 |

| | | | | | | |
|---|---|---|---|---|---|---|
| A_33_P3251552 | 5.56E-07 | 7.69E-09 | -6.5914057 | 10.000865 | -2.08218382 | RSPO4 |
| A_24_P13041 | 4.36E-15 | 2.24E-18 | -11.9592311 | 31.333156 | -3.70831558 | RTKN2 |
| A_23_P58266 | 8.08E-03 | 8.20E-04 | 3.5024214 | -1.102408 | 2.91730207 | S100P |
| A_23_P94103 | 6.55E-08 | 6.51E-10 | -7.1885351 | 12.400642 | -2.99593841 | SCARA5 |
| A_23_P150583 | 1.65E-05 | 4.25E-07 | -5.5965703 | 6.112323 | -4.8580522 | SCGB1A1 |
| A_23_P145006 | 1.23E-02 | 1.38E-03 | -3.3358347 | -1.587553 | -2.12064649 | SCGB3A2 |
| A_23_P303833 | 7.95E-12 | 1.63E-14 | -9.7348589 | 22.709505 | -2.85598893 | SCN4B |
| A_23_P102331 | 9.63E-11 | 3.01E-13 | -9.0297348 | 19.876681 | -2.5658912 | SCN7A |
| A_23_P72668 | 2.24E-10 | 8.23E-13 | -8.7876736 | 18.897131 | -2.63373817 | SDPR |
| A_23_P86021 | 1.93E-03 | 1.41E-04 | -4.0357062 | 0.555337 | -2.02290171 | SELENBP1 |
| A_33_P3372727 | 4.74E-10 | 1.99E-12 | -8.5760179 | 18.038505 | -3.08761553 | SEMA5A |
| A_21_P0006968 | 5.49E-06 | 1.14E-07 | -5.9274904 | 7.385361 | -2.64920701 | SFTA1P |
| A_24_P223874 | 2.66E-06 | 4.86E-08 | -6.1391258 | 8.211249 | -3.82595821 | SFTPA1 |
| A_33_P3268919 | 1.68E-05 | 4.40E-07 | -5.588228 | 6.080554 | -4.14485851 | SFTPA2 |
| A_23_P95213 | 5.69E-03 | 5.33E-04 | -3.636692 | -0.699567 | -3.38069268 | SFTPC |
| A_32_P107617 | 2.02E-04 | 8.85E-06 | -4.8065847 | 3.19038 | -2.87076437 | SFTPD |
| A_33_P3386547 | 1.26E-05 | 3.10E-07 | 5.6769803 | 6.419399 | 2.00233223 | SGPP2 |

| | | | | | | |
|---|---|---|---|---|---|---|
| A_23_P48988 | 4.69E-13 | 5.37E-16 | -10.5741115 | 26.026865 | -2.23052424 | SH3GL3 |
| A_33_P3274562 | 2.03E-10 | 7.21E-13 | -8.8193091 | 19.025311 | -2.04071266 | SLC19A3 |
| A_23_P216468 | 1.49E-07 | 1.70E-09 | -6.9568893 | 11.465863 | -2.60960157 | SLC1A1 |
| A_23_P571 | 1.56E-05 | 3.98E-07 | 5.6136227 | 6.177316 | 2.56625979 | SLC2A1 |
| A_33_P3283824 | 4.70E-13 | 5.48E-16 | -10.5690118 | 26.006918 | -3.3590863 | SLC39A8 |
| A_23_P152995 | 5.00E-15 | 2.67E-18 | -11.9144674 | 31.165354 | -4.42912631 | SLC6A4 |
| A_33_P3242623 | 6.38E-03 | 6.13E-04 | 3.5934708 | -0.83036 | 2.15168823 | SLC7A11 |
| A_24_P335620 | 3.72E-04 | 1.85E-05 | 4.6073051 | 2.485731 | 2.20644574 | SLC7A5 |
| A_23_P144348 | 1.92E-10 | 6.76E-13 | -8.834767 | 19.087927 | -2.17091392 | SLIT2 |
| A_23_P58588 | 8.49E-06 | 1.91E-07 | -5.7980123 | 6.884408 | -2.26289888 | SLIT3 |
| A_24_P190472 | 1.95E-04 | 8.48E-06 | -4.8181017 | 3.231552 | -2.51686878 | SLPI |
| A_33_P3220470 | 2.15E-11 | 5.43E-14 | -9.4432125 | 21.542089 | -2.56460292 | SMAD6 |
| A_23_P145841 | 1.47E-07 | 1.67E-09 | -6.9609782 | 11.482327 | -2.66298819 | SOSTDC1 |
| A_23_P82775 | 2.72E-08 | 2.29E-10 | -7.4397175 | 13.418162 | -2.29965111 | SOX17 |
| A_23_P89509 | 2.40E-07 | 2.90E-09 | 6.8280923 | 10.948041 | 2.35526734 | SPAG5 |
| A_23_P113351 | 6.40E-08 | 6.33E-10 | -7.1955402 | 12.42897 | -2.24115941 | SPARCL1 |
| A_23_P214079 | 5.93E-05 | 1.95E-06 | 5.2057553 | 4.643503 | 4.79181845 | SPINK1 |
| A_33_P3214948 | 2.54E-12 | 3.92E-15 | -10.0838528 | 24.097131 | -3.0384658 | SPOCK2 |

| | | | | | | |
|---|---|---|---|---|---|---|
| A_23_P7313 | 1.62E-08 | 1.23E-10 | 7.5877269 | 14.01911 | 4.20401307 | SPP1 |
| A_23_P96383 | 8.36E-07 | 1.24E-08 | -6.4758748 | 9.54094 | -2.01110847 | SRPX |
| A_24_P244706 | 1.11E-07 | 1.21E-09 | -7.0384578 | 11.794556 | -2.25852019 | SSTR1 |
| A_33_P3307495 | 6.96E-04 | 3.97E-05 | 4.3964053 | 1.756946 | 2.53953987 | STRA6 |
| A_23_P168556 | 4.62E-08 | 4.35E-10 | 7.2855182 | 12.793103 | 2.27977942 | STX1A |
| A_23_P205713 | 6.41E-08 | 6.35E-10 | -7.1946857 | 12.425514 | -2.40896909 | STXBP6 |
| A_23_P314101 | 6.07E-04 | 3.35E-05 | -4.4431808 | 1.917003 | -2.82058652 | SUSD2 |
| A_23_P216596 | 1.39E-11 | 3.14E-14 | -9.5763947 | 22.076018 | -2.28829219 | SVEP1 |
| A_23_P421306 | 1.55E-03 | 1.07E-04 | 4.1160907 | 0.817912 | 2.57013644 | SYT12 |
| A_33_P3239185 | 3.51E-05 | 1.04E-06 | 5.3677012 | 5.247124 | 2.43972207 | SYT7 |
| A_23_P164451 | 6.57E-08 | 6.55E-10 | -7.1870592 | 12.394674 | -2.24764394 | TBX2 |
| A_23_P156890 | 8.50E-14 | 7.06E-17 | -11.0807637 | 27.994217 | -2.78731144 | TCF21 |
| A_23_P374695 | 5.71E-10 | 2.47E-12 | -8.5237914 | 17.826392 | -2.93349731 | TEK |
| A_24_P322771 | 1.35E-01 | 3.15E-02 | 2.1960366 | -4.408201 | 2.11166015 | TFF1 |
| A_23_P91390 | 5.09E-07 | 6.90E-09 | -6.6178012 | 10.106181 | -2.13355328 | THBD |
| A_23_P399078 | 1.05E-09 | 5.11E-12 | -8.3499317 | 17.119738 | -2.38188549 | TIMP3 |
| A_23_P107421 | 2.57E-06 | 4.65E-08 | 6.150293 | 8.255053 | 2.43515271 | TK1 |
| A_23_P15450 | 5.73E-18 | 6.79E-22 | -14.1237412 | 39.126079 | -4.46650537 | TMEM100 |

| | | | | | | |
|---|---|---|---|---|---|---|
| A_23_P107775 | 5.04E-04 | 2.68E-05 | -4.5051453 | 2.130439 | -2.85382762 | TMEM190 |
| A_33_P3406661 | 4.60E-04 | 2.41E-05 | 4.5350163 | 2.233892 | 2.19606983 | TMEM63C |
| A_33_P3296846 | 7.91E-05 | 2.80E-06 | 5.1118157 | 4.296902 | 2.68173095 | TMPRSS4 |
| A_23_P166823 | 8.13E-09 | 5.37E-11 | -7.7871861 | 14.830015 | -3.3995383 | TNNC1 |
| A_33_P3209491 | 3.87E-09 | 2.31E-11 | -7.9888211 | 15.650453 | -2.20627702 | TNS1 |
| A_23_P207850 | 2.58E-02 | 3.55E-03 | 3.0210655 | -2.457167 | 2.13632053 | TNS4 |
| A_23_P156708 | 3.78E-08 | 3.38E-10 | -7.3461285 | 13.038649 | -3.06070129 | TNXB |
| A_23_P26386 | 1.09E-05 | 2.63E-07 | -5.7184982 | 6.578542 | -2.68935813 | TPPP3 |
| A_23_P37702 | 1.94E-05 | 5.22E-07 | -5.5446443 | 5.914853 | -2.04028515 | TPSAB1 |
| A_23_P68610 | 5.80E-05 | 1.90E-06 | 5.2123212 | 4.667828 | 2.18603549 | TPX2 |
| A_23_P150935 | 1.72E-06 | 2.89E-08 | 6.2672484 | 8.715081 | 2.39758279 | TROAP |
| A_23_P85269 | 7.53E-09 | 4.91E-11 | -7.8083676 | 14.916179 | -2.39203926 | TTN |
| A_23_P77493 | 1.06E-09 | 5.23E-12 | 8.3441557 | 17.096249 | 4.68610019 | TUBB3 |
| A_24_P297539 | 4.07E-06 | 8.08E-08 | 6.0133266 | 7.71932 | 3.05493415 | UBE2C |
| A_23_P115482 | 1.37E-09 | 7.01E-12 | 8.2742106 | 16.811763 | 2.64967501 | UBE2T |
| A_23_P208880 | 3.40E-09 | 1.98E-11 | 8.026066 | 15.80203 | 2.21448629 | UHRF1 |
| A_24_P112160 | 4.87E-10 | 2.07E-12 | -8.5665357 | 18 | -2.18327738 | UPK3B |
| A_23_P380208 | 3.30E-07 | 4.13E-09 | -6.7427591 | 10.605848 | -2.81158138 | VEPH1 |
| A_33_P3276703 | 4.76E-03 | 4.25E-04 | 3.7063094 | -0.486707 | 2.01898647 | VGF |

| | | | | | | |
|---|---|---|---|---|---|---|
| A_23_P132763 | 7.58E-12 | 1.53E-14 | -9.75122 | 22.774795 | -2.38643837 | VGLL3 |
| A_24_P105933 | 7.58E-13 | 9.89E-16 | -10.4231085 | 25.435092 | -2.95091425 | VIPR1 |
| A_23_P36018 | 4.18E-05 | 1.29E-06 | -5.3136691 | 5.0449 | -2.44861975 | VSIG2 |
| A_23_P217269 | 1.28E-06 | 2.03E-08 | -6.3537371 | 9.056674 | -2.46668511 | VSIG4 |
| A_23_P105562 | 3.50E-07 | 4.42E-09 | -6.7261254 | 10.539235 | -2.11612799 | VWF |
| A_32_P216520 | 3.16E-06 | 5.99E-08 | -6.087494 | 8.009001 | -3.66701015 | WIF1 |
| A_23_P102611 | 1.17E-06 | 1.83E-08 | -6.380311 | 9.161856 | -2.62524308 | WISP2 |
| A_23_P385690 | 6.02E-16 | 2.02E-19 | -12.5857904 | 33.65434 | -3.52023585 | WNT3A |
| A_24_P271696 | 5.04E-02 | 8.42E-03 | 2.7136951 | -3.242447 | 3.32459048 | XAGE1A |
| A_33_P3270863 | 1.92E-04 | 8.34E-06 | 4.8226074 | 3.247672 | 2.29180712 | XDH |
| A_21_P0002189 | 1.25E-04 | 4.88E-06 | 4.9652 | 3.761497 | 2.08888727 | XLOC_001406 |
| A_19_P00320998 | 5.88E-12 | 1.08E-14 | -9.8355886 | 23.111113 | -2.77606362 | XLOC_001412 |
| A_21_P0001778 | 3.87E-09 | 2.31E-11 | -7.9890606 | 15.651428 | -4.13182336 | XLOC_001575 |
| A_21_P0006273 | 8.70E-07 | 1.29E-08 | -6.4650123 | 9.497787 | -2.24157698 | XLOC_007724 |
| A_21_P0006969 | 5.96E-06 | 1.26E-07 | -5.9030708 | 7.290617 | -2.44389648 | XLOC_008730 |
| A_21_P0009008 | 1.67E-08 | 1.28E-10 | -7.5782175 | 13.980477 | -2.19296276 | XLOC_011872 |
| A_21_P0009734 | 6.16E-11 | 1.75E-13 | -9.1596836 | 20.401253 | -2.2027974 | XLOC_013194 |

| | | | | | | |
|---|---|---|---|---|---|---|
| A_21_P0009825 | 8.75E-09 | 5.91E-11 | -7.764075 | 14.736011 | -2.78171343 | XLOC_013457 |
| A_21_P0011969 | 3.96E-18 | 3.92E-22 | -14.276924 | 39.653141 | -2.08058077 | XLOC_l2_008275 |
| A_23_P113793 | 1.87E-05 | 4.98E-07 | -5.556405 | 5.95952 | -2.0420415 | ZBED2 |

**Supplementary Table S3:**

Details of protein-protein interaction network in LUAD

| node1 | node2 | homology | coexpression | combined_score |
|---|---|---|---|---|
| ANGPT1 | TEK | 0 | 0.076 | 0.999 |
| CCNB2 | CCNB1 | 0.943 | 0.955 | 0.999 |
| CDC45 | GINS1 | 0 | 0.651 | 0.998 |
| UBE2C | CCNB1 | 0 | 0.924 | 0.998 |
| NPNT | ITGA8 | 0 | 0.062 | 0.998 |
| PKMYT1 | CCNB1 | 0 | 0.405 | 0.997 |
| ALOX5AP | ALOX5 | 0 | 0.32 | 0.997 |
| KIF2C | CCNB2 | 0 | 0.914 | 0.996 |
| KIF2C | CCNB1 | 0 | 0.929 | 0.996 |
| BIRC5 | CCNB1 | 0 | 0.888 | 0.996 |
| CALCRL | RAMP3 | 0 | 0.061 | 0.996 |
| CD19 | CR2 | 0 | 0.16 | 0.996 |
| BIRC5 | CCNB2 | 0 | 0.88 | 0.995 |
| CENPF | CCNB2 | 0 | 0.901 | 0.995 |
| CCNB2 | PKMYT1 | 0 | 0.607 | 0.995 |
| HBB | HBA2 | 0.9 | 0.57 | 0.995 |
| KIF2C | CENPF | 0 | 0.884 | 0.995 |
| GPIHBP1 | LPL | 0 | 0.147 | 0.993 |
| UBE2C | CCNB2 | 0 | 0.976 | 0.993 |
| NMUR1 | NMU | 0 | 0 | 0.993 |
| CENPF | CCNB1 | 0 | 0.849 | 0.993 |
| SFTPA1 | SFTPC | 0 | 0.533 | 0.992 |
| KIF2C | BIRC5 | 0 | 0.872 | 0.992 |
| SFTPA2 | SFTPC | 0 | 0.569 | 0.992 |
| BIRC5 | CDCA5 | 0 | 0.881 | 0.991 |
| CCNB2 | CDCA5 | 0 | 0.837 | 0.99 |
| CENPF | BIRC5 | 0 | 0.809 | 0.99 |
| HBD | HBA2 | 0.9 | 0.292 | 0.989 |
| LGR4 | RSPO4 | 0 | 0 | 0.988 |

| | | | | |
|---|---|---|---|---|
| CDC45 | CDT1 | 0 | 0.852 | 0.987 |
| STRA6 | RBP4 | 0 | 0 | 0.986 |
| SFTPD | SFTPC | 0 | 0.334 | 0.985 |
| UBE2C | BIRC5 | 0 | 0.956 | 0.983 |
| CENPF | CDCA5 | 0 | 0.714 | 0.982 |
| ADH1B | ADH1A | 0.985 | 0.828 | 0.982 |
| CDCA5 | CCNB1 | 0 | 0.741 | 0.981 |
| CAV1 | CAV2 | 0.858 | 0.819 | 0.981 |
| ASPM | CENPF | 0 | 0.962 | 0.98 |
| HHIP | IHH | 0 | 0 | 0.98 |
| LPL | FABP4 | 0 | 0.186 | 0.98 |
| KIF2C | CDCA5 | 0 | 0.67 | 0.979 |
| HIST2H3A | HIST1H2AI | 0 | 0.466 | 0.978 |
| UBE2T | UBE2C | 0.815 | 0.733 | 0.978 |
| KIF2C | TPX2 | 0 | 0.942 | 0.977 |
| GPC3 | GOLM1 | 0 | 0 | 0.976 |
| UBE2C | TPX2 | 0 | 0.95 | 0.976 |
| GRK5 | ADRB2 | 0 | 0 | 0.975 |
| ADH1C | ADH1B | 0.986 | 0.179 | 0.973 |
| TPX2 | CCNB1 | 0 | 0.946 | 0.972 |
| ASPM | CCNB1 | 0 | 0.95 | 0.97 |
| MMP9 | TIMP3 | 0 | 0 | 0.969 |
| ASPM | TPX2 | 0 | 0.966 | 0.968 |
| SFTPA1 | SFTPA2 | 0.986 | 0.522 | 0.968 |
| CEP55 | CENPF | 0 | 0.934 | 0.967 |
| COL1A1 | DCN | 0 | 0.868 | 0.966 |
| WIF1 | WNT3A | 0 | 0.05 | 0.965 |
| CD36 | FABP4 | 0 | 0.267 | 0.964 |
| SLPI | CAMP | 0 | 0.088 | 0.964 |
| VIPR1 | RAMP3 | 0 | 0.079 | 0.963 |
| BIRC5 | TPX2 | 0 | 0.932 | 0.962 |
| MMP9 | MMP1 | 0.75 | 0.518 | 0.961 |
| SPARCL1 | SPP1 | 0 | 0.11 | 0.961 |
| TPX2 | CCNB2 | 0 | 0.935 | 0.961 |
| TTN | TNNC1 | 0 | 0.143 | 0.96 |
| F11 | F12 | 0.643 | 0.396 | 0.959 |
| EDN1 | ADRB2 | 0 | 0 | 0.958 |
| CEP55 | UBE2C | 0 | 0.911 | 0.957 |
| SFTPA1 | SFTPD | 0.809 | 0.307 | 0.956 |
| MFAP4 | FBLN5 | 0 | 0.217 | 0.956 |
| KIF2C | UBE2C | 0 | 0.919 | 0.955 |
| KIF2C | SPAG5 | 0 | 0.919 | 0.955 |

| | | | | |
|---|---|---|---|---|
| HBB | CAMP | 0 | 0.076 | 0.955 |
| CEP55 | CCNB1 | 0 | 0.867 | 0.954 |
| HS6ST2 | GPC3 | 0 | 0.11 | 0.953 |
| AGER | S100P | 0 | 0 | 0.953 |
| A2M | VWF | 0 | 0.213 | 0.952 |
| CEP55 | ASPM | 0 | 0.932 | 0.952 |
| SLIT2 | ROBO4 | 0 | 0.079 | 0.951 |
| EDN1 | MMP9 | 0 | 0 | 0.95 |
| TTN | DES | 0 | 0.245 | 0.95 |
| SPAG5 | CCNB2 | 0 | 0.928 | 0.95 |
| DUOX1 | DUOXA1 | 0 | 0.207 | 0.949 |
| KIF2C | CEP55 | 0 | 0.905 | 0.946 |
| ASPM | CCNB2 | 0 | 0.909 | 0.946 |
| UBE2C | KIAA0101 | 0 | 0.937 | 0.945 |
| CENPF | TPX2 | 0 | 0.891 | 0.944 |
| GPC3 | DCN | 0 | 0.061 | 0.943 |
| KIAA0101 | CCNB2 | 0 | 0.936 | 0.943 |
| SPARCL1 | GPC3 | 0 | 0.063 | 0.943 |
| KIF2C | TROAP | 0 | 0.928 | 0.943 |
| SFTPA2 | SFTPD | 0.825 | 0.34 | 0.942 |
| KIF2C | ASPM | 0 | 0.915 | 0.942 |
| SCN7A | SCN4B | 0 | 0.061 | 0.942 |
| ACVRL1 | CAV1 | 0 | 0.062 | 0.942 |
| CX3CR1 | CXCL13 | 0 | 0 | 0.939 |
| MMRN1 | VWF | 0 | 0.149 | 0.939 |
| DLC1 | TNS1 | 0 | 0.108 | 0.939 |
| TK1 | BIRC5 | 0 | 0.892 | 0.939 |
| UBE2C | SPAG5 | 0 | 0.924 | 0.938 |
| CACNA2D2 | CACNG6 | 0 | 0 | 0.938 |
| CEP55 | SPAG5 | 0 | 0.913 | 0.938 |
| RBP4 | CES1 | 0 | 0.154 | 0.938 |
| CTHRC1 | WNT3A | 0 | 0 | 0.938 |
| B3GNT3 | FUT3 | 0 | 0.203 | 0.938 |
| SPP1 | GPC3 | 0 | 0.069 | 0.938 |
| CDC45 | CCNB2 | 0 | 0.864 | 0.937 |
| MMP9 | CAMP | 0 | 0.14 | 0.937 |
| CD36 | OLR1 | 0 | 0 | 0.936 |
| MARCO | SCGB3A2 | 0 | 0 | 0.935 |
| SPAG5 | BIRC5 | 0 | 0.903 | 0.934 |
| SPAG5 | TROAP | 0 | 0.894 | 0.932 |
| MMP9 | A2M | 0 | 0 | 0.932 |
| CAV1 | WNT3A | 0 | 0 | 0.932 |

| | | | | |
|---|---|---|---|---|
| ASPM | SPAG5 | 0 | 0.916 | 0.932 |
| A2M | MMP1 | 0 | 0 | 0.932 |
| FIGF | VWF | 0 | 0 | 0.932 |
| COL10A1 | COL1A1 | 0.752 | 0.109 | 0.931 |
| ADRB1 | ADRB2 | 0.937 | 0 | 0.93 |
| MUC4 | B3GNT3 | 0 | 0.119 | 0.928 |
| ASPM | UBE2C | 0 | 0.898 | 0.928 |
| PCOLCE2 | COL1A1 | 0 | 0.083 | 0.927 |
| CALCRL | ADRB2 | 0 | 0 | 0.926 |
| SPP1 | GOLM1 | 0 | 0 | 0.926 |
| CAMP | RETN | 0 | 0.215 | 0.926 |
| GRK5 | GNG11 | 0 | 0 | 0.924 |
| DES | TNNC1 | 0 | 0.119 | 0.924 |
| CDT1 | CCNB1 | 0 | 0.746 | 0.924 |
| CALCRL | GNG11 | 0 | 0.062 | 0.924 |
| HBD | HBB | 0.983 | 0.27 | 0.924 |
| GRK5 | EDN1 | 0 | 0 | 0.923 |
| CENPF | UBE2C | 0 | 0.85 | 0.923 |
| CENPF | SPAG5 | 0 | 0.886 | 0.923 |
| RECQL4 | CDC45 | 0 | 0.407 | 0.922 |
| HJURP | UBE2C | 0 | 0.89 | 0.922 |
| MUC4 | GALNT14 | 0 | 0 | 0.921 |
| LYPD1 | LY6D | 0 | 0 | 0.92 |
| HJURP | CENPF | 0 | 0.893 | 0.92 |
| CDT1 | CCNB2 | 0 | 0.867 | 0.92 |
| GRK5 | NMUR1 | 0 | 0 | 0.92 |
| LEPREL1 | COL1A1 | 0 | 0.064 | 0.919 |
| GPX3 | ALOX5 | 0 | 0.061 | 0.919 |
| CLDN5 | CLDN18 | 0.755 | 0.061 | 0.918 |
| CALCRL | ADRB1 | 0 | 0 | 0.918 |
| UHRF1 | CDCA7 | 0 | 0.897 | 0.917 |
| SPARCL1 | CHRDL1 | 0 | 0.138 | 0.917 |
| HJURP | KIF2C | 0 | 0.888 | 0.917 |
| FABP5 | RETN | 0 | 0.062 | 0.917 |
| ASPM | BIRC5 | 0 | 0.892 | 0.916 |
| SPP1 | ITGA8 | 0 | 0.055 | 0.916 |
| HJURP | CCNB2 | 0 | 0.9 | 0.916 |
| ADH1C | ADH1A | 0.985 | 0.182 | 0.916 |
| SLIT3 | ROBO4 | 0 | 0.096 | 0.915 |
| CFD | VWF | 0 | 0.077 | 0.915 |
| LGI3 | STX1A | 0 | 0.136 | 0.915 |
| GRK5 | CHRM1 | 0 | 0 | 0.914 |

| | | | | |
|---|---|---|---|---|
| HJURP | TROAP | 0 | 0.881 | 0.914 |
| MMP12 | A2M | 0 | 0.074 | 0.914 |
| CEP55 | CCNB2 | 0 | 0.852 | 0.914 |
| MMP1 | MMP11 | 0.81 | 0 | 0.913 |
| MMRN1 | A2M | 0 | 0 | 0.913 |
| COL6A6 | COL1A1 | 0.57 | 0.061 | 0.913 |
| CLEC3B | TIMP3 | 0 | 0.095 | 0.913 |
| CHRM1 | ADRB2 | 0.681 | 0.05 | 0.913 |
| VIPR1 | GNG11 | 0 | 0 | 0.913 |
| EDN1 | NMU | 0 | 0 | 0.913 |
| CFD | A2M | 0 | 0 | 0.913 |
| NMUR1 | GNG11 | 0 | 0 | 0.912 |
| EDN1 | NMUR1 | 0 | 0 | 0.912 |
| CD52 | CEACAM5 | 0 | 0 | 0.912 |
| ADRB1 | GNG11 | 0 | 0 | 0.912 |
| SSTR1 | GNG11 | 0 | 0 | 0.912 |
| ADRB2 | GNG11 | 0 | 0 | 0.912 |
| BIRC5 | KIAA0101 | 0 | 0.893 | 0.911 |
| GPC3 | IHH | 0 | 0.052 | 0.911 |
| CALCRL | VIPR1 | 0.763 | 0 | 0.911 |
| SPP1 | VGF | 0 | 0.069 | 0.911 |
| KIAA0101 | CCNB1 | 0 | 0.892 | 0.911 |
| COL6A6 | COL10A1 | 0.677 | 0.062 | 0.91 |
| MMP9 | HBB | 0 | 0.054 | 0.91 |
| ADRB2 | RAMP3 | 0 | 0 | 0.91 |
| SPAG5 | KIAA0101 | 0 | 0.909 | 0.909 |
| CDH13 | CDH3 | 0.802 | 0 | 0.908 |
| CHRM1 | NMU | 0 | 0 | 0.908 |
| NMUR1 | SSTR1 | 0.673 | 0.063 | 0.907 |
| PCSK9 | VGF | 0 | 0 | 0.907 |
| GPC3 | CHRDL1 | 0 | 0.061 | 0.907 |
| FIGF | COL1A1 | 0 | 0 | 0.907 |
| HJURP | BIRC5 | 0 | 0.887 | 0.907 |
| SLIT2 | SLIT3 | 0.973 | 0.097 | 0.907 |
| GPC3 | PCSK9 | 0 | 0.076 | 0.907 |
| MSR1 | COL1A1 | 0.595 | 0 | 0.906 |
| CD36 | MCEMP1 | 0 | 0.102 | 0.906 |
| ASPM | KIAA0101 | 0 | 0.903 | 0.906 |
| SPAG5 | TPX2 | 0 | 0.878 | 0.906 |
| FABP5 | FABP4 | 0.945 | 0.069 | 0.906 |
| HJURP | ASPM | 0 | 0.893 | 0.906 |
| GPIHBP1 | LY6D | 0 | 0.058 | 0.906 |

| | | | | |
|---|---|---|---|---|
| A2M | FIGF | 0 | 0 | 0.906 |
| MMRN1 | CFD | 0 | 0.063 | 0.906 |
| GPC3 | WNT3A | 0 | 0 | 0.906 |
| MCEMP1 | OLR1 | 0 | 0.091 | 0.905 |
| EDN1 | CHRM1 | 0 | 0 | 0.905 |
| ENTPD8 | TK1 | 0 | 0.063 | 0.905 |
| SPP1 | PCSK9 | 0 | 0 | 0.905 |
| MMRN1 | FIGF | 0 | 0.061 | 0.905 |
| ADAMTSL3 | ADAMTS8 | 0.596 | 0.061 | 0.904 |
| RECK | NTNG1 | 0 | 0.062 | 0.904 |
| CACNA2D2 | STX1A | 0 | 0.049 | 0.904 |
| CHRM1 | NMUR1 | 0 | 0.049 | 0.904 |
| RNF182 | UBE2C | 0 | 0 | 0.904 |
| SLPI | RETN | 0 | 0 | 0.903 |
| MFAP4 | SFTPD | 0 | 0 | 0.903 |
| SBSPON | ADAMTSL3 | 0 | 0.073 | 0.903 |
| CEP55 | BIRC5 | 0 | 0.834 | 0.903 |
| SH3GL3 | ADRB2 | 0 | 0 | 0.903 |
| RECK | CEACAM5 | 0 | 0 | 0.903 |
| METTL7A | HBB | 0 | 0.076 | 0.903 |
| A2M | MMP11 | 0 | 0 | 0.903 |
| CD36 | PTPRB | 0 | 0.062 | 0.903 |
| SPARCL1 | VGF | 0 | 0.062 | 0.902 |
| VIPR1 | ADRB2 | 0 | 0 | 0.902 |
| LYPD1 | NTNG1 | 0 | 0.061 | 0.902 |
| CX3CR1 | GNG11 | 0 | 0 | 0.902 |
| PLAC8 | FABP5 | 0 | 0.063 | 0.902 |
| PLAC8 | RETN | 0 | 0.069 | 0.902 |
| EDN1 | GNG11 | 0 | 0.065 | 0.902 |
| CHRM1 | GNG11 | 0 | 0.061 | 0.902 |
| CX3CR1 | NMUR1 | 0.599 | 0 | 0.902 |
| PTPRB | OLR1 | 0 | 0.062 | 0.902 |
| GPIHBP1 | NTNG1 | 0 | 0.061 | 0.902 |
| ADRB1 | VIPR1 | 0 | 0.061 | 0.902 |
| DNAI2 | DYNLRB2 | 0 | 0.063 | 0.902 |
| GRK5 | WNT3A | 0 | 0 | 0.901 |
| SSTR1 | NMU | 0 | 0 | 0.901 |
| GPIHBP1 | LYPD1 | 0 | 0 | 0.901 |
| SPARCL1 | GOLM1 | 0 | 0 | 0.901 |
| TPSAB1 | MMP1 | 0 | 0 | 0.901 |
| LYPD1 | CEACAM5 | 0 | 0 | 0.901 |
| GPER1 | NMU | 0 | 0 | 0.901 |

| | | | | |
|---|---|---|---|---|
| GPER1 | SSTR1 | 0.649 | 0 | 0.901 |
| CFD | FIGF | 0 | 0 | 0.901 |
| B3GNT3 | OGN | 0 | 0.053 | 0.901 |
| GPIHBP1 | RECK | 0 | 0 | 0.9 |
| CXCL13 | SSTR1 | 0 | 0 | 0.9 |
| SEMA5A | SBSPON | 0 | 0 | 0.9 |
| GPIHBP1 | CD52 | 0 | 0 | 0.9 |
| CHRDL1 | PCSK9 | 0 | 0 | 0.9 |
| COL6A6 | LEPREL1 | 0 | 0 | 0.9 |
| CXCL13 | NMU | 0 | 0 | 0.9 |
| LYPD1 | CD52 | 0 | 0 | 0.9 |
| NMU | GNG11 | 0 | 0 | 0.9 |
| LY6D | CEACAM5 | 0 | 0 | 0.9 |
| GPER1 | CXCL13 | 0 | 0 | 0.9 |
| COL10A1 | LEPREL1 | 0 | 0 | 0.9 |
| METTL7A | CAMP | 0 | 0 | 0.9 |
| CACNA2D2 | GNG11 | 0 | 0.049 | 0.9 |
| CD52 | LY6D | 0 | 0 | 0.9 |
| SPAG5 | CCNB1 | 0 | 0.872 | 0.9 |
| NTNG1 | CEACAM5 | 0 | 0 | 0.9 |
| NMUR1 | CXCL13 | 0 | 0 | 0.9 |
| GPC3 | VGF | 0 | 0 | 0.9 |
| NTNG1 | LY6D | 0 | 0 | 0.9 |
| LYPD1 | RECK | 0 | 0 | 0.9 |
| GOLM1 | PCSK9 | 0 | 0 | 0.9 |
| CXCL13 | GNG11 | 0 | 0 | 0.9 |
| GPER1 | NMUR1 | 0.594 | 0 | 0.9 |
| GNG11 | RAMP3 | 0 | 0 | 0.9 |
| SPARCL1 | PCSK9 | 0 | 0 | 0.9 |
| GPER1 | CX3CR1 | 0.704 | 0 | 0.9 |
| GPER1 | GNG11 | 0 | 0 | 0.9 |
| ADRB1 | RAMP3 | 0 | 0 | 0.9 |
| SEMA5A | ADAMTS8 | 0.544 | 0 | 0.9 |
| GPIHBP1 | CEACAM5 | 0 | 0 | 0.9 |
| GRK5 | NMU | 0 | 0 | 0.9 |
| CHRDL1 | VGF | 0 | 0 | 0.9 |
| CX3CR1 | NMU | 0 | 0 | 0.9 |
| SEMA5A | ADAMTSL3 | 0 | 0 | 0.9 |
| CD52 | NTNG1 | 0 | 0 | 0.9 |
| CX3CR1 | SSTR1 | 0.687 | 0 | 0.9 |
| SPP1 | CHRDL1 | 0 | 0 | 0.9 |
| PTPRB | MCEMP1 | 0 | 0 | 0.9 |

| | | | | |
|---|---|---|---|---|
| SBSPON | ADAMTS8 | 0 | 0 | 0.9 |
| GOLM1 | CHRDL1 | 0 | 0 | 0.9 |
| RECK | CD52 | 0 | 0 | 0.9 |
| RECK | LY6D | 0 | 0.049 | 0.9 |
| METTL7A | MMP9 | 0 | 0 | 0.9 |
| GOLM1 | VGF | 0 | 0 | 0.9 |

**Supplementary Table S4:**

Interactions of miRNA-mRNA regulatory network in LUAD

| Node | Target |
|---|---|
| A2M | VWF |
| A2M | MMP1 |
| A2M | FIGF |
| A2M | MMP11 |
| ACVRL1 | CAV1 |
| ADAMTSL3 | ADAMTS8 |
| ADH1B | ADH1A |
| ADH1C | ADH1B |
| ADH1C | ADH1A |
| ADRB1 | ADRB2 |
| ADRB1 | GNG11 |
| ADRB1 | VIPR1 |
| ADRB1 | RAMP3 |
| ADRB2 | GNG11 |
| ADRB2 | RAMP3 |
| AGER | S100P |
| ALOX5AP | ALOX5 |
| ANGPT1 | TEK |
| ASPM | CENPF |
| ASPM | CCNB1 |
| ASPM | TPX2 |
| ASPM | CCNB2 |
| ASPM | SPAG5 |
| ASPM | UBE2C |
| ASPM | BIRC5 |
| ASPM | KIAA0101 |
| B3GNT3 | FUT3 |
| B3GNT3 | OGN |
| BIRC5 | CCNB1 |
| BIRC5 | CCNB2 |

| | |
|---|---|
| BIRC5 | CDCA5 |
| BIRC5 | TPX2 |
| BIRC5 | KIAA0101 |
| CACNA2D2 | CACNG6 |
| CACNA2D2 | STX1A |
| CACNA2D2 | GNG11 |
| CALCRL | RAMP3 |
| CALCRL | ADRB2 |
| CALCRL | GNG11 |
| CALCRL | ADRB1 |
| CALCRL | VIPR1 |
| CAMP | RETN |
| CAV1 | CAV2 |
| CAV1 | WNT3A |
| CCNB2 | CCNB1 |
| CCNB2 | PKMYT1 |
| CCNB2 | CDCA5 |
| CD19 | CR2 |
| CD36 | FABP4 |
| CD36 | OLR1 |
| CD36 | MCEMP1 |
| CD36 | PTPRB |
| CD52 | CEACAM5 |
| CD52 | LY6D |
| CD52 | NTNG1 |
| CDC45 | GINS1 |
| CDC45 | CDT1 |
| CDC45 | CCNB2 |
| CDCA5 | CCNB1 |
| CDH13 | CDH3 |
| CDT1 | CCNB1 |
| CDT1 | CCNB2 |
| CENPF | CCNB2 |
| CENPF | CCNB1 |
| CENPF | BIRC5 |
| CENPF | CDCA5 |
| CENPF | TPX2 |
| CENPF | UBE2C |
| CENPF | SPAG5 |
| CEP55 | CENPF |
| CEP55 | UBE2C |
| CEP55 | CCNB1 |

| | |
|---|---|
| CEP55 | ASPM |
| CEP55 | SPAG5 |
| CEP55 | CCNB2 |
| CEP55 | BIRC5 |
| CFD | VWF |
| CFD | A2M |
| CFD | FIGF |
| CHRDL1 | PCSK9 |
| CHRDL1 | VGF |
| CHRM1 | ADRB2 |
| CHRM1 | NMU |
| CHRM1 | NMUR1 |
| CHRM1 | GNG11 |
| CLDN5 | CLDN18 |
| CLEC3B | TIMP3 |
| COL10A1 | COL1A1 |
| COL10A1 | LEPREL1 |
| COL1A1 | DCN |
| COL6A6 | COL1A1 |
| COL6A6 | COL10A1 |
| COL6A6 | LEPREL1 |
| CTHRC1 | WNT3A |
| CX3CR1 | CXCL13 |
| CX3CR1 | GNG11 |
| CX3CR1 | NMUR1 |
| CX3CR1 | NMU |
| CX3CR1 | SSTR1 |
| CXCL13 | SSTR1 |
| CXCL13 | NMU |
| CXCL13 | GNG11 |
| DES | TNNC1 |
| DLC1 | TNS1 |
| DNAI2 | DYNLRB2 |
| DUOX1 | DUOXA1 |
| EDN1 | ADRB2 |
| EDN1 | MMP9 |
| EDN1 | NMU |
| EDN1 | NMUR1 |
| EDN1 | CHRM1 |
| EDN1 | GNG11 |
| ENTPD8 | TK1 |
| F11 | F12 |

| | |
|---|---|
| FABP5 | RETN |
| FABP5 | FABP4 |
| FIGF | VWF |
| FIGF | COL1A1 |
| GNG11 | RAMP3 |
| GOLM1 | PCSK9 |
| GOLM1 | CHRDL1 |
| GOLM1 | VGF |
| GPC3 | GOLM1 |
| GPC3 | DCN |
| GPC3 | IHH |
| GPC3 | CHRDL1 |
| GPC3 | PCSK9 |
| GPC3 | WNT3A |
| GPC3 | VGF |
| GPER1 | NMU |
| GPER1 | SSTR1 |
| GPER1 | CXCL13 |
| GPER1 | NMUR1 |
| GPER1 | CX3CR1 |
| GPER1 | GNG11 |
| GPIHBP1 | LPL |
| GPIHBP1 | LY6D |
| GPIHBP1 | NTNG1 |
| GPIHBP1 | LYPD1 |
| GPIHBP1 | RECK |
| GPIHBP1 | CD52 |
| GPIHBP1 | CEACAM5 |
| GPX3 | ALOX5 |
| GRK5 | ADRB2 |
| GRK5 | GNG11 |
| GRK5 | EDN1 |
| GRK5 | NMUR1 |
| GRK5 | CHRM1 |
| GRK5 | WNT3A |
| GRK5 | NMU |
| HBB | HBA2 |
| HBB | CAMP |
| HBD | HBA2 |
| HBD | HBB |
| HHIP | IHH |
| HIST2H3A | HIST1H2AI |

| | |
|---|---|
| HJURP | UBE2C |
| HJURP | CENPF |
| HJURP | KIF2C |
| HJURP | CCNB2 |
| HJURP | TROAP |
| HJURP | BIRC5 |
| HJURP | ASPM |
| HS6ST2 | GPC3 |
| hsa-miR-126-3p | CYBRD1 |
| hsa-miR-126-3p | ITLN2 |
| hsa-miR-126-3p | SELENBP1 |
| hsa-miR-126-3p | DLC1 |
| hsa-miR-126-3p | SGPP2 |
| hsa-miR-126-3p | KCNN4 |
| hsa-miR-126-3p | VWF |
| hsa-miR-126-3p | ABCA12 |
| hsa-miR-126-3p | MS4A15 |
| hsa-miR-126-3p | TMPRSS4 |
| hsa-miR-126-3p | SPOCK2 |
| hsa-miR-126-3p | NLRC4 |
| hsa-miR-126-3p | MEX3A |
| hsa-miR-126-3p | GALNT14 |
| hsa-miR-126-3p | ITLN1 |
| hsa-miR-126-5p | NPNT |
| hsa-miR-126-5p | RNF182 |
| hsa-miR-126-5p | C1orf21 |
| hsa-miR-126-5p | ABCA12 |
| hsa-miR-126-5p | SLIT3 |
| hsa-miR-126-5p | TTN |
| hsa-miR-126-5p | TMEM100 |
| hsa-miR-126-5p | MME |
| hsa-miR-130b-5p | PDK4 |
| hsa-miR-130b-5p | DUOX1 |
| hsa-miR-130b-5p | CACNA2D2 |
| hsa-miR-130b-5p | GPA33 |
| hsa-miR-130b-5p | NCKAP5 |
| hsa-miR-130b-5p | C1orf116 |
| hsa-miR-130b-5p | CYP2F1 |
| hsa-miR-130b-5p | CDHR4 |
| hsa-miR-130b-5p | AOC3 |
| hsa-miR-130b-5p | CACNG6 |
| hsa-miR-130b-5p | RSPH1 |

| | |
|---|---|
| hsa-miR-130b-5p | LRRN3 |
| hsa-miR-130b-5p | TCF21 |
| hsa-miR-130b-5p | CDH13 |
| hsa-miR-130b-5p | CLIC5 |
| hsa-miR-130b-5p | CLIC3 |
| hsa-miR-130b-5p | ARHGEF26 |
| hsa-miR-130b-5p | NTNG1 |
| hsa-miR-130b-5p | CAV1 |
| hsa-miR-130b-5p | GIMAP6 |
| hsa-miR-130b-5p | GIMAP8 |
| hsa-miR-130b-5p | GPRIN2 |
| hsa-miR-130b-5p | ADRB1 |
| hsa-miR-130b-5p | HLF |
| hsa-miR-130b-5p | CES1 |
| hsa-miR-130b-5p | TROAP |
| hsa-miR-130b-5p | PDZD2 |
| hsa-miR-130b-5p | UHRF1 |
| hsa-miR-130b-5p | STRA6 |
| hsa-miR-130b-5p | ITGA8 |
| hsa-miR-130b-5p | NMUR1 |
| hsa-miR-130b-5p | CENPF |
| hsa-miR-130b-5p | DUOXA1 |
| hsa-miR-130b-5p | TUBB3 |
| hsa-miR-130b-5p | MGAT3 |
| hsa-miR-130b-5p | SVEP1 |
| hsa-miR-130b-5p | FAM189A2 |
| hsa-miR-130b-5p | FAM183A |
| hsa-miR-130b-5p | SPOCK2 |
| hsa-miR-130b-5p | ANGPT1 |
| hsa-miR-130b-5p | SPAG5 |
| hsa-miR-130b-5p | LYPD1 |
| hsa-miR-130b-5p | TNS4 |
| hsa-miR-130b-5p | TNS1 |
| hsa-miR-130b-5p | SFTPA1 |
| hsa-miR-130b-5p | SFTPA2 |
| hsa-miR-130b-5p | CAV3 |
| hsa-miR-130b-5p | MAMDC2 |
| hsa-miR-130b-5p | FRAS1 |
| hsa-miR-130b-5p | ABCA12 |
| hsa-miR-130b-5p | C9orf24 |
| hsa-miR-130b-5p | PTPN21 |
| hsa-miR-130b-5p | SLIT2 |

| | |
|---|---|
| hsa-miR-130b-5p | SLIT3 |
| hsa-miR-130b-5p | PCSK9 |
| hsa-miR-130b-5p | C1orf21 |
| hsa-miR-130b-5p | TTN |
| hsa-miR-130b-5p | TMEM63C |
| hsa-miR-130b-5p | ACADL |
| hsa-miR-130b-5p | TMPRSS4 |
| hsa-miR-130b-5p | AQP4 |
| hsa-miR-130b-5p | F11 |
| hsa-miR-130b-5p | VGF |
| hsa-miR-130b-5p | ROBO4 |
| hsa-miR-130b-5p | ERBB4 |
| hsa-miR-130b-5p | HMGB3 |
| hsa-miR-130b-5p | RTKN2 |
| hsa-miR-130b-5p | BTNL9 |
| hsa-miR-130b-5p | GPIHBP1 |
| hsa-miR-130b-5p | CDO1 |
| hsa-miR-130b-5p | A2M |
| hsa-miR-130b-5p | POU2AF1 |
| hsa-miR-130b-5p | LGSN |
| hsa-miR-130b-5p | ITLN1 |
| hsa-miR-130b-5p | HEG1 |
| hsa-miR-130b-5p | LRRC36 |
| hsa-miR-130b-5p | EPHA10 |
| hsa-miR-130b-5p | RSPO4 |
| hsa-miR-130b-5p | KIAA0408 |
| hsa-miR-130b-5p | ZBED2 |
| hsa-miR-130b-5p | SYT7 |
| hsa-miR-130b-5p | NPNT |
| hsa-miR-130b-5p | CGNL1 |
| hsa-miR-130b-5p | FAM83A |
| hsa-miR-130b-5p | MME |
| hsa-miR-130b-5p | SEMA5A |
| hsa-miR-130b-5p | DCDC2B |
| hsa-miR-130b-5p | CYBRD1 |
| hsa-miR-130b-5p | CEACAM5 |
| hsa-miR-130b-5p | DSP |
| hsa-miR-130b-5p | CR2 |
| hsa-miR-130b-5p | PI16 |
| hsa-miR-130b-5p | MUC4 |
| hsa-miR-130b-5p | SFTPC |
| hsa-miR-130b-5p | NDRG4 |

| | |
|---|---|
| hsa-miR-130b-5p | RHPN1 |
| hsa-miR-130b-5p | CYP4B1 |
| hsa-miR-130b-5p | CLDN18 |
| hsa-miR-130b-5p | EMCN |
| hsa-miR-130b-5p | MSR1 |
| hsa-miR-130b-5p | CABYR |
| hsa-miR-130b-5p | C10orf67 |
| hsa-miR-135b-5p | SEMA5A |
| hsa-miR-135b-5p | SPAG5 |
| hsa-miR-135b-5p | FAM189A2 |
| hsa-miR-135b-5p | ANGPT1 |
| hsa-miR-135b-5p | ACOXL |
| hsa-miR-135b-5p | CES1 |
| hsa-miR-135b-5p | GINS1 |
| hsa-miR-135b-5p | PTPN21 |
| hsa-miR-135b-5p | SLC1A1 |
| hsa-miR-135b-5p | ABCA8 |
| hsa-miR-135b-5p | MME |
| hsa-miR-135b-5p | ABI3BP |
| hsa-miR-135b-5p | FAM107A |
| hsa-miR-135b-5p | VWF |
| hsa-miR-135b-5p | GSTM5 |
| hsa-miR-135b-5p | TIMP3 |
| hsa-miR-135b-5p | KIAA0408 |
| hsa-miR-135b-5p | SLIT2 |
| hsa-miR-135b-5p | PID1 |
| hsa-miR-135b-5p | PTPRB |
| hsa-miR-135b-5p | A2M |
| hsa-miR-135b-5p | TROAP |
| hsa-miR-135b-5p | F11 |
| hsa-miR-135b-5p | PCSK9 |
| hsa-miR-135b-5p | DSP |
| hsa-miR-135b-5p | EPN3 |
| hsa-miR-135b-5p | SLC39A8 |
| hsa-miR-135b-5p | CR2 |
| hsa-miR-135b-5p | SLC19A3 |
| hsa-miR-135b-5p | DES |
| hsa-miR-135b-5p | HEG1 |
| hsa-miR-135b-5p | CARD14 |
| hsa-miR-135b-5p | PXMP4 |
| hsa-miR-135b-5p | VEPH1 |
| hsa-miR-135b-5p | DLC1 |

| | |
|---|---|
| hsa-miR-135b-5p | MUC4 |
| hsa-miR-135b-5p | WISP2 |
| hsa-miR-135b-5p | AGER |
| hsa-miR-135b-5p | SYT7 |
| hsa-miR-135b-5p | TTN |
| hsa-miR-135b-5p | NTNG1 |
| hsa-miR-135b-5p | ASPM |
| hsa-miR-135b-5p | C1orf21 |
| hsa-miR-135b-5p | VIPR1 |
| hsa-miR-135b-5p | RNF182 |
| hsa-miR-135b-5p | ADAMTS8 |
| hsa-miR-135b-5p | PSAT1 |
| hsa-miR-135b-5p | CABYR |
| hsa-miR-135b-5p | CHRM1 |
| hsa-miR-135b-5p | EPHA10 |
| hsa-miR-135b-5p | DCN |
| hsa-miR-135b-5p | PI16 |
| hsa-miR-135b-5p | DNAI2 |
| hsa-miR-139-5p | CDCA7 |
| hsa-miR-139-5p | RBP4 |
| hsa-miR-139-5p | TTN |
| hsa-miR-139-5p | GRK5 |
| hsa-miR-139-5p | AFF3 |
| hsa-miR-139-5p | FAM107A |
| hsa-miR-139-5p | ARHGEF26 |
| hsa-miR-139-5p | CFD |
| hsa-miR-139-5p | GINS1 |
| hsa-miR-139-5p | AQP1 |
| hsa-miR-139-5p | CACNA2D2 |
| hsa-miR-139-5p | HBD |
| hsa-miR-139-5p | PYCR1 |
| hsa-miR-139-5p | PSAT1 |
| hsa-miR-139-5p | SLC1A1 |
| hsa-miR-139-5p | JAM2 |
| hsa-miR-139-5p | TPPP3 |
| hsa-miR-139-5p | ROBO4 |
| hsa-miR-139-5p | SYT12 |
| hsa-miR-139-5p | GPM6A |
| hsa-miR-139-5p | GPM6B |
| hsa-miR-139-5p | PLAC8 |
| hsa-miR-139-5p | HHIP |
| hsa-miR-139-5p | LIMCH1 |

| | |
|---|---|
| hsa-miR-139-5p | CX3CR1 |
| hsa-miR-139-5p | LRRC36 |
| hsa-miR-139-5p | TUBB3 |
| hsa-miR-139-5p | CHIA |
| hsa-miR-139-5p | AGER |
| hsa-miR-139-5p | ABCA8 |
| hsa-miR-139-5p | EDN1 |
| hsa-miR-139-5p | STXBP6 |
| hsa-miR-139-5p | CLEC14A |
| hsa-miR-139-5p | FRAS1 |
| hsa-miR-139-5p | METTL7A |
| hsa-miR-139-5p | TMPRSS4 |
| hsa-miR-139-5p | SEMA5A |
| hsa-miR-139-5p | LGALSL |
| hsa-miR-139-5p | PTPRB |
| hsa-miR-139-5p | FABP4 |
| hsa-miR-139-5p | POU2AF1 |
| hsa-miR-139-5p | PXMP4 |
| hsa-miR-139-5p | FAM83A |
| hsa-miR-139-5p | CPAMD8 |
| hsa-miR-139-5p | TNS1 |
| hsa-miR-139-5p | HMGB3 |
| hsa-miR-139-5p | CRYAB |
| hsa-miR-139-5p | NLRC4 |
| hsa-miR-139-5p | XDH |
| hsa-miR-139-5p | GKN2 |
| hsa-miR-139-5p | BTNL9 |
| hsa-miR-139-5p | TMEM63C |
| hsa-miR-139-5p | SPOCK2 |
| hsa-miR-139-5p | DUOX1 |
| hsa-miR-139-5p | SVEP1 |
| hsa-miR-139-5p | VIPR1 |
| hsa-miR-139-5p | BIRC5 |
| hsa-miR-139-5p | SLC6A4 |
| hsa-miR-139-5p | SRPX |
| hsa-miR-139-5p | FRMD3 |
| hsa-miR-139-5p | FGL1 |
| hsa-miR-139-5p | CAV2 |
| hsa-miR-139-5p | LYVE1 |
| hsa-miR-139-5p | NTNG1 |
| hsa-miR-139-5p | PEBP4 |
| hsa-miR-139-5p | MUC4 |

| | |
|---|---|
| hsa-miR-139-5p | C14orf132 |
| hsa-miR-139-5p | PKMYT1 |
| hsa-miR-139-5p | MRC1 |
| hsa-miR-139-5p | RSPH1 |
| hsa-miR-139-5p | C8B |
| hsa-miR-139-5p | C2orf40 |
| hsa-miR-139-5p | ANKRD29 |
| hsa-miR-139-5p | ANGPT1 |
| hsa-miR-139-5p | ZBED2 |
| hsa-miR-139-5p | DNAI2 |
| hsa-miR-139-5p | SUSD2 |
| hsa-miR-139-5p | HEG1 |
| hsa-miR-139-5p | CGNL1 |
| hsa-miR-139-5p | LIMS2 |
| hsa-miR-139-5p | HSD17B6 |
| hsa-miR-139-5p | F12 |
| hsa-miR-139-5p | TIMP3 |
| hsa-miR-139-5p | CHRDL1 |
| hsa-miR-139-5p | LRRK2 |
| hsa-miR-144-3p | LMO2 |
| hsa-miR-144-3p | TMPRSS4 |
| hsa-miR-144-3p | TTN |
| hsa-miR-144-3p | SCN7A |
| hsa-miR-144-3p | ASPM |
| hsa-miR-144-3p | PDZD2 |
| hsa-miR-144-3p | CHRDL1 |
| hsa-miR-144-3p | LGI3 |
| hsa-miR-144-3p | STX1A |
| hsa-miR-144-5p | CACNA2D2 |
| hsa-miR-144-5p | SPOCK2 |
| hsa-miR-144-5p | TTN |
| hsa-miR-144-5p | MFAP4 |
| hsa-miR-144-5p | C14orf132 |
| hsa-miR-144-5p | SFTPA1 |
| hsa-miR-144-5p | PTPRB |
| hsa-miR-183-5p | LYPD1 |
| hsa-miR-183-5p | CCL14 |
| hsa-miR-183-5p | FERMT1 |
| hsa-miR-183-5p | SH3GL3 |
| hsa-miR-183-5p | FAM189A2 |
| hsa-miR-183-5p | LRRK2 |
| hsa-miR-183-5p | DUOXA1 |

| | |
|---|---|
| hsa-miR-183-5p | GLDN |
| hsa-miR-183-5p | PTPRB |
| hsa-miR-183-5p | MUC4 |
| hsa-miR-183-5p | THBD |
| hsa-miR-183-5p | IQGAP3 |
| hsa-miR-183-5p | SCARA5 |
| hsa-miR-183-5p | RECQL4 |
| hsa-miR-183-5p | C11orf88 |
| hsa-miR-183-5p | LGSN |
| hsa-miR-183-5p | PITX1 |
| hsa-miR-183-5p | DENND3 |
| hsa-miR-183-5p | MRC1 |
| hsa-miR-183-5p | LDLRAD1 |
| hsa-miR-183-5p | CX3CR1 |
| hsa-miR-183-5p | ITGA8 |
| hsa-miR-183-5p | AQP1 |
| hsa-miR-183-5p | GJB2 |
| hsa-miR-183-5p | TTN |
| hsa-miR-183-5p | CALCRL |
| hsa-miR-183-5p | MYZAP |
| hsa-miR-183-5p | SLPI |
| hsa-miR-183-5p | ABCA12 |
| hsa-miR-183-5p | MS4A7 |
| hsa-miR-183-5p | C14orf132 |
| hsa-miR-183-5p | EPAS1 |
| hsa-miR-183-5p | SYT7 |
| hsa-miR-183-5p | DES |
| hsa-miR-183-5p | GPM6B |
| hsa-miR-183-5p | MME |
| hsa-miR-183-5p | ABCA3 |
| hsa-miR-183-5p | FAM183A |
| hsa-miR-183-5p | PGC |
| hsa-miR-183-5p | ENTPD8 |
| hsa-miR-183-5p | KCNN4 |
| hsa-miR-183-5p | TNS1 |
| hsa-miR-183-5p | CABYR |
| hsa-miR-183-5p | F11 |
| hsa-miR-183-5p | ERBB4 |
| hsa-miR-183-5p | EPN3 |
| hsa-miR-183-5p | CCNB1 |
| hsa-miR-183-5p | CHRM1 |
| hsa-miR-183-5p | SLIT2 |

| | |
|---|---|
| hsa-miR-183-5p | CDH13 |
| hsa-miR-183-5p | ADH1C |
| hsa-miR-183-5p | NCKAP5 |
| hsa-miR-183-5p | ACOXL |
| hsa-miR-183-5p | AOC3 |
| hsa-miR-183-5p | AGER |
| hsa-miR-183-5p | CRTAC1 |
| hsa-miR-183-5p | DUOX1 |
| hsa-miR-183-5p | RNF182 |
| hsa-miR-183-5p | GPR152 |
| hsa-miR-183-5p | HIGD1B |
| hsa-miR-183-5p | SCN7A |
| hsa-miR-183-5p | ANGPT1 |
| hsa-miR-183-5p | XDH |
| hsa-miR-183-5p | PSAT1 |
| hsa-miR-183-5p | CDHR4 |
| hsa-miR-183-5p | S100P |
| hsa-miR-183-5p | CD36 |
| hsa-miR-183-5p | RTKN2 |
| hsa-miR-183-5p | FRMD3 |
| hsa-miR-183-5p | PDZD2 |
| hsa-miR-183-5p | PDE5A |
| hsa-miR-183-5p | VGLL3 |
| hsa-miR-183-5p | PXMP4 |
| hsa-miR-183-5p | PYCR1 |
| hsa-miR-183-5p | CR2 |
| hsa-miR-183-5p | SLC7A11 |
| hsa-miR-183-5p | MMP9 |
| hsa-miR-183-5p | C1orf116 |
| hsa-miR-183-5p | RAB26 |
| hsa-miR-183-5p | BTNL9 |
| hsa-miR-183-5p | SPP1 |
| hsa-miR-183-5p | DCN |
| hsa-miR-183-5p | LGR4 |
| hsa-miR-183-5p | TNXB |
| hsa-miR-183-5p | C2orf40 |
| hsa-miR-183-5p | POU2AF1 |
| hsa-miR-183-5p | OGN |
| hsa-miR-183-5p | SFTPA2 |
| hsa-miR-183-5p | CAV1 |
| hsa-miR-183-5p | EMCN |
| hsa-miR-183-5p | CPNE7 |

| | |
|---|---|
| hsa-miR-183-5p | VSIG4 |
| hsa-miR-183-5p | HLF |
| hsa-miR-183-5p | MYOC |
| hsa-miR-183-5p | COL6A6 |
| hsa-miR-183-5p | CACNA2D2 |
| hsa-miR-183-5p | LIMS2 |
| hsa-miR-200b-5p | ACADL |
| hsa-miR-200b-5p | SFTPD |
| hsa-miR-200b-5p | FCRL5 |
| hsa-miR-200b-5p | XDH |
| hsa-miR-200b-5p | CABYR |
| hsa-miR-200b-5p | SVEP1 |
| hsa-miR-200b-5p | STXBP6 |
| hsa-miR-200b-5p | CCDC85A |
| hsa-miR-200b-5p | CES1 |
| hsa-miR-200b-5p | FXYD1 |
| hsa-miR-200b-5p | LYPD1 |
| hsa-miR-200b-5p | CARD14 |
| hsa-miR-200b-5p | KIF2C |
| hsa-miR-200b-5p | VWF |
| hsa-miR-200b-5p | GOLM1 |
| hsa-miR-200b-5p | PTPN21 |
| hsa-miR-200b-5p | DLC1 |
| hsa-miR-200b-5p | CDCA5 |
| hsa-miR-200b-5p | LAMP3 |
| hsa-miR-200b-5p | CX3CR1 |
| hsa-miR-200b-5p | MME |
| hsa-miR-200b-5p | EPAS1 |
| hsa-miR-200b-5p | CDH3 |
| hsa-miR-200b-5p | HLF |
| hsa-miR-200b-5p | CACNG6 |
| hsa-miR-200b-5p | NPNT |
| hsa-miR-200b-5p | CD36 |
| hsa-miR-200b-5p | GSTM5 |
| hsa-miR-200b-5p | ANKRD29 |
| hsa-miR-200b-5p | SPARCL1 |
| hsa-miR-200b-5p | ACVRL1 |
| hsa-miR-200b-5p | LIMCH1 |
| hsa-miR-200b-5p | S100P |
| hsa-miR-200b-5p | CR2 |
| hsa-miR-200b-5p | CYP4B1 |
| hsa-miR-200b-5p | CHRDL1 |

| | |
|---|---|
| hsa-miR-200b-5p | GPX3 |
| hsa-miR-200b-5p | MAMDC2 |
| hsa-miR-200b-5p | SPOCK2 |
| hsa-miR-200b-5p | FHL2 |
| hsa-miR-200b-5p | TROAP |
| hsa-miR-200b-5p | ANGPT1 |
| hsa-miR-200b-5p | ABCA3 |
| hsa-miR-200b-5p | CAV2 |
| hsa-miR-200b-5p | SLIT2 |
| hsa-miR-200b-5p | SLIT3 |
| hsa-miR-200b-5p | ITLN1 |
| hsa-miR-200b-5p | GPC3 |
| hsa-miR-200b-5p | MSR1 |
| hsa-miR-200b-5p | AOC3 |
| hsa-miR-200b-5p | BIRC5 |
| hsa-miR-200b-5p | PHLDA2 |
| hsa-miR-200b-5p | KRT4 |
| hsa-miR-200b-5p | TMPRSS4 |
| hsa-miR-200b-5p | SFTPA2 |
| hsa-miR-200b-5p | HIGD1B |
| hsa-miR-200b-5p | ABCA12 |
| hsa-miR-200b-5p | AQP4 |
| hsa-miR-200b-5p | HMGB3 |
| hsa-miR-200b-5p | MMP1 |
| hsa-miR-200b-5p | WNT3A |
| hsa-miR-200b-5p | TPX2 |
| hsa-miR-200b-5p | GPM6B |
| hsa-miR-200b-5p | A2M |
| hsa-miR-200b-5p | LMO2 |
| hsa-miR-200b-5p | DENND3 |
| hsa-miR-200b-5p | TMEM100 |
| hsa-miR-200b-5p | C10orf67 |
| hsa-miR-200b-5p | CLEC3B |
| hsa-miR-200b-5p | NLRC4 |
| hsa-miR-200b-5p | CALCRL |
| hsa-miR-200b-5p | AGER |
| hsa-miR-200b-5p | FUT3 |
| hsa-miR-200b-5p | INMT |
| hsa-miR-200b-5p | PID1 |
| hsa-miR-200b-5p | CCNB1 |
| hsa-miR-200b-5p | UBE2T |
| hsa-miR-200b-5p | EMCN |

| | |
|---|---|
| hsa-miR-200b-5p | ABCA8 |
| hsa-miR-200b-5p | MGAT3 |
| hsa-miR-200b-5p | CD19 |
| hsa-miR-200b-5p | NTNG1 |
| hsa-miR-200b-5p | RTKN2 |
| hsa-miR-200b-5p | RASAL1 |
| hsa-miR-200b-5p | KIAA0408 |
| hsa-miR-200b-5p | RNF182 |
| hsa-miR-200b-5p | CAV1 |
| hsa-miR-200b-5p | PTPRB |
| hsa-miR-200b-5p | PRAM1 |
| hsa-miR-200b-5p | ADH1A |
| hsa-miR-200b-5p | PROM2 |
| hsa-miR-200b-5p | DUOXA1 |
| hsa-miR-200b-5p | TTN |
| hsa-miR-200b-5p | FRAS1 |
| hsa-miR-200b-5p | HEG1 |
| hsa-miR-200b-5p | CLIC5 |
| hsa-miR-200b-5p | DSP |
| hsa-miR-200b-5p | FRMD3 |
| hsa-miR-200b-5p | SLC7A11 |
| hsa-miR-200b-5p | F12 |
| hsa-miR-200b-5p | CDHR4 |
| hsa-miR-200b-5p | CDC45 |
| hsa-miR-200b-5p | CRYAB |
| hsa-miR-200b-5p | C8B |
| hsa-miR-200b-5p | CPNE7 |
| hsa-miR-200b-5p | FAM107A |
| hsa-miR-200b-5p | ROBO4 |
| hsa-miR-200b-5p | AFF3 |
| hsa-miR-200b-5p | PCSK9 |
| hsa-miR-200b-5p | COL6A6 |
| hsa-miR-21-5p | SPP1 |
| hsa-miR-21-5p | CEP55 |
| hsa-miR-21-5p | CYBRD1 |
| hsa-miR-21-5p | AQP1 |
| hsa-miR-21-5p | GPM6A |
| hsa-miR-21-5p | C8B |
| hsa-miR-21-5p | TTN |
| hsa-miR-21-5p | SCN4B |
| hsa-miR-21-5p | SPOCK2 |
| hsa-miR-21-5p | FRAS1 |

| | |
|---|---|
| hsa-miR-21-5p | CHIA |
| hsa-miR-21-5p | CAMP |
| hsa-miR-21-5p | RECK |
| hsa-miR-21-5p | ASPM |
| hsa-miR-21-5p | PTPRB |
| hsa-miR-21-5p | DSP |
| hsa-miR-21-5p | SPAG5 |
| hsa-miR-21-5p | ADAMTSL3 |
| hsa-miR-21-5p | NLRC4 |
| hsa-miR-21-5p | GALNT14 |
| hsa-miR-21-5p | NCKAP5 |
| hsa-miR-21-5p | ITGA8 |
| hsa-miR-21-5p | SLC39A8 |
| hsa-miR-218-1-3p | C1orf116 |
| hsa-miR-218-1-3p | FCRL5 |
| hsa-miR-218-1-3p | SSTR1 |
| hsa-miR-218-1-3p | METTL7A |
| hsa-miR-218-1-3p | CAMP |
| hsa-miR-218-1-3p | SLIT2 |
| hsa-miR-218-1-3p | AQP1 |
| hsa-miR-218-1-3p | VEPH1 |
| hsa-miR-218-1-3p | GIMAP8 |
| hsa-miR-218-1-3p | CPA3 |
| hsa-miR-218-1-3p | GPRIN2 |
| hsa-miR-218-1-3p | CAV1 |
| hsa-miR-218-1-3p | MME |
| hsa-miR-218-1-3p | ACOXL |
| hsa-miR-218-1-3p | STRA6 |
| hsa-miR-218-1-3p | SYT12 |
| hsa-miR-218-1-3p | EPN3 |
| hsa-miR-218-1-3p | EMCN |
| hsa-miR-218-1-3p | NCKAP5 |
| hsa-miR-218-1-3p | SGPP2 |
| hsa-miR-218-1-3p | DSP |
| hsa-miR-218-1-3p | DENND3 |
| hsa-miR-218-1-3p | FRAS1 |
| hsa-miR-218-1-3p | ENTPD8 |
| hsa-miR-218-1-3p | GPM6A |
| hsa-miR-218-1-3p | ANXA3 |
| hsa-miR-218-1-3p | TCF21 |
| hsa-miR-218-1-3p | F11 |
| hsa-miR-218-1-3p | F12 |

| | |
|---|---|
| hsa-miR-218-1-3p | ABCA12 |
| hsa-miR-218-1-3p | UHRF1 |
| hsa-miR-218-1-3p | DUOX1 |
| hsa-miR-218-1-3p | CDHR4 |
| hsa-miR-218-1-3p | IQGAP3 |
| hsa-miR-218-1-3p | PDE5A |
| hsa-miR-218-1-3p | CD19 |
| hsa-miR-218-1-3p | EFCAB1 |
| hsa-miR-218-1-3p | SCARA5 |
| hsa-miR-218-1-3p | ARHGEF26 |
| hsa-miR-218-1-3p | CDH3 |
| hsa-miR-218-1-3p | CYP2F1 |
| hsa-miR-218-1-3p | HEG1 |
| hsa-miR-218-1-3p | GALNT14 |
| hsa-miR-218-1-3p | PTPRB |
| hsa-miR-218-1-3p | NLRC4 |
| hsa-miR-218-1-3p | ASPM |
| hsa-miR-218-1-3p | ANKRD29 |
| hsa-miR-218-1-3p | ADAMTSL3 |
| hsa-miR-218-1-3p | FXYD1 |
| hsa-miR-218-1-3p | DLC1 |
| hsa-miR-218-1-3p | DNAH12 |
| hsa-miR-218-1-3p | CEACAM5 |
| hsa-miR-218-1-3p | CENPF |
| hsa-miR-218-1-3p | ADAMTS8 |
| hsa-miR-218-1-3p | FAM107A |
| hsa-miR-218-1-3p | ERBB4 |
| hsa-miR-218-1-3p | PTPN21 |
| hsa-miR-218-1-3p | COL1A1 |
| hsa-miR-218-1-3p | PDZD2 |
| hsa-miR-218-1-3p | FBLN5 |
| hsa-miR-218-1-3p | VSIG2 |
| hsa-miR-218-1-3p | COL6A6 |
| hsa-miR-218-1-3p | C2orf40 |
| hsa-miR-218-1-3p | C9orf24 |
| hsa-miR-218-1-3p | TNS1 |
| hsa-miR-218-1-3p | LIMCH1 |
| hsa-miR-218-1-3p | SVEP1 |
| hsa-miR-218-1-3p | PI16 |
| hsa-miR-218-1-3p | SCGB1A1 |
| hsa-miR-218-1-3p | SH3GL3 |
| hsa-miR-218-1-3p | SCN4B |

| | |
|---|---|
| hsa-miR-218-1-3p | ITGA8 |
| hsa-miR-218-1-3p | EMP2 |
| hsa-miR-218-1-3p | C8B |
| hsa-miR-218-1-3p | TTN |
| hsa-miR-218-1-3p | DNAI2 |
| hsa-miR-218-1-3p | CD36 |
| hsa-miR-218-2-3p | HSD17B6 |
| hsa-miR-218-2-3p | C1orf21 |
| hsa-miR-218-2-3p | GPIHBP1 |
| hsa-miR-218-2-3p | GDF10 |
| hsa-miR-218-2-3p | AFF3 |
| hsa-miR-218-2-3p | LYVE1 |
| hsa-miR-218-2-3p | DUOXA1 |
| hsa-miR-218-2-3p | CCNB1 |
| hsa-miR-218-2-3p | ADH1C |
| hsa-miR-218-2-3p | CYBRD1 |
| hsa-miR-218-2-3p | LGSN |
| hsa-miR-218-2-3p | ABI3BP |
| hsa-miR-218-2-3p | PTPN21 |
| hsa-miR-218-2-3p | DENND3 |
| hsa-miR-218-2-3p | MMP12 |
| hsa-miR-218-2-3p | COL6A6 |
| hsa-miR-218-2-3p | CALCRL |
| hsa-miR-218-2-3p | CX3CR1 |
| hsa-miR-218-2-3p | CENPF |
| hsa-miR-218-2-3p | SEMA5A |
| hsa-miR-218-2-3p | FCRL5 |
| hsa-miR-218-2-3p | TNS4 |
| hsa-miR-218-2-3p | C8B |
| hsa-miR-218-2-3p | STX1A |
| hsa-miR-218-2-3p | ITGA8 |
| hsa-miR-218-2-3p | GALNT14 |
| hsa-miR-218-2-3p | TMEM63C |
| hsa-miR-218-2-3p | RECQL4 |
| hsa-miR-218-2-3p | SSTR1 |
| hsa-miR-218-2-3p | SH3GL3 |
| hsa-miR-218-2-3p | SFTPC |
| hsa-miR-218-2-3p | SGPP2 |
| hsa-miR-218-2-3p | CHIA |
| hsa-miR-218-2-3p | CAV1 |
| hsa-miR-218-2-3p | CTHRC1 |
| hsa-miR-218-2-3p | DNASE1L3 |

| | |
|---|---|
| hsa-miR-218-2-3p | DLC1 |
| hsa-miR-218-2-3p | AOC3 |
| hsa-miR-218-2-3p | TTN |
| hsa-miR-218-2-3p | EPHA10 |
| hsa-miR-218-2-3p | C9orf24 |
| hsa-miR-218-2-3p | GPRIN2 |
| hsa-miR-218-2-3p | TMPRSS4 |
| hsa-miR-218-2-3p | NCKAP5 |
| hsa-miR-218-2-3p | CRABP2 |
| hsa-miR-218-2-3p | TPX2 |
| hsa-miR-218-2-3p | FRMD3 |
| hsa-miR-218-2-3p | ANGPT1 |
| hsa-miR-218-2-3p | DCN |
| hsa-miR-218-2-3p | TUBB3 |
| hsa-miR-218-2-3p | CEP55 |
| hsa-miR-218-2-3p | CARD14 |
| hsa-miR-218-2-3p | CGNL1 |
| hsa-miR-218-2-3p | MFAP4 |
| hsa-miR-218-2-3p | PLAC9 |
| hsa-miR-218-2-3p | ANKRD29 |
| hsa-miR-218-2-3p | AQP1 |
| hsa-miR-218-2-3p | MMP9 |
| hsa-miR-218-2-3p | INMT |
| hsa-miR-218-2-3p | VSIG2 |
| hsa-miR-218-2-3p | ACADL |
| hsa-miR-218-2-3p | CES1 |
| hsa-miR-218-2-3p | SFTPD |
| hsa-miR-218-2-3p | MEX3A |
| hsa-miR-218-2-3p | SLIT2 |
| hsa-miR-218-2-3p | GPM6A |
| hsa-miR-218-2-3p | LDLRAD1 |
| hsa-miR-218-2-3p | ENTPD8 |
| hsa-miR-218-2-3p | C1orf116 |
| hsa-miR-218-2-3p | SYT12 |
| hsa-miR-218-2-3p | IRX1 |
| hsa-miR-218-2-3p | SLC19A3 |
| hsa-miR-218-2-3p | TNXB |
| hsa-miR-218-2-3p | CACNA2D2 |
| hsa-miR-218-2-3p | KIF2C |
| hsa-miR-218-2-3p | LRRC36 |
| hsa-miR-218-2-3p | GOLM1 |
| hsa-miR-218-2-3p | MNX1 |

| | |
|---|---|
| hsa-miR-218-2-3p | TIMP3 |
| hsa-miR-218-5p | SPAG5 |
| hsa-miR-218-5p | STX1A |
| hsa-miR-218-5p | CALCRL |
| hsa-miR-218-5p | SLC19A3 |
| hsa-miR-218-5p | FGL1 |
| hsa-miR-218-5p | GKN2 |
| hsa-miR-218-5p | PDZD2 |
| hsa-miR-218-5p | ACADL |
| hsa-miR-218-5p | ASPM |
| hsa-miR-218-5p | COL6A6 |
| hsa-miR-218-5p | ADRB1 |
| hsa-miR-218-5p | GALNT14 |
| hsa-miR-218-5p | GOLM1 |
| hsa-miR-218-5p | FAM83A |
| hsa-miR-218-5p | CSH1 |
| hsa-miR-218-5p | EMCN |
| hsa-miR-218-5p | ABI3BP |
| hsa-miR-218-5p | VEPH1 |
| hsa-miR-218-5p | PXMP4 |
| hsa-miR-30a-3p | TPX2 |
| hsa-miR-30a-3p | LIMS2 |
| hsa-miR-30a-3p | MS4A7 |
| hsa-miR-30a-3p | PROM2 |
| hsa-miR-30a-3p | HS6ST2 |
| hsa-miR-30a-3p | GLDN |
| hsa-miR-30a-3p | MRC1 |
| hsa-miR-30a-3p | RECQL4 |
| hsa-miR-30a-3p | SLC19A3 |
| hsa-miR-30a-3p | SLC1A1 |
| hsa-miR-30a-3p | NDNF |
| hsa-miR-30a-3p | ABCA12 |
| hsa-miR-30a-3p | PTPRB |
| hsa-miR-30a-3p | ADAMTSL3 |
| hsa-miR-30a-3p | LYPD1 |
| hsa-miR-30a-3p | FRAS1 |
| hsa-miR-30a-3p | NDRG4 |
| hsa-miR-30a-3p | CENPF |
| hsa-miR-30a-3p | HJURP |
| hsa-miR-30a-3p | CDH13 |
| hsa-miR-30a-3p | CDH3 |
| hsa-miR-30a-3p | CHRM1 |

| | |
|---|---|
| hsa-miR-30a-3p | ABCA8 |
| hsa-miR-30a-3p | DCN |
| hsa-miR-30a-3p | GPM6B |
| hsa-miR-30a-3p | CABYR |
| hsa-miR-30a-3p | AQP1 |
| hsa-miR-30a-3p | PKMYT1 |
| hsa-miR-30a-3p | NCKAP5 |
| hsa-miR-30a-3p | CTHRC1 |
| hsa-miR-30a-3p | AFF3 |
| hsa-miR-30a-3p | SELENBP1 |
| hsa-miR-30a-3p | ADH1B |
| hsa-miR-30a-3p | SCN7A |
| hsa-miR-30a-3p | MEX3A |
| hsa-miR-30a-3p | MARCO |
| hsa-miR-30a-3p | NLRC4 |
| hsa-miR-30a-3p | COL1A1 |
| hsa-miR-30a-3p | FCRL5 |
| hsa-miR-30a-3p | ZBED2 |
| hsa-miR-30a-3p | DNASE1L3 |
| hsa-miR-30a-3p | PCOLCE2 |
| hsa-miR-30a-3p | LIMCH1 |
| hsa-miR-30a-3p | OLR1 |
| hsa-miR-30a-3p | COL6A6 |
| hsa-miR-30a-3p | PCSK9 |
| hsa-miR-30a-3p | FRMD3 |
| hsa-miR-30a-3p | CHRDL1 |
| hsa-miR-30a-3p | ACVRL1 |
| hsa-miR-30a-3p | TTN |
| hsa-miR-30a-3p | SFTPA2 |
| hsa-miR-30a-3p | SOSTDC1 |
| hsa-miR-30a-3p | ABI3BP |
| hsa-miR-30a-3p | MMP12 |
| hsa-miR-30a-3p | GIMAP6 |
| hsa-miR-30a-3p | MAMDC2 |
| hsa-miR-30a-3p | DLC1 |
| hsa-miR-30a-3p | MS4A2 |
| hsa-miR-30a-3p | FAM189A2 |
| hsa-miR-30a-3p | ANGPT1 |
| hsa-miR-30a-3p | LRRK2 |
| hsa-miR-30a-3p | RSPH1 |
| hsa-miR-30a-3p | TPPP3 |
| hsa-miR-30a-3p | GPM6A |

| | |
|---|---|
| hsa-miR-30a-3p | CEP55 |
| hsa-miR-30a-3p | TMEM100 |
| hsa-miR-30a-3p | ANXA3 |
| hsa-miR-30a-3p | CLIC5 |
| hsa-miR-30a-3p | TEK |
| hsa-miR-30a-3p | C1orf21 |
| hsa-miR-30a-3p | MME |
| hsa-miR-30a-3p | CHIA |
| hsa-miR-30a-3p | CGNL1 |
| hsa-miR-30a-3p | MMRN1 |
| hsa-miR-30a-3p | LPL |
| hsa-miR-30a-3p | ASPM |
| hsa-miR-30a-3p | KRT4 |
| hsa-miR-30a-3p | DENND3 |
| hsa-miR-30a-3p | FHL2 |
| hsa-miR-30a-3p | CLDN18 |
| hsa-miR-30a-3p | HSD17B6 |
| hsa-miR-30a-3p | GDF10 |
| hsa-miR-30a-3p | GPRIN2 |
| hsa-miR-30a-5p | TCF21 |
| hsa-miR-30a-5p | FMO2 |
| hsa-miR-30a-5p | LRRC36 |
| hsa-miR-30a-5p | AOC3 |
| hsa-miR-30a-5p | LAMP3 |
| hsa-miR-30a-5p | FRAS1 |
| hsa-miR-30a-5p | ABI3BP |
| hsa-miR-30a-5p | ROBO4 |
| hsa-miR-30a-5p | GLIPR2 |
| hsa-miR-30a-5p | SPP1 |
| hsa-miR-30a-5p | IHH |
| hsa-miR-30a-5p | EPHA10 |
| hsa-miR-30a-5p | IQGAP3 |
| hsa-miR-30a-5p | ABCA8 |
| hsa-miR-30a-5p | FBLN5 |
| hsa-miR-338-3p | GDF10 |
| hsa-miR-338-3p | GALNT14 |
| hsa-miR-338-3p | TPX2 |
| hsa-miR-338-3p | ASPM |
| hsa-miR-338-3p | LRRC36 |
| hsa-miR-338-3p | MSR1 |
| hsa-miR-338-3p | ERBB4 |
| hsa-miR-338-3p | BTNL9 |

| | |
|---|---|
| hsa-miR-338-3p | ROBO4 |
| hsa-miR-338-3p | AGER |
| hsa-miR-338-3p | CDO1 |
| hsa-miR-338-3p | DYNLRB2 |
| hsa-miR-338-3p | TNS1 |
| hsa-miR-338-3p | GPM6B |
| hsa-miR-338-3p | HSD17B6 |
| hsa-miR-338-3p | STXBP6 |
| hsa-miR-338-3p | SLC1A1 |
| hsa-miR-338-3p | DSP |
| hsa-miR-338-3p | COL1A1 |
| hsa-miR-338-3p | TPPP3 |
| hsa-miR-338-3p | NLRC4 |
| hsa-miR-338-3p | GPRIN2 |
| hsa-miR-338-3p | SELENBP1 |
| hsa-miR-338-3p | NPNT |
| hsa-miR-338-3p | CENPF |
| hsa-miR-338-3p | CTHRC1 |
| hsa-miR-338-3p | XDH |
| hsa-miR-338-3p | ACVRL1 |
| hsa-miR-338-3p | VEPH1 |
| hsa-miR-338-3p | RASAL1 |
| hsa-miR-338-3p | CDH3 |
| hsa-miR-338-3p | CCNB1 |
| hsa-miR-338-3p | LGSN |
| hsa-miR-338-3p | HHIP |
| hsa-miR-338-3p | STRA6 |
| hsa-miR-338-3p | HIGD1B |
| hsa-miR-338-3p | CRTAC1 |
| hsa-miR-338-3p | HBD |
| hsa-miR-338-3p | CD36 |
| hsa-miR-338-3p | PKMYT1 |
| hsa-miR-338-3p | CHIA |
| hsa-miR-338-3p | ADH1A |
| hsa-miR-338-3p | OGN |
| hsa-miR-338-3p | TTN |
| hsa-miR-338-3p | ITGA8 |
| hsa-miR-338-3p | RECK |
| hsa-miR-338-3p | DNAI2 |
| hsa-miR-338-3p | CA4 |
| hsa-miR-338-3p | FAM107A |
| hsa-miR-338-3p | ACADL |

| | |
|---|---|
| hsa-miR-338-3p | MME |
| hsa-miR-338-3p | FRMD3 |
| hsa-miR-338-3p | HBB |
| hsa-miR-338-3p | SLC6A4 |
| hsa-miR-338-3p | MRC1 |
| hsa-miR-338-3p | PLAC8 |
| hsa-miR-338-3p | COL6A6 |
| hsa-miR-338-3p | MMP9 |
| hsa-miR-338-3p | METTL7A |
| hsa-miR-338-3p | NCKAP5 |
| hsa-miR-338-3p | MUC4 |
| hsa-miR-338-3p | CDC45 |
| hsa-miR-338-3p | ANGPT1 |
| hsa-miR-338-3p | SVEP1 |
| hsa-miR-338-3p | GIMAP6 |
| hsa-miR-451a | ASPM |
| hsa-miR-451a | NPNT |
| hsa-miR-451a | MS4A7 |
| hsa-miR-451a | VGLL3 |
| hsa-miR-451a | TK1 |
| hsa-miR-451a | EFCAB1 |
| hsa-miR-451a | EMP2 |
| hsa-miR-451a | PTPRB |
| hsa-miR-451a | KIAA0408 |
| hsa-miR-451a | SSTR1 |
| hsa-miR-451a | SVEP1 |
| hsa-miR-451a | FRMD3 |
| hsa-miR-451a | CCNB1 |
| hsa-miR-451a | CALCRL |
| hsa-miR-451a | FABP5 |
| hsa-miR-451a | ERBB4 |
| hsa-miR-451b | FRAS1 |
| hsa-miR-451b | SVEP1 |
| hsa-miR-451b | TMPRSS4 |
| hsa-miR-451b | FGL1 |
| hsa-miR-451b | CALCRL |
| hsa-miR-451b | CDO1 |
| hsa-miR-451b | BCHE |
| hsa-miR-451b | CGNL1 |
| hsa-miR-451b | NLRC4 |
| hsa-miR-451b | MRC1 |
| hsa-miR-451b | EPHA10 |

| | |
|---|---|
| hsa-miR-451b | LGSN |
| hsa-miR-451b | AOC3 |
| hsa-miR-451b | EPAS1 |
| hsa-miR-451b | CD36 |
| hsa-miR-451b | TTN |
| hsa-miR-486-5p | CCNB1 |
| hsa-miR-486-5p | TNXB |
| hsa-miR-486-5p | HHIP |
| hsa-miR-486-5p | COL10A1 |
| hsa-miR-486-5p | UBE2T |
| hsa-miR-486-5p | CRTAC1 |
| hsa-miR-486-5p | ANXA3 |
| hsa-miR-486-5p | LRRN3 |
| hsa-miR-486-5p | ADH1B |
| hsa-miR-486-5p | XDH |
| hsa-miR-486-5p | SGPP2 |
| hsa-miR-486-5p | FMO2 |
| hsa-miR-486-5p | EMP2 |
| hsa-miR-486-5p | PDE5A |
| hsa-miR-486-5p | GPA33 |
| hsa-miR-486-5p | SPINK1 |
| hsa-miR-486-5p | PCSK9 |
| hsa-miR-486-5p | FAM83A |
| hsa-miR-486-5p | AFF3 |
| hsa-miR-486-5p | MMP12 |
| hsa-miR-486-5p | GSTM5 |
| hsa-miR-486-5p | DENND3 |
| hsa-miR-486-5p | CHRM1 |
| hsa-miR-486-5p | RETN |
| hsa-miR-486-5p | EPHA10 |
| hsa-miR-486-5p | CYBRD1 |
| hsa-miR-486-5p | RAB26 |
| hsa-miR-486-5p | PYCR1 |
| hsa-miR-486-5p | GPM6A |
| hsa-miR-486-5p | GPM6B |
| hsa-miR-486-5p | SFTPA2 |
| hsa-miR-486-5p | SFTPA1 |
| hsa-miR-486-5p | ASPM |
| hsa-miR-486-5p | CDHR4 |
| hsa-miR-486-5p | CDH3 |
| hsa-miR-486-5p | CYP4B1 |
| hsa-miR-486-5p | CD36 |

| | |
|---|---|
| hsa-miR-486-5p | OLR1 |
| hsa-miR-486-5p | B3GNT3 |
| hsa-miR-486-5p | FGL1 |
| hsa-miR-486-5p | SPP1 |
| hsa-miR-486-5p | DUOXA1 |
| hsa-miR-486-5p | CA2 |
| hsa-miR-486-5p | TEK |
| hsa-miR-486-5p | GLIPR2 |
| hsa-miR-486-5p | ADH1A |
| hsa-miR-486-5p | DUOX1 |
| hsa-miR-486-5p | MUC4 |
| hsa-miR-486-5p | MME |
| hsa-miR-486-5p | STRA6 |
| hsa-miR-486-5p | TPX2 |
| hsa-miR-486-5p | CAV3 |
| hsa-miR-486-5p | GINS1 |
| hsa-miR-486-5p | MYZAP |
| hsa-miR-486-5p | MYOC |
| hsa-miR-486-5p | METTL7A |
| hsa-miR-486-5p | FCN3 |
| hsa-miR-486-5p | PITX1 |
| hsa-miR-486-5p | GALNT14 |
| hsa-miR-486-5p | TTN |
| hsa-miR-486-5p | SCARA5 |
| hsa-miR-486-5p | CACNA2D2 |
| hsa-miR-486-5p | DES |
| hsa-miR-486-5p | LDLRAD1 |
| hsa-miR-486-5p | NCKAP5 |
| hsa-miR-486-5p | CABYR |
| hsa-miR-486-5p | RASAL1 |
| hsa-miR-486-5p | HEG1 |
| hsa-miR-486-5p | HSD17B6 |
| hsa-miR-486-5p | FRAS1 |
| hsa-miR-486-5p | LGSN |
| hsa-miR-486-5p | MMP11 |
| hsa-miR-486-5p | TPPP3 |
| hsa-miR-486-5p | DSP |
| hsa-miR-486-5p | LAMP3 |
| hsa-miR-486-5p | ABCA8 |
| hsa-miR-486-5p | ADRB2 |
| hsa-miR-486-5p | CDCA7 |
| hsa-miR-486-5p | CDC45 |

| | |
|---|---|
| hsa-miR-486-5p | PXMP4 |
| hsa-miR-486-5p | FAM107A |
| hsa-miR-486-5p | TMEM100 |
| hsa-miR-486-5p | ALOX5AP |
| hsa-miR-486-5p | TMEM190 |
| hsa-miR-486-5p | GLDN |
| hsa-miR-486-5p | ALOX5 |
| hsa-miR-486-5p | CEACAM5 |
| hsa-miR-486-5p | RTKN2 |
| hsa-miR-486-5p | CLIC5 |
| hsa-miR-486-5p | RECK |
| hsa-miR-486-5p | STXBP6 |
| hsa-miR-486-5p | CARD14 |
| hsa-miR-486-5p | FRMD3 |
| hsa-miR-486-5p | SLIT3 |
| hsa-miR-486-5p | RBP4 |
| hsa-miR-486-5p | CRABP2 |
| hsa-miR-486-5p | VIPR1 |
| hsa-miR-486-5p | NPNT |
| hsa-miR-486-5p | HMGB3 |
| hsa-miR-486-5p | HS6ST2 |
| hsa-miR-486-5p | SLIT2 |
| hsa-miR-486-5p | MAMDC2 |
| hsa-miR-486-5p | C8B |
| hsa-miR-486-5p | ABCA12 |
| hsa-miR-486-5p | CAV1 |
| hsa-miR-486-5p | RNF182 |
| hsa-miR-486-5p | SCN7A |
| hsa-miR-486-5p | KRT4 |
| hsa-miR-486-5p | WIF1 |
| hsa-miR-486-5p | ANGPT1 |
| hsa-miR-486-5p | JAM2 |
| hsa-miR-486-5p | VEPH1 |
| hsa-miR-486-5p | STX1A |
| hsa-miR-486-5p | C1orf21 |
| hsa-miR-486-5p | PDK4 |
| hsa-miR-486-5p | TNS1 |
| hsa-miR-486-5p | PROM2 |
| hsa-miR-486-5p | CR2 |
| hsa-miR-486-5p | SPOCK2 |
| hsa-miR-96-5p | WISP2 |
| hsa-miR-96-5p | ERBB4 |

| | |
|---|---|
| hsa-miR-96-5p | GPRIN2 |
| hsa-miR-96-5p | TMPRSS4 |
| hsa-miR-96-5p | LGSN |
| hsa-miR-96-5p | CRYAB |
| hsa-miR-96-5p | C1orf21 |
| hsa-miR-96-5p | LPL |
| hsa-miR-96-5p | FRAS1 |
| hsa-miR-96-5p | TPX2 |
| hsa-miR-96-5p | SEMA5A |
| hsa-miR-96-5p | PROM2 |
| hsa-miR-96-5p | CACNA2D2 |
| hsa-miR-96-5p | SVEP1 |
| hsa-miR-96-5p | NLRC4 |
| hsa-miR-96-5p | NPNT |
| hsa-miR-96-5p | COL6A6 |
| hsa-miR-96-5p | EMP2 |
| hsa-miR-96-5p | PDK4 |
| hsa-miR-96-5p | JAM2 |
| hsa-miR-96-5p | LGALSL |
| hsa-miR-96-5p | CAV1 |
| hsa-miR-96-5p | EPHA10 |
| hsa-miR-96-5p | RTKN2 |
| hsa-miR-96-5p | PTPRB |
| hsa-miR-96-5p | PTPRH |
| hsa-miR-96-5p | STRA6 |
| hsa-miR-96-5p | ADAMTS8 |
| hsa-miR-96-5p | AOC3 |
| hsa-miR-96-5p | IQGAP3 |
| hsa-miR-96-5p | SFTPA2 |
| hsa-miR-96-5p | CD19 |
| hsa-miR-96-5p | SLC39A8 |
| hsa-miR-96-5p | CSH1 |
| hsa-miR-96-5p | HMGB3 |
| hsa-miR-96-5p | PTPN21 |
| hsa-miR-96-5p | DLC1 |
| hsa-miR-96-5p | CDH13 |
| hsa-miR-96-5p | LIMCH1 |
| hsa-miR-96-5p | GLDN |
| hsa-miR-96-5p | ACADL |
| hsa-miR-96-5p | SLIT2 |
| hsa-miR-96-5p | CD36 |
| hsa-miR-96-5p | SRPX |

| | |
|---|---|
| hsa-miR-96-5p | CHRM1 |
| hsa-miR-96-5p | CPNE7 |
| hsa-miR-96-5p | HS6ST2 |
| hsa-miR-96-5p | SCN7A |
| hsa-miR-96-5p | DUOXA1 |
| hsa-miR-96-5p | SPOCK2 |
| hsa-miR-96-5p | PITX1 |
| hsa-miR-96-5p | RECQL4 |
| hsa-miR-96-5p | GALNT14 |
| hsa-miR-96-5p | CEP55 |
| hsa-miR-96-5p | SELENBP1 |
| hsa-miR-96-5p | VIPR1 |
| hsa-miR-96-5p | FAM189A2 |
| hsa-miR-96-5p | HBD |
| hsa-miR-96-5p | GOLM1 |
| hsa-miR-96-5p | KIAA0408 |
| hsa-miR-96-5p | NTNG1 |
| KIAA0101 | CCNB2 |
| KIAA0101 | CCNB1 |
| KIF2C | CCNB2 |
| KIF2C | CCNB1 |
| KIF2C | CENPF |
| KIF2C | BIRC5 |
| KIF2C | CDCA5 |
| KIF2C | TPX2 |
| KIF2C | UBE2C |
| KIF2C | SPAG5 |
| KIF2C | CEP55 |
| KIF2C | TROAP |
| KIF2C | ASPM |
| LEPREL1 | COL1A1 |
| LGI3 | STX1A |
| LGR4 | RSPO4 |
| LPL | FABP4 |
| LY6D | CEACAM5 |
| LYPD1 | LY6D |
| LYPD1 | NTNG1 |
| LYPD1 | CEACAM5 |
| LYPD1 | CD52 |
| LYPD1 | RECK |
| MARCO | SCGB3A2 |
| MCEMP1 | OLR1 |

| | |
|---|---|
| METTL7A | HBB |
| METTL7A | CAMP |
| METTL7A | MMP9 |
| MFAP4 | FBLN5 |
| MFAP4 | SFTPD |
| MMP1 | MMP11 |
| MMP12 | A2M |
| MMP9 | TIMP3 |
| MMP9 | MMP1 |
| MMP9 | CAMP |
| MMP9 | A2M |
| MMP9 | HBB |
| MMRN1 | VWF |
| MMRN1 | A2M |
| MMRN1 | CFD |
| MMRN1 | FIGF |
| MSR1 | COL1A1 |
| MUC4 | B3GNT3 |
| MUC4 | GALNT14 |
| NMU | GNG11 |
| NMUR1 | NMU |
| NMUR1 | GNG11 |
| NMUR1 | SSTR1 |
| NMUR1 | CXCL13 |
| NPNT | ITGA8 |
| NTNG1 | CEACAM5 |
| NTNG1 | LY6D |
| PCOLCE2 | COL1A1 |
| PCSK9 | VGF |
| PKMYT1 | CCNB1 |
| PLAC8 | FABP5 |
| PLAC8 | RETN |
| PTPRB | OLR1 |
| PTPRB | MCEMP1 |
| RBP4 | CES1 |
| RECK | NTNG1 |
| RECK | CEACAM5 |
| RECK | CD52 |
| RECK | LY6D |
| RECQL4 | CDC45 |
| RNF182 | UBE2C |
| SBSPON | ADAMTSL3 |

| | |
|---|---|
| SBSPON | ADAMTS8 |
| SCN7A | SCN4B |
| SEMA5A | SBSPON |
| SEMA5A | ADAMTS8 |
| SEMA5A | ADAMTSL3 |
| SFTPA1 | SFTPC |
| SFTPA1 | SFTPA2 |
| SFTPA1 | SFTPD |
| SFTPA2 | SFTPC |
| SFTPA2 | SFTPD |
| SFTPD | SFTPC |
| SH3GL3 | ADRB2 |
| SLIT2 | ROBO4 |
| SLIT2 | SLIT3 |
| SLIT3 | ROBO4 |
| SLPI | CAMP |
| SLPI | RETN |
| SPAG5 | CCNB2 |
| SPAG5 | BIRC5 |
| SPAG5 | TROAP |
| SPAG5 | KIAA0101 |
| SPAG5 | TPX2 |
| SPAG5 | CCNB1 |
| SPARCL1 | SPP1 |
| SPARCL1 | GPC3 |
| SPARCL1 | CHRDL1 |
| SPARCL1 | VGF |
| SPARCL1 | GOLM1 |
| SPARCL1 | PCSK9 |
| SPP1 | GPC3 |
| SPP1 | GOLM1 |
| SPP1 | ITGA8 |
| SPP1 | VGF |
| SPP1 | PCSK9 |
| SPP1 | CHRDL1 |
| SSTR1 | GNG11 |
| SSTR1 | NMU |
| STRA6 | RBP4 |
| TK1 | BIRC5 |
| TPSAB1 | MMP1 |
| TPX2 | CCNB1 |
| TPX2 | CCNB2 |

| | |
|---|---|
| TTN | TNNC1 |
| TTN | DES |
| UBE2C | CCNB1 |
| UBE2C | CCNB2 |
| UBE2C | BIRC5 |
| UBE2C | TPX2 |
| UBE2C | KIAA0101 |
| UBE2C | SPAG5 |
| UBE2T | UBE2C |
| UHRF1 | CDCA7 |
| VIPR1 | RAMP3 |
| VIPR1 | GNG11 |
| VIPR1 | ADRB2 |
| WIF1 | WNT3A |

**Supplementary Table S5:**

Node details of miRNA-mRNA regulatory network in LUAD

| node_name | Degree | Closeness | Betweenness |
|---|---|---|---|
| GPER1 | 6 | 121.46667 | 0 |
| TPSAB1 | 1 | 100.46667 | 0 |
| SBSPON | 3 | 113.4 | 0 |
| MCEMP1 | 3 | 120.3 | 0 |
| CD52 | 6 | 123.26667 | 0 |
| CLDN5 | 1 | 98.58333 | 0 |
| LEPREL1 | 3 | 119.9 | 0 |
| LY6D | 6 | 123.26667 | 0 |
| GNG11 | 15 | 131.06667 | 227.40329 |
| FIGF | 5 | 120.56667 | 25.94767 |
| SCGB3A2 | 1 | 87.51667 | 0 |
| CXCL13 | 6 | 121.46667 | 0 |
| KIAA0101 | 6 | 126.26667 | 0 |
| TNNC1 | 2 | 121.05 | 0 |
| HIST1H2AI | 1 | 1 | 0 |
| HIST2H3A | 1 | 1 | 0 |
| CDT1 | 3 | 119 | 0.51504 |
| NMU | 9 | 126.81667 | 41.67276 |
| HBA2 | 2 | 104.9 | 0 |
| RAMP3 | 5 | 123.21667 | 0 |
| UBE2C | 13 | 130.18333 | 62.33283 |
| CCNB2 | 15 | 131.35 | 109.13152 |

| | | | |
|---|---|---|---|
| MNX1 | 1 | 117.35 | 0 |
| VSIG4 | 1 | 122.43333 | 0 |
| RHPN1 | 1 | 122.68333 | 0 |
| CLDN18 | 3 | 138.7 | 740.13452 |
| SUSD2 | 1 | 119.38333 | 0 |
| WIF1 | 2 | 131.93333 | 36.40679 |
| IRX1 | 1 | 117.35 | 0 |
| SCGB1A1 | 1 | 116.01667 | 0 |
| PI16 | 3 | 140.78333 | 34.17422 |
| DCDC2B | 1 | 122.68333 | 0 |
| LGR4 | 2 | 127.7 | 124.47034 |
| FABP5 | 4 | 116 | 43.84731 |
| IHH | 3 | 117.63333 | 22.29164 |
| C2orf40 | 3 | 147.31667 | 76.43487 |
| CPNE7 | 3 | 147.31667 | 58.90694 |
| MMRN1 | 5 | 133.08333 | 182.17575 |
| FBLN5 | 3 | 124.51667 | 110.56014 |
| RSPO4 | 2 | 128.23333 | 126.7337 |
| SLC7A11 | 2 | 141.18333 | 25.45663 |
| VSIG2 | 2 | 131.31667 | 13.41517 |
| MMP9 | 10 | 159.03333 | 1203.06267 |
| LGI3 | 2 | 110.41667 | 0 |
| PRAM1 | 1 | 124.96667 | 0 |
| ALOX5 | 3 | 131.76667 | 121.04226 |
| HBB | 6 | 132.98333 | 347.17484 |
| TMEM190 | 1 | 127.26667 | 0 |
| ALOX5AP | 2 | 127.93333 | 0 |
| CEACAM5 | 9 | 161.66667 | 494.42977 |
| PEBP4 | 1 | 119.38333 | 0 |
| CA4 | 1 | 112.28333 | 0 |
| DNAH12 | 1 | 116.01667 | 0 |
| CDC45 | 7 | 159.16667 | 502.44188 |
| PLAC9 | 1 | 117.35 | 0 |
| ADRB2 | 10 | 148.5 | 405.46051 |
| DNAI2 | 5 | 146.73333 | 318.80816 |
| MS4A2 | 1 | 117.15 | 0 |
| MMP11 | 3 | 137.36667 | 179.12657 |
| CGNL1 | 5 | 156.33333 | 190.33727 |
| SRPX | 2 | 130.93333 | 12.85828 |
| SLC6A4 | 2 | 129.48333 | 8.34628 |
| VGF | 7 | 142.5 | 111.91285 |
| OGN | 3 | 138.1 | 77.01333 |

| | | | |
|---|---|---|---|
| VIPR1 | 9 | 161.36667 | 637.04314 |
| RECK | 9 | 153.41667 | 373.39247 |
| GPR152 | 1 | 122.43333 | 0 |
| SOSTDC1 | 1 | 117.15 | 0 |
| INMT | 2 | 138.4 | 16.71735 |
| FUT3 | 2 | 130.3 | 57.12636 |
| CLEC3B | 2 | 133.31667 | 62.75578 |
| C10orf67 | 2 | 140.15 | 15.93372 |
| TMEM100 | 4 | 153.25 | 107.60027 |
| CSH1 | 2 | 116.05 | 7.552 |
| CRABP2 | 2 | 140.48333 | 21.93886 |
| FRMD3 | 8 | 172.41667 | 560.44054 |
| FCN3 | 1 | 127.26667 | 0 |
| HEG1 | 6 | 167.03333 | 255.5161 |
| BCHE | 1 | 96.35 | 0 |
| CD19 | 4 | 151.7 | 83.81699 |
| MYOC | 2 | 141.51667 | 25.67183 |
| C9orf24 | 3 | 144.2 | 45.59714 |
| SLC39A8 | 3 | 126.65 | 13.5023 |
| MS4A15 | 1 | 94.96667 | 0 |
| ADAMTS8 | 6 | 144.33333 | 211.93689 |
| WNT3A | 6 | 144.41667 | 293.41613 |
| MMP1 | 5 | 141.28333 | 793.85525 |
| AQP4 | 2 | 140.15 | 15.93372 |
| HIGD1B | 3 | 148.35 | 57.17209 |
| KRT4 | 3 | 152.58333 | 63.58636 |
| CAV3 | 2 | 143.45 | 21.64943 |
| HMGB3 | 5 | 163.61667 | 177.02846 |
| CPAMD8 | 1 | 119.38333 | 0 |
| PCOLCE2 | 2 | 124.78333 | 0 |
| PHLDA2 | 1 | 124.96667 | 0 |
| BIRC5 | 15 | 158.91667 | 845.66985 |
| PXMP4 | 5 | 157.73333 | 174.42778 |
| F12 | 4 | 154.95 | 154.78327 |
| DNASE1L3 | 2 | 131.75 | 14.89729 |
| ADH1A | 5 | 155.33333 | 264.53842 |
| GLIPR2 | 2 | 129.85 | 18.92386 |
| TEK | 3 | 149.41667 | 26.16195 |
| F11 | 5 | 155.7 | 206.31827 |
| CA2 | 1 | 127.26667 | 0 |
| A2M | 11 | 159.33333 | 1253.32751 |
| RASAL1 | 3 | 149.01667 | 42.20664 |

| | | | |
|---|---|---|---|
| POU2AF1 | 3 | 151.36667 | 81.52859 |
| PTPRH | 1 | 111.96667 | 0 |
| FABP4 | 4 | 137.33333 | 190.5225 |
| EFCAB1 | 2 | 119.18333 | 13.03328 |
| GPC3 | 11 | 146.41667 | 410.62948 |
| ZBED2 | 3 | 149.16667 | 66.94055 |
| ITLN1 | 3 | 142.48333 | 109.32924 |
| SLIT3 | 6 | 161.28333 | 123.78391 |
| PID1 | 2 | 132.43333 | 7.91143 |
| RTKN2 | 5 | 164.28333 | 191.44069 |
| SFTPC | 5 | 146.91667 | 63.56198 |
| MFAP4 | 4 | 128.23333 | 121.27258 |
| CAV2 | 3 | 149.28333 | 38.9611 |
| TK1 | 3 | 121.63333 | 13.83233 |
| SCN4B | 3 | 132.11667 | 44.53771 |
| TMEM63C | 3 | 148.36667 | 58.03228 |
| FHL2 | 2 | 138.28333 | 22.37397 |
| B3GNT3 | 4 | 140.2 | 233.54379 |
| LGALSL | 2 | 130.93333 | 12.85828 |
| KCNN4 | 2 | 126.43333 | 77.82515 |
| ENTPD8 | 4 | 146.15 | 340.13624 |
| MGAT3 | 2 | 140.15 | 15.93372 |
| PGC | 1 | 122.43333 | 0 |
| FAM183A | 2 | 140.2 | 26.79079 |
| MAMDC2 | 4 | 160.5 | 123.30402 |
| GPX3 | 2 | 130.05 | 100.6293 |
| OLR1 | 5 | 151.5 | 158.47494 |
| CHRDL1 | 10 | 161.83333 | 523.02819 |
| CR2 | 6 | 165.53333 | 336.44149 |
| ABCA3 | 2 | 141.18333 | 25.45663 |
| MARCO | 2 | 117.81667 | 718 |
| CYP4B1 | 3 | 153.53333 | 52.63358 |
| MEX3A | 3 | 133.83333 | 86.79495 |
| NMUR1 | 10 | 145.08333 | 575.91057 |
| SFTPA1 | 6 | 153.33333 | 171.7711 |
| SFTPA2 | 9 | 171.66667 | 801.72729 |
| DES | 5 | 156.83333 | 183.05094 |
| CTHRC1 | 4 | 144.5 | 129.19539 |
| KIAA0408 | 5 | 152.03333 | 136.8145 |
| SYT7 | 3 | 145.28333 | 49.11555 |
| CLEC14A | 1 | 119.38333 | 0 |
| UHRF1 | 3 | 141.86667 | 131.02867 |

| | | | |
|---|---|---|---|
| PKMYT1 | 5 | 151.08333 | 113.09789 |
| S100P | 3 | 146.23333 | 25.45663 |
| EDN1 | 8 | 140.75 | 178.66654 |
| TIMP3 | 5 | 148.56667 | 263.60298 |
| C8B | 6 | 165.65 | 243.27355 |
| TNS4 | 2 | 135.9 | 15.72603 |
| PDZD2 | 5 | 151.86667 | 210.13401 |
| RAB26 | 2 | 141.51667 | 25.67183 |
| DCN | 6 | 157.33333 | 278.7489 |
| C14orf132 | 3 | 140.06667 | 70.48317 |
| TROAP | 6 | 151.2 | 135.0094 |
| CHIA | 5 | 150.41667 | 113.31787 |
| ACVRL1 | 4 | 153.75 | 71.50157 |
| EPHA10 | 7 | 161.61667 | 309.18802 |
| NLRC4 | 9 | 166.58333 | 648.16728 |
| SPARCL1 | 7 | 143.58333 | 28.83048 |
| TUBB3 | 3 | 148.36667 | 58.03228 |
| GKN2 | 2 | 123.68333 | 14.73247 |
| ANKRD29 | 4 | 156.73333 | 118.35181 |
| EMCN | 5 | 159.86667 | 195.87567 |
| CD36 | 11 | 172.41667 | 1144.19436 |
| RETN | 5 | 140.4 | 492.83026 |
| SLPI | 3 | 132.23333 | 118.94756 |
| hsa-miR-451b | 16 | 134.58333 | 969.80989 |
| ADRB1 | 7 | 144.61667 | 257.79806 |
| ABCA8 | 6 | 164.08333 | 271.67202 |
| MYZAP | 2 | 141.51667 | 25.67183 |
| CHRM1 | 11 | 166.25 | 1285.69902 |
| COL1A1 | 10 | 154.83333 | 839.97159 |
| GIMAP6 | 3 | 145.16667 | 49.42549 |
| EPN3 | 3 | 141.4 | 44.37888 |
| LIMCH1 | 5 | 160.66667 | 189.0019 |
| HJURP | 8 | 140.41667 | 100.13032 |
| PLAC8 | 4 | 136.23333 | 250.58198 |
| HLF | 3 | 152.11667 | 68.18113 |
| GPM6A | 6 | 162.83333 | 245.63968 |
| SYT12 | 3 | 144.15 | 55.13543 |
| DSP | 7 | 166.03333 | 257.50206 |
| GJB2 | 1 | 122.43333 | 0 |
| STRA6 | 6 | 160.95 | 244.49458 |
| RNF182 | 6 | 158.56667 | 242.23155 |
| CDH3 | 6 | 166.66667 | 300.99705 |

| | | | |
|---|---|---|---|
| GSTM5 | 3 | 147.81667 | 34.52736 |
| COL6A6 | 11 | 173 | 1306.8232 |
| SCN7A | 6 | 159.75 | 562.14856 |
| CENPF | 16 | 170 | 840.98782 |
| EPAS1 | 3 | 143.1 | 69.67113 |
| GPM6B | 6 | 168.25 | 314.3509 |
| NTNG1 | 11 | 166.58333 | 811.34151 |
| SELENBP1 | 4 | 139.41667 | 130.25034 |
| NDRG4 | 2 | 138.03333 | 22.13452 |
| CLIC3 | 1 | 122.68333 | 0 |
| TPPP3 | 4 | 153.08333 | 105.15875 |
| JAM2 | 3 | 147.31667 | 52.42309 |
| MMP12 | 4 | 154.41667 | 140.18419 |
| ITGA8 | 8 | 165.45 | 357.61559 |
| MME | 9 | 173.83333 | 604.88028 |
| CX3CR1 | 10 | 165.53333 | 1486.53588 |
| LAMP3 | 3 | 144.76667 | 55.42873 |
| CDCA5 | 6 | 143.66667 | 25.31988 |
| DLC1 | 8 | 167 | 484.10647 |
| ITLN2 | 1 | 94.96667 | 0 |
| CAV1 | 10 | 174.83333 | 1080.94029 |
| PSAT1 | 3 | 144.81667 | 55.08667 |
| ADAMTSL3 | 6 | 144.11667 | 299.21722 |
| FAM83A | 4 | 155.2 | 116.47085 |
| ABI3BP | 5 | 144.75 | 132.34177 |
| TNS1 | 7 | 170.75 | 417.02199 |
| PCSK9 | 11 | 170.25 | 619.62753 |
| PTPN21 | 6 | 160.61667 | 192.97137 |
| LDLRAD1 | 3 | 151.65 | 73.57321 |
| FGL1 | 4 | 146.4 | 106.97971 |
| DYNLRB2 | 2 | 122.4 | 0 |
| CLIC5 | 4 | 160.5 | 123.30402 |
| VGLL3 | 2 | 125.18333 | 17.89475 |
| CDH13 | 5 | 159.91667 | 196.60685 |
| CDO1 | 3 | 136.7 | 46.35355 |
| GOLM1 | 10 | 159 | 373.20293 |
| DENND3 | 6 | 169.33333 | 333.06251 |
| VWF | 7 | 145.33333 | 383.4447 |
| AGER | 6 | 159.31667 | 278.6465 |
| CPA3 | 1 | 116.01667 | 0 |
| FRAS1 | 10 | 174.16667 | 756.35909 |
| KIF2C | 14 | 157.33333 | 269.45607 |

| | | | |
|---|---|---|---|
| PITX1 | 3 | 148.31667 | 60.64262 |
| LPL | 4 | 138.36667 | 138.20335 |
| ABCA12 | 8 | 171.66667 | 751.42594 |
| GIMAP8 | 2 | 135.28333 | 16.45594 |
| SPINK1 | 1 | 127.26667 | 0 |
| ROBO4 | 7 | 163.03333 | 257.80373 |
| CARD14 | 4 | 155.65 | 82.11616 |
| hsa-miR-126-5p | 8 | 127.66667 | 35.27159 |
| PYCR1 | 3 | 152.31667 | 82.03948 |
| BTNL9 | 4 | 155.45 | 122.61938 |
| C11orf88 | 1 | 122.43333 | 0 |
| ADH1C | 4 | 145.25 | 225.09934 |
| NDNF | 1 | 117.15 | 0 |
| PDE5A | 3 | 150.31667 | 71.18528 |
| SLC1A1 | 4 | 146.58333 | 68.97325 |
| SPOCK2 | 8 | 167.36667 | 582.64778 |
| SCARA5 | 3 | 150.31667 | 71.18528 |
| HBD | 5 | 139.31667 | 501.02049 |
| VEPH1 | 5 | 152.98333 | 120.424 |
| MSR1 | 4 | 151.75 | 117.02201 |
| CRYAB | 3 | 146.56667 | 49.59729 |
| IQGAP3 | 4 | 144.98333 | 121.49484 |
| RSPH1 | 3 | 149.16667 | 66.94055 |
| THBD | 1 | 122.43333 | 0 |
| MUC4 | 8 | 169.45 | 559.26137 |
| ACOXL | 3 | 141.4 | 44.37888 |
| EMP2 | 4 | 147.48333 | 100.00154 |
| CACNG6 | 3 | 149.36667 | 36.37701 |
| AOC3 | 7 | 164.95 | 389.77552 |
| FMO2 | 2 | 129.85 | 18.92386 |
| CALCRL | 11 | 165.03333 | 1021.28729 |
| SGPP2 | 4 | 151.06667 | 188.40084 |
| PTPRB | 13 | 174.5 | 1224.06834 |
| hsa-miR-144-5p | 7 | 128.33333 | 92.84573 |
| SLC19A3 | 4 | 142 | 67.28264 |
| LYVE1 | 2 | 134.4 | 19.44116 |
| AQP1 | 6 | 161.75 | 266.76558 |
| CDHR4 | 5 | 164.86667 | 215.83164 |
| CYP2F1 | 2 | 135.28333 | 16.45594 |
| LRRC36 | 5 | 154.7 | 170.04146 |
| LGSN | 7 | 167.61667 | 369.65501 |
| GINS1 | 4 | 150.81667 | 97.2574 |

| | | | |
|---|---|---|---|
| FXYD1 | 2 | 138.31667 | 23.30942 |
| CFD | 5 | 134.41667 | 250.39777 |
| SLIT2 | 10 | 174.78333 | 686.44917 |
| ADH1B | 4 | 148.08333 | 152.39171 |
| LRRN3 | 2 | 143.45 | 21.64943 |
| CEP55 | 12 | 157.58333 | 264.35836 |
| ARHGEF26 | 3 | 147.28333 | 61.60014 |
| CAMP | 7 | 137.4 | 296.07919 |
| SPP1 | 11 | 159.66667 | 627.249 |
| hsa-miR-21-5p | 23 | 144 | 694.77126 |
| CES1 | 5 | 156.78333 | 193.58328 |
| CCDC85A | 1 | 124.96667 | 0 |
| DUOXA1 | 7 | 170.53333 | 342.29135 |
| RECQL4 | 5 | 157.16667 | 248.29139 |
| LRRK2 | 3 | 149.41667 | 87.67515 |
| FAM107A | 6 | 165.4 | 223.30993 |
| METTL7A | 7 | 159.73333 | 542.02157 |
| AFF3 | 5 | 163.91667 | 206.2065 |
| ANXA3 | 3 | 149.66667 | 72.12447 |
| NPNT | 8 | 167.53333 | 387.95201 |
| STXBP6 | 4 | 156.65 | 98.23747 |
| CRTAC1 | 3 | 148.1 | 55.34004 |
| SVEP1 | 8 | 165.7 | 444.09062 |
| TMPRSS4 | 8 | 164.45 | 621.76487 |
| NCKAP5 | 8 | 170.66667 | 465.73499 |
| MRC1 | 5 | 154.25 | 197.49989 |
| GPRIN2 | 6 | 160 | 216.05751 |
| GRK5 | 8 | 140.58333 | 184.26708 |
| GPIHBP1 | 9 | 151.83333 | 416.55297 |
| LMO2 | 2 | 126.88333 | 31.22987 |
| hsa-miR-144-3p | 9 | 131.33333 | 403.58538 |
| UBE2T | 3 | 144.35 | 55.15485 |
| ANGPT1 | 10 | 176.41667 | 892.35035 |
| COL10A1 | 4 | 142.75 | 190.22047 |
| GALNT14 | 9 | 165.7 | 507.12757 |
| HHIP | 4 | 148.15 | 393.48239 |
| ASPM | 20 | 174 | 1409.60048 |
| hsa-miR-451a | 16 | 137.31667 | 832.49219 |
| GLDN | 4 | 156.33333 | 134.65671 |
| STX1A | 6 | 152.36667 | 524.23793 |
| HS6ST2 | 4 | 149.08333 | 112.16845 |
| ERBB4 | 6 | 158.95 | 243.00244 |

| | | | |
|---|---|---|---|
| GPA33 | 2 | 143.45 | 21.64943 |
| FAM189A2 | 5 | 157.83333 | 167.29424 |
| SH3GL3 | 4 | 149.73333 | 139.15196 |
| WISP2 | 2 | 123.15 | 5.41036 |
| hsa-miR-96-5p | 61 | 173.16667 | 6224.055 |
| C1orf21 | 7 | 164.33333 | 302.73213 |
| SEMA5A | 8 | 159.66667 | 841.90514 |
| hsa-miR-135b-5p | 53 | 167.91667 | 4186.44533 |
| TTN | 17 | 183 | 2561.85054 |
| CACNA2D2 | 10 | 172.53333 | 1046.94087 |
| CYBRD1 | 5 | 155.45 | 234.07007 |
| hsa-miR-126-3p | 15 | 131.83333 | 1692.90521 |
| CABYR | 6 | 168.25 | 287.39956 |
| XDH | 5 | 163.81667 | 198.35236 |
| FERMT1 | 1 | 122.43333 | 0 |
| RBP4 | 4 | 150.2 | 148.41248 |
| PROM2 | 4 | 155.91667 | 106.19626 |
| SPAG5 | 15 | 156.58333 | 421.40008 |
| hsa-miR-218-5p | 19 | 139.16667 | 682.95505 |
| GDF10 | 3 | 140.33333 | 37.83438 |
| hsa-miR-338-3p | 65 | 174.5 | 6627.11746 |
| TCF21 | 3 | 137.7 | 57.82919 |
| hsa-miR-30a-5p | 15 | 131.81667 | 490.15826 |
| SSTR1 | 9 | 147.7 | 589.49893 |
| MS4A7 | 3 | 139.91667 | 75.94884 |
| CCL14 | 1 | 122.43333 | 0 |
| TNXB | 3 | 151.65 | 73.57321 |
| FCRL5 | 4 | 154.91667 | 113.44533 |
| LIMS2 | 3 | 149.41667 | 87.67515 |
| C1orf116 | 4 | 154.61667 | 120.63351 |
| hsa-miR-218-1-3p | 77 | 183.91667 | 10779.92394 |
| CCNB1 | 19 | 176.91667 | 2015.24643 |
| hsa-miR-486-5p | 119 | 213.41667 | 22792.81553 |
| LYPD1 | 10 | 167.83333 | 877.30481 |
| hsa-miR-183-5p | 97 | 200.08333 | 18239.90403 |
| TPX2 | 14 | 173.75 | 641.69738 |
| hsa-miR-30a-3p | 83 | 187.16667 | 12694.00847 |
| DUOX1 | 6 | 167.78333 | 270.8367 |
| CDCA7 | 3 | 146.45 | 185.36751 |
| hsa-miR-139-5p | 85 | 192 | 13732.94821 |
| SFTPD | 6 | 149.66667 | 238.10598 |
| HSD17B6 | 5 | 158.5 | 172.71298 |

| | | | |
|---|---|---|---|
| hsa-miR-218-2-3p | 79 | 186.91667 | 11190.22554 |
| ACADL | 6 | 160.03333 | 198.91399 |
| hsa-miR-200b-5p | 110 | 207.33333 | 19421.27697 |
| PDK4 | 3 | 148.95 | 49.08698 |
| hsa-miR-130b-5p | 102 | 201.41667 | 16604.76576 |

**Supplementary Table S6:**

Proteins enriched in the extracellular matrix organization pathway

| Gene Symbol | Uniprot ID | Ensemble ID |
|---|---|---|
| ACVRL1 | P37023 | ENSG00000139567 |
| NDRG4 | Q9ULP0 | ENSG00000103034 |
| TNXB | P22105 | ENSG00000168477 |
| PCOLCE2 | Q9UKZ9 | ENSG00000163710 |
| SLC2A1 | P11166 | ENSG00000227533 |
| FBLN5 | Q9UBX5 | ENSG00000140092 |
| HS6ST2 | Q96MM7 | ENSG00000171004 |
| SPP1 | P10451, Q9BX95 | ENSG00000118785 |
| COL10A1 | Q03692 | ENSG00000123500 |
| TPSAB1 | Q15661 | ENSG00000172236 |
| A2M | P01023 | ENSG00000175899 |
| ADAMTS8 | Q9UP79 | ENSG00000134917 |
| JAM2 | P57087 | ENSG00000154721 |
| GPM6B | Q13491 | ENSG00000046653 |
| MME | P08473 | ENSG00000196549 |
| VWF | P04275 | ENSG00000110799 |
| MMP1 | P03956 | ENSG00000196611 |
| F11 | P03951 | ENSG00000088926 |
| CACNA2D2 | Q9NY47 | ENSG00000007402 |
| LEPREL1 | Q8IVL5 | ENSG00000090530 |
| MMP9 | P14780 | ENSG00000100985 |
| DCN | P07585 | ENSG00000011465 |
| MMP12 | P39900 | ENSG00000262406 |
| COL1A1 | P02452 | ENSG00000108821 |
| MMP11 | P24347 | ENSG00000099953 |
| MFAP4 | P55083 | ENSG00000166482 |
| ITGA8 | P53708 | ENSG00000077943 |
| COL6A6 | A6NMZ7 | ENSG00000206384 |

**Supplementary Table S7:**

Tendency of cooccurrence of the hub genes in LUAD

| A | B | Neither | A Not B | B Not A | Both | Log2 Odds Ratio | p-Value | q-Value | Tendency |
|---|---|---|---|---|---|---|---|---|---|
| ASPM | CENPF | 922 | 142 | 65 | 60 | 2.583 | <0.001 | <0.001 | Co-occurrence |
| ASPM | TTN | 565 | 69 | 422 | 133 | 1.368 | <0.001 | <0.001 | Co-occurrence |
| TTN | SPAG5 | 630 | 531 | 4 | 24 | 2.832 | <0.001 | <0.001 | Co-occurrence |
| SPAG5 | TPX2 | 1119 | 21 | 42 | 7 | >3 | <0.001 | <0.001 | Co-occurrence |
| CENPF | CCNB2 | 1059 | 119 | 5 | 6 | >3 | <0.001 | 0.003 | Co-occurrence |
| TTN | CENPF | 585 | 479 | 49 | 76 | 0.922 | <0.001 | 0.004 | Co-occurrence |
| GNG11 | KIF2C | 1146 | 20 | 19 | 4 | >3 | <0.001 | 0.006 | Co-occurrence |
| TTN | KIF2C | 629 | 537 | 5 | 18 | 2.076 | 0.002 | 0.011 | Co-occurrence |
| TTN | TPX2 | 618 | 522 | 16 | 33 | 1.288 | 0.002 | 0.012 | Co-occurrence |
| CCNB1 | KIF2C | 1151 | 15 | 20 | 3 | >3 | 0.004 | 0.019 | Co-occurrence |
| CCNB1 | GNG11 | 1150 | 15 | 21 | 3 | >3 | 0.005 | 0.02 | Co-occurrence |
| CCNB2 | TPX2 | 1132 | 8 | 46 | 3 | >3 | 0.009 | 0.032 | Co-occurrence |
| ASPM | SPAG5 | 969 | 192 | 18 | 10 | 1.487 | 0.012 | 0.043 | Co-occurrence |
| SPAG5 | KIF2C | 1141 | 25 | 20 | 3 | 2.775 | 0.015 | 0.049 | Co-occurrence |
| GNG11 | SPAG5 | 1140 | 21 | 25 | 3 | 2.704 | 0.017 | 0.051 | Co-occurrence |
| ASPM | KIF2C | 972 | 194 | 15 | 8 | 1.418 | 0.029 | 0.082 | Co-occurrence |
| TTN | GNG11 | 626 | 539 | 8 | 16 | 1.216 | 0.038 | 0.099 | Co-occurrence |
| ASPM | CCNB2 | 980 | 198 | 7 | 4 | 1.5 | 0.1 | 0.25 | Co-occurrence |
| ASPM | TPX2 | 950 | 190 | 37 | 12 | 0.697 | 0.111 | 0.264 | Co-occurrence |
| CENPF | BIRC5 | 1034 | 124 | 30 | 1 | -1.847 | 0.145 | 0.326 | Mutual exclusivity |
| ASPM | CCNB1 | 970 | 201 | 17 | 1 | -1.817 | 0.162 | 0.335 | Mutual exclusivity |
| BIRC5 | SPAG5 | 1132 | 29 | 26 | 2 | 1.586 | 0.164 | 0.335 | Co-occurrence |
| GNG11 | CCNB2 | 1155 | 23 | 10 | 1 | 2.328 | 0.202 | 0.395 | Co-occurrence |

| | | | | | | | | | |
|---|---|---|---|---|---|---|---|---|---|
| TTN | BIR C5 | 620 | 538 | 14 | 17 | 0.485 | 0.229 | 0.43 | Co-occurrence |
| ASP M | BIR C5 | 963 | 195 | 24 | 7 | 0.526 | 0.264 | 0.476 | Co-occurrence |
| CCN B1 | TTN | 626 | 8 | 545 | 10 | 0.522 | 0.3 | 0.519 | Co-occurrence |
| GNG 11 | TPX 2 | 1116 | 24 | 49 | 0 | <-3 | 0.361 | 0.601 | Mutual exclusivity |
| ASP M | GNG 11 | 966 | 199 | 21 | 3 | -0.528 | 0.397 | 0.638 | Mutual exclusivity |
| CEN PF | TPX 2 | 1021 | 119 | 43 | 6 | 0.26 | 0.412 | 0.639 | Co-occurrence |
| CEN PF | KIF2 C | 1044 | 122 | 20 | 3 | 0.36 | 0.442 | 0.663 | Co-occurrence |
| BIR C5 | KIF2 C | 1136 | 30 | 22 | 1 | 0.783 | 0.458 | 0.666 | Co-occurrence |
| GNG 11 | BIR C5 | 1134 | 24 | 31 | 0 | <-3 | 0.527 | 0.706 | Mutual exclusivity |
| CEN PF | GNG 11 | 1042 | 123 | 22 | 2 | -0.377 | 0.529 | 0.706 | Mutual exclusivity |
| CCN B1 | TPX 2 | 1123 | 17 | 48 | 1 | 0.461 | 0.534 | 0.706 | Co-occurrence |
| CEN PF | SPA G5 | 1039 | 122 | 25 | 3 | 0.031 | 0.578 | 0.711 | Co-occurrence |
| CCN B1 | CEN PF | 1048 | 16 | 123 | 2 | 0.091 | 0.58 | 0.711 | Co-occurrence |
| TTN | CCN B2 | 628 | 550 | 6 | 5 | -0.072 | 0.59 | 0.711 | Mutual exclusivity |
| CCN B1 | BIR C5 | 1140 | 18 | 31 | 0 | <-3 | 0.619 | 0.711 | Mutual exclusivity |
| KIF2 C | TPX 2 | 1118 | 22 | 48 | 1 | 0.082 | 0.624 | 0.711 | Co-occurrence |
| BIR C5 | TPX 2 | 1110 | 30 | 48 | 1 | -0.376 | 0.632 | 0.711 | Mutual exclusivity |
| CCN B1 | SPA G5 | 1143 | 18 | 28 | 0 | <-3 | 0.649 | 0.712 | Mutual exclusivity |
| CCN B2 | BIR C5 | 1147 | 11 | 31 | 0 | <-3 | 0.747 | 0.8 | Mutual exclusivity |
| CCN B2 | SPA G5 | 1150 | 11 | 28 | 0 | <-3 | 0.769 | 0.804 | Mutual exclusivity |
| CCN B2 | KIF2 C | 1155 | 11 | 23 | 0 | <-3 | 0.806 | 0.824 | Mutual exclusivity |
| CCN B1 | CCN B2 | 1160 | 18 | 11 | 0 | <-3 | 0.845 | 0.845 | Mutual exclusivity |
